# Early acquisition of S-specific Tfh clonotypes after SARS-CoV-2 vaccination is associated with the longevity of anti-S antibodies

**DOI:** 10.1101/2023.06.06.543529

**Authors:** Xiuyuan Lu, Hiroki Hayashi, Eri Ishikawa, Yukiko Takeuchi, Julian Vincent Tabora Dychiao, Hironori Nakagami, Sho Yamasaki

## Abstract

SARS-CoV-2 vaccines have been used worldwide to combat COVID-19 pandemic. To elucidate the factors that determine the longevity of spike (S)-specific antibodies, we traced the characteristics of S-specific T cell clonotypes together with their epitopes and anti-S antibody titers before and after BNT162b2 vaccination over time. T cell receptor (TCR) αβ sequences and mRNA expression of the S-responded T cells were investigated using single-cell TCR- and RNA-sequencing. Highly expanded 199 TCR clonotypes upon stimulation with S peptide pools were reconstituted into a reporter T cell line for the determination of epitopes and restricting HLAs. Among them, we could determine 78 S epitopes, most of which were conserved in variants of concern (VOCs). After the 2nd vaccination, T cell clonotypes highly responsive to recall S stimulation were polarized to follicular helper T (Tfh)-like cells in donors exhibiting sustained anti-S antibody titers (designated as “sustainers”), but not in “decliners”. Even before vaccination, S-reactive CD4^+^ T cell clonotypes did exist, most of which cross-reacted with environmental or symbiotic bacteria. However, these clonotypes contracted after vaccination. Conversely, S-reactive clonotypes dominated after vaccination were undetectable in pre-vaccinated T cell pool, suggesting that highly-responding S-reactive T cells were established by vaccination from rare clonotypes. These results suggest that *de novo* acquisition of memory Tfh-like cells upon vaccination may contribute to the longevity of anti-S antibody titers.

## Introduction

The pandemic COVID-19, caused by the severe acute respiratory syndrome coronavirus 2 (SARS-CoV-2), has expanded worldwide [1]. Many types of vaccines have been developed or in basic and clinical phases to combat infection and deterioration of COVID-19 [2,3]. Among them, messenger ribonucleic acid (mRNA) vaccines, BNT162b2/Comirnaty and mRNA-1273/Spikevax, have been approved with over 90% efficacy at 2 months post-2nd dose vaccination [4,5], and widely used. Pathogen-specific antibodies are one of the most efficient components to prevent infection. Yet, mRNA vaccine-induced serum antibody titer is known to be waning over 6 months [6,7]. Accordingly, the effectiveness of the vaccines decreases over time, and thus multiple doses and repeated boosters are necessary [8].

The production and sustainability of spike (S)-specific antibody could be related to multiple factors, especially in the case of humans [7,9]. Among them, the characteristics of SARS-CoV-2-specific T cells are critically involved in the affinity and longevity of the antibodies [10–12]. Elucidation of the key factors of T cell responses that contribute to the durable immune responses induced by vaccination would provide valuable information for the vaccine development in the future. However, the relationship between antibody sustainability and the types of antigen-specific T cells has not been investigated in a clonotype resolution.

Recent studies reported that S-reactive T cells pre-existed before exposure to SARS-CoV-2 [13–17]. Common cold human coronaviruses (HCoVs) including strains 229E, NL63, OC43, and HKU1 are considered major cross-reactive antigens that primed these pre-existing T cells [15,18–20], while bacterial cross-reactive antigens were also reported [21,22]. However, the functional relevance of cross-reactive T cells during infection or vaccination is still in debate.

In this study, both humoral and cellular immune responses were evaluated at 3, 6 and 24 weeks after BNT162b2/Comirnaty vaccination. S-specific T cells before and after vaccination were analyzed on clonotype level using single cell-based T cell receptor (TCR) and RNA sequencing to determine their characteristics and epitopes in antibody sustainers and decliners. These analyses suggest the importance of early acquisition of S-specific Tfh cells in the longevity of antibodies.

## Results

### SARS-CoV-2 mRNA vaccine elicits transient humoral immunity

Blood samples were collected from a total of 43 individuals (Table 1) who had no SARS-CoV-2 infection history when they received two doses of SARS-CoV-2 mRNA vaccine BNT162b2. Samples were taken before and after the vaccination (Fig. 1A). Consistent with the previous report [4], most participants exhibited more severe side effects after 2nd dose of vaccination than 1st dose locally (Table 2) and systemically (Table 3). At 3 weeks, anti-S IgG antibody titer increased in most participants. At 6 weeks, anti-S antibody titer was at its peak. S antibody titer gradually decreased over 24 weeks (Fig. 1B). The antibody titer was reduced by 56.8% on average. Donors of different genders or age groups showed no significant difference in anti-S antibody titer (Fig. S1). The neutralization activity of the post-vaccinated sera showed similar tendency with the anti-S antibody titer during the study period (Fig. 1C). The above results indicate that the mRNA vaccine effectively activated humoral immune responses in healthy individuals, but decreased by 24 weeks over time as reported [6,7].

**Fig. 1.**
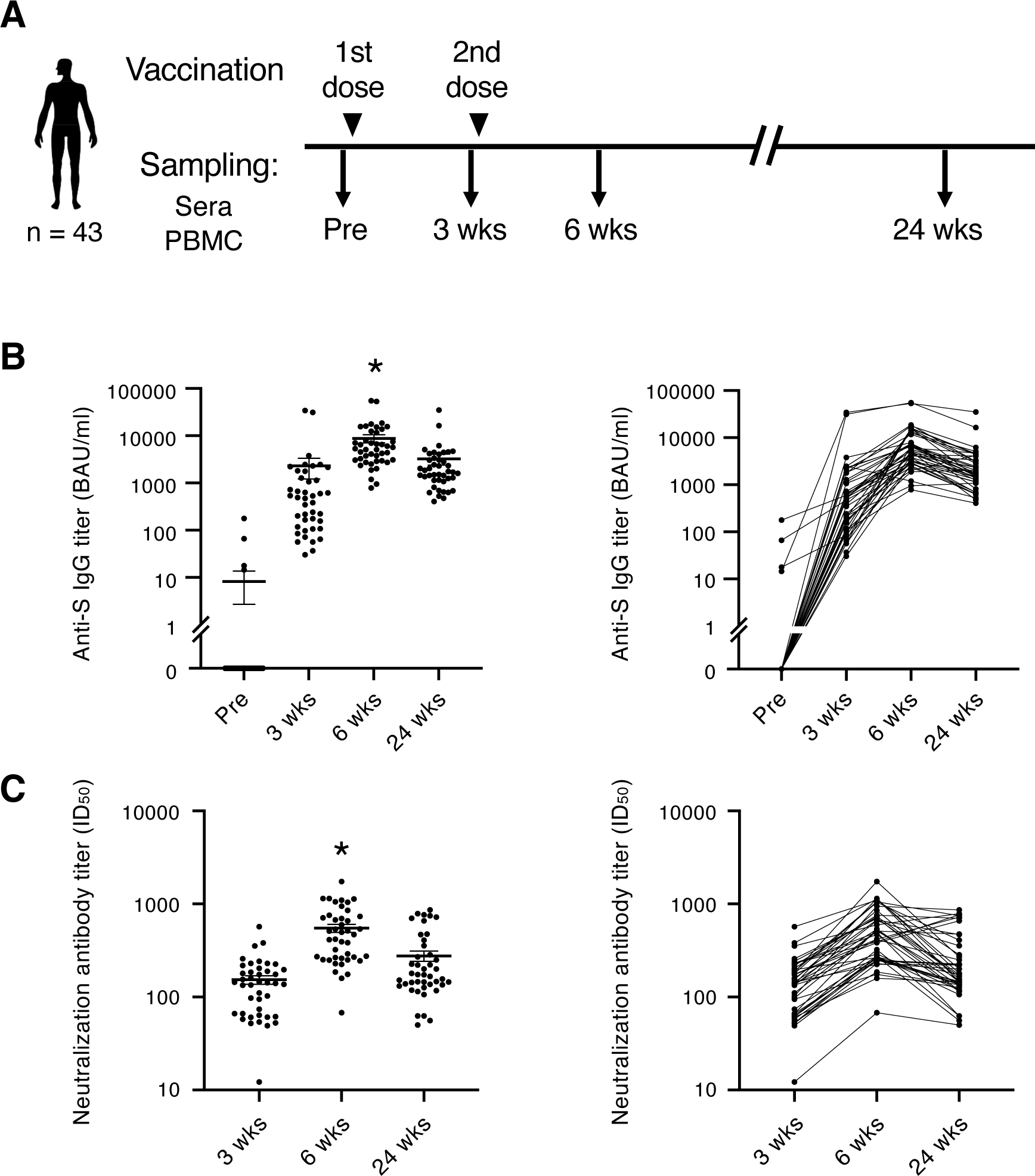
SARS-CoV-2 mRNA vaccine elicits transient humoral immunity. (**A**) Vaccination and sampling timeline of blood donors in this study. (**B**) Anti-S IgG titer of serum samples was determined by ELISA. Mean ± SEM (left) and individual data (right) are shown. *, *P* < 0.05 vs. Pre, 3 wks, 24 wks, respectively. (**C**) Neutralization activity (ID50) of serum samples was determined by pseudo-virus assay. Mean ± SEM (left) and individual data (right) are shown. *, *P* < 0.05 vs. 3 wks, 24 wks, respectively. Wks, weeks.

**Table 1.**
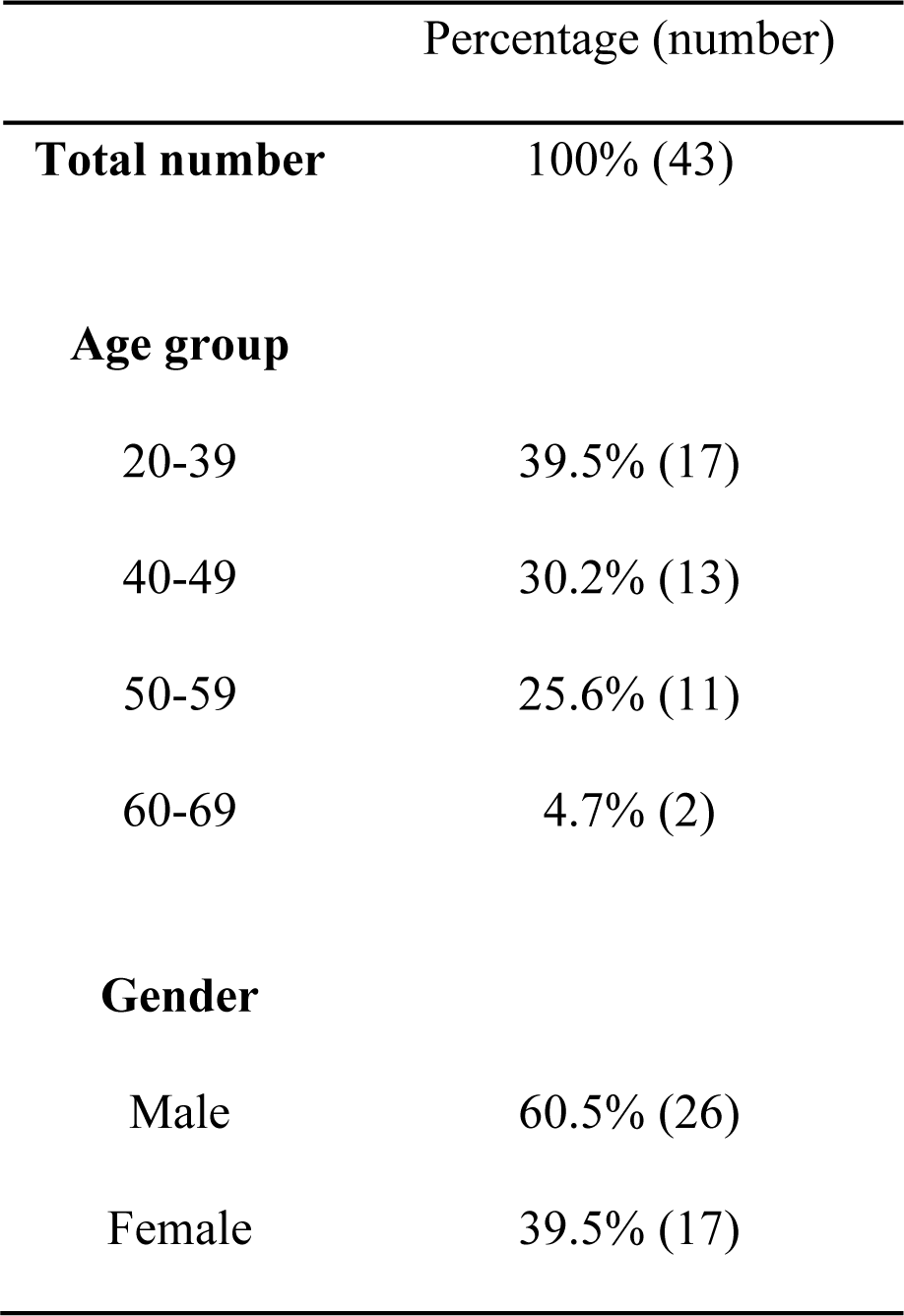
Demographic data of the participants.

**Table 2.** Demographic data of the reported clinical adverse effects (at injection site)

|  | Percentage (number) |
| --- | --- |
| **Swelling (injection site)** |  |
| After 1st dose | 27.9% (12) |
| After 2nd dose | 51.2% (22) |
| **Sore/pain (injection site)** |  |
| After 1st dose | 88.4% (38) |
| After 2nd dose | 86.0% (37) |
| **Warmth (injection site)** |  |
| After 1st dose | 32.6% (14) |
| After 2nd dose | 41.9% (18) |

**Table 3.** Demographic data of the reported clinical adverse effects (systemic symptoms)

|  | Percentage (number) |
| --- | --- |
| **Fever** |  |
| After 1st dose |  |
| Mild ($37.5^{\circ}\text{C} \geq$) | 2.3% (1) |
| Severe ($\geq 38.0^{\circ}\text{C}$) | 0% (0) |
| After 2nd dose |  |
| Mild ($37.5^{\circ}\text{C} \geq$) | 25.6% (11) |
| Severe ($\geq 38.0^{\circ}\text{C}$) | 23.3% (10) |
| **Fatigue** |  |
| After 1st dose |  |
| Mild | 18.6% (8) |
| Severe | 0% (0) |
| After 2nd dose |  |
| Mild | 67.4% (29) |
| Severe | 18.6% (8) |
| **Headache** |  |
| After 1st dose |  |
| Mild | 7.0% (3) |
| Severe | 0% (0) |
| After 2nd dose |  |
| Mild | 32.6% (14) |
| Severe | 7.0% (3) |
| **Chill** |  |
| After 1st dose |  |
| Mild | 4.7% (2) |
| Severe | 0% (0) |
| After 2nd dose |  |
| Mild | 23.3% (10) |
| Severe | 9.3% (4) |
| **Nausea** |  |
| After 1st dose |  |
| Mild | 0% (0) |
| Severe | 0% (0) |
| After 2nd dose |  |
| Mild | 4.7% (2) |
| Severe | 0% (0) |
| **Diarrhea** |  |
| After 1st dose |  |
| Mild | 0% (0) |
| Severe | 0% (0) |
| After 2nd dose |  |
| Mild | 0% (0) |
| Severe | 0% (0) |
| **Muscle pain** |  |
| After 1st dose |  |
| Mild | 48.8% (21) |
| Severe | 0% (0) |
| After 2nd dose |  |
| Mild | 55.8% (24) |
| Severe | 4.7% (2) |
| **Joint pain** |  |
| After 1st dose |  |
| Mild | 4.7% (2) |
| Severe | 0% (0) |
| After 2nd dose |  |
| Mild | 25.6% (11) |
| Severe | 4.7% (2) |

### Antibody sustainers had highly expanded S-reactive Tfh clonotypes

To address the role of T cells in maintaining the antibody titer, we analyzed the S-responsive T cells in the post-vaccination samples from 8 donors, among whom 4 donors showed relatively sustained anti-S antibody titer during 6 weeks to 24 weeks (reduction < 30%) (sustainers, donors #8, #25, #27 and #28), while the other 4 donors showed largely declined anti-S antibody titer (reduction > 80%) (decliners, donors #4, #13, #15 and #17) (Fig. 2A and Fig. S2A). The possibility of SARS-CoV-2 infection of sustainers was ruled out by analyzing anti-nucleocapsid protein (N) antibody titer in the sera samples at 24 weeks (Fig. S2B). Antibody sustainability did not correlate with bulk T cell responses to S protein, such as IFNγ production (Fig. S2C).

**Fig. 2.**
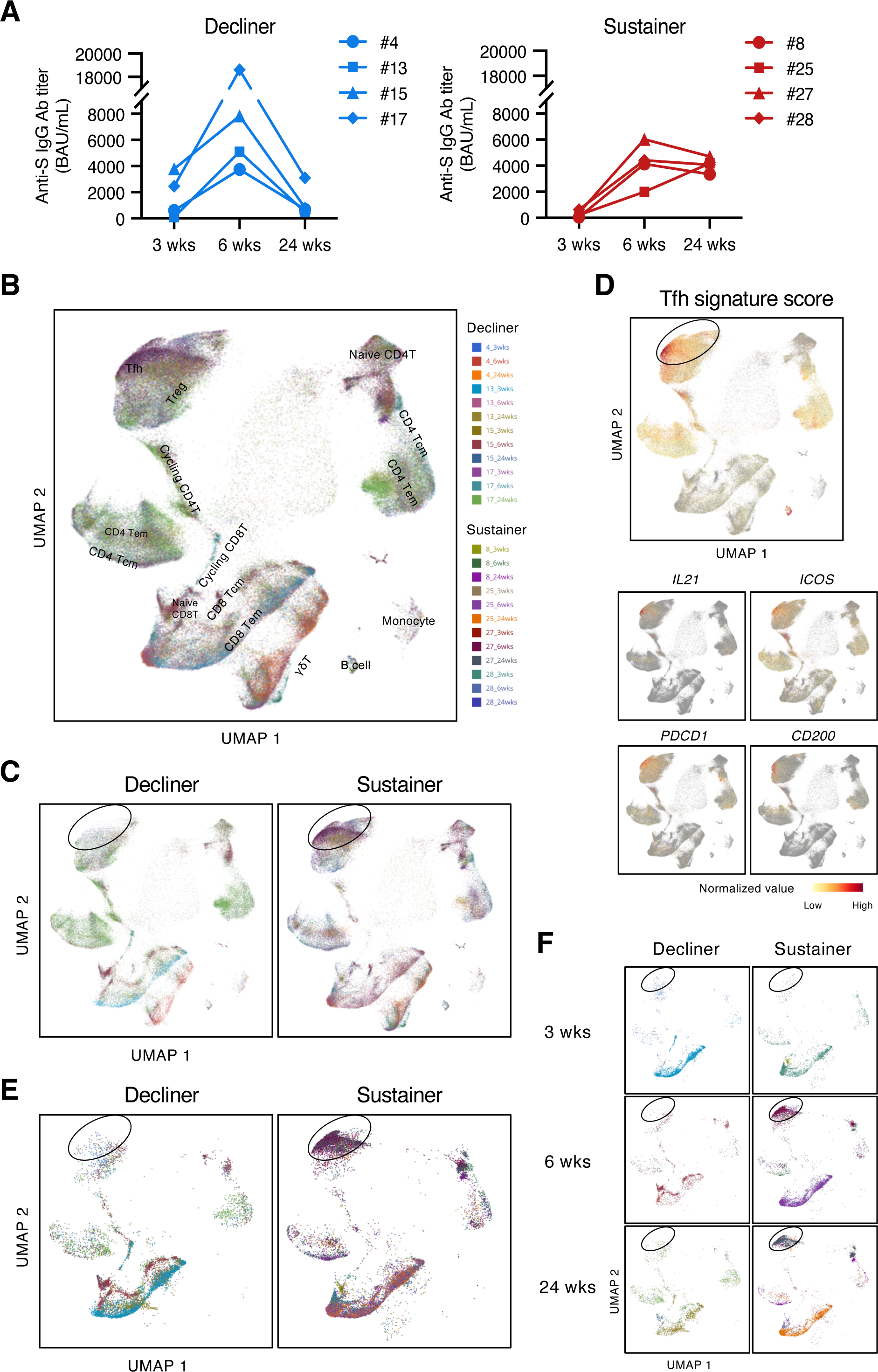
Antibody sustainers had highly expanded S-reactive Tfh clonotypes. (**A**) Anti-S IgG titer of serum samples from sustainers and decliners is shown individually. (**B, C, E,** and **F**) UMAP projection of T cells in single-cell analysis of post-vaccinated samples collected from all donors. Each dot corresponds to a single cell and is colored according to the samples from different time points of donors. All samples together with annotated cell types (B), samples grouped by donor type (decliners and sustainers) (C), top 16 expanded clonotypes (16 clonotypes that had the most cell numbers from each donor) grouped by donor type (E), and top 16 expanded clonotypes grouped by time point and donor type (F) are shown. Tcm, central memory T cells; Tem, effector memory T cells; Treg, regulatory T cells; γδT, γδ T cells. (**D**) Tfh signature score and expression levels of the canonical Tfh cell markers, *IL21*, *ICOS*, *PDCD1* and *CD200*, are shown as heat maps in the UMAP plot.

To enrich the S-reactive T cells, we labeled the peripheral blood mononuclear cells (PBMCs) with a cell proliferation tracer and stimulated the PBMCs with an S peptide pool for 10 days. Proliferated T cells were sorted and analyzed by single-cell TCR- and RNA-sequencing (scTCR/RNA-seq). Clustering analysis was done with pooled samples of 3 time points from 8 donors, and various T cell subtypes were identified (Fig. 2B). We found that, overall, the S-reactive T cells did not skew to any particular T cell subset (Fig. 2B). However, by grouping the cells from decliners and sustainers separately, we found difference in the frequency of the cells within the circled population (Fig. 2C), and overall, the sustainer individuals had more cells in this region (Fig. S3). These cells showed high Tfh signature scores and expressed characteristic genes of Tfh cells (Fig. 2D). This tendency became more pronounced when we selected highly expanded (top 16) clonotypes in each donor (Fig. 2E). In sustainers, S-specific Tfh clusters appeared from 6 weeks (Fig. 2F), suggesting that vaccine-induced Tfh-like cells that have potency of deriving to Tfh cells were established immediately after 2nd vaccination.

### Identification of dominant S epitopes recognized by vaccine-induced T cell clonotypes

To elucidate the epitopes of the highly expanded clonotypes, we reconstituted their TCRs into a T cell hybridoma lacking endogenous TCRs and having an NFAT-GFP reporter gene. These cell lines were stimulated with S peptides using transformed autologous B cells as antigen-presenting cells (APCs). The epitopes of 53 out of 128 reconstituted clonotypes were successfully determined (Fig. 3, Table 4, Figs. S4A–S4D). Epitopes of expanded Tfh cells were not limited in any particular region of S protein (Fig. 3). About 72% of these epitopes conserved in Delta and Omicron variants (Tables 4 and 5). Within the rest of 28% of epitopes which were mutated in variants of concern (VOCs), although some mutated epitopes located in the receptor-binding domain (RBD) of VOCs lost antigenicity, recognition of most epitopes outside the RBD region was maintained or rather increased in the variants (Table 5 and Figs. S4E and S4F). These results suggest that the majority of S-reactive clonotypes after vaccination can respond to antibody-escaping VOCs.

**Fig. 3.**
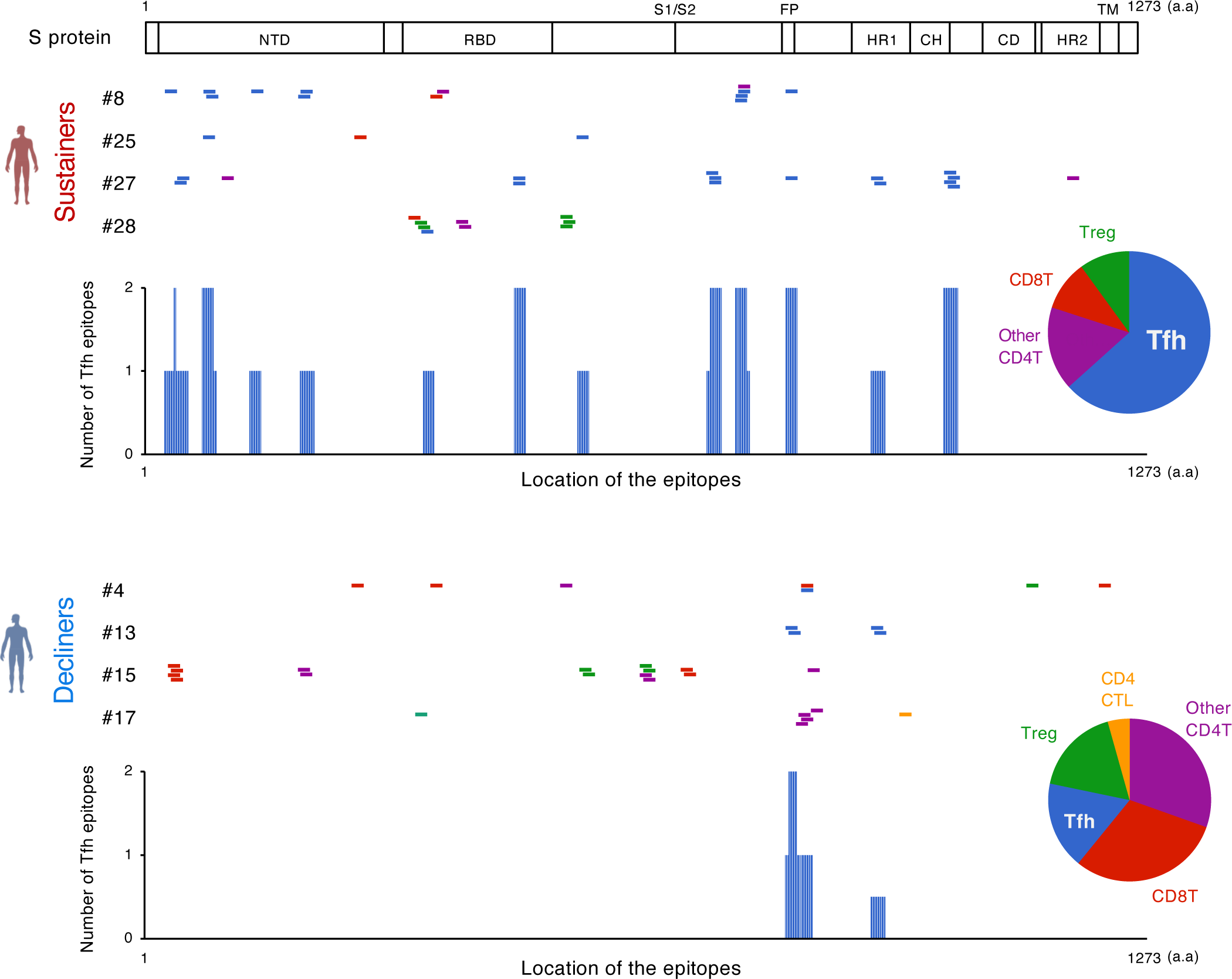
The location of S epitopes recognized by top expanded T clonotypes from post-vaccination samples. T cell S epitopes recognized by top expanded TCR clonotypes in post-vaccinated samples from sustainers and decliners are mapped by their locations in S protein. Each short bar indicates a 15-mer peptide that activated the TCRs. Epitopes are shown in different colors according to the subsets of the T cells they activated. Relative frequencies of the T cell subsets are shown in pie charts. Numbers of identified epitopes recognized by a dominant T subset in sustainers (Tfh) are shown in blue bars. NTD, N-terminal domain; RBD, receptor-binding domain; FP, fusion peptide; HR1, heptad repeat 1; CH, central helix; CD, connector domain; HR2, heptad repeat 2; TM, transmembrane domain.

**Table 4.**
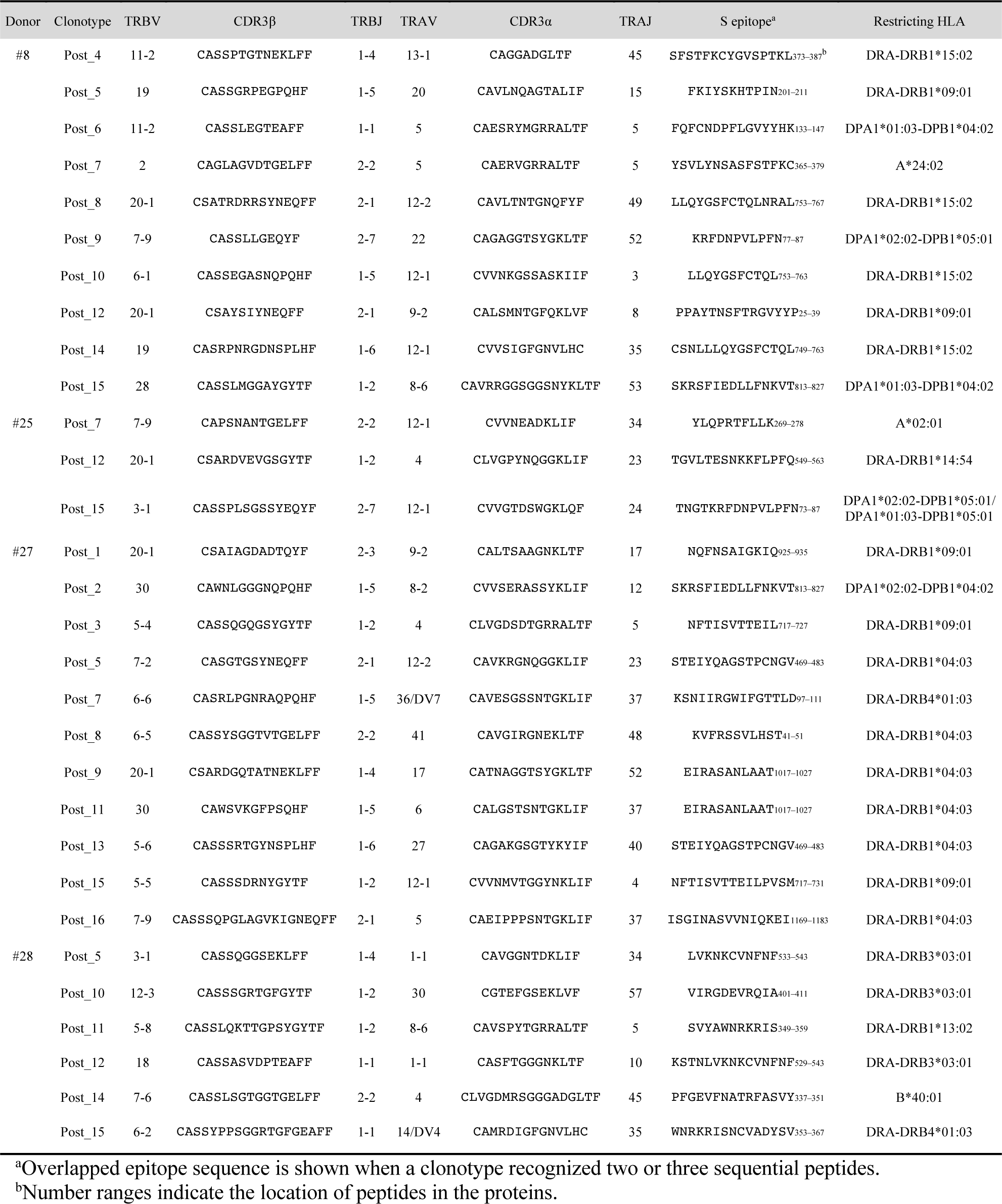
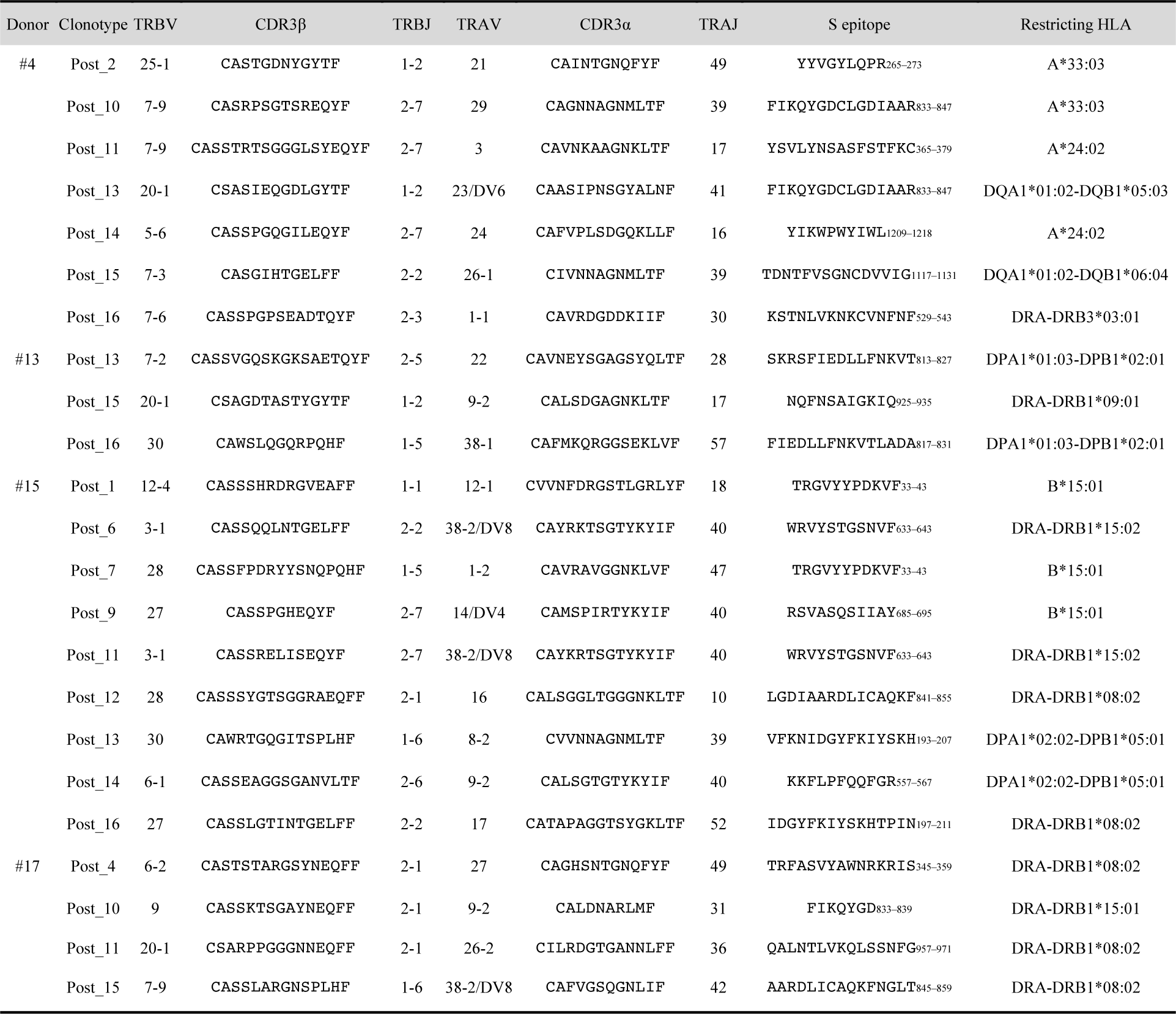
TCR clonotypes expanded in post-vaccinated samples and their TCR usages, epitopes and restricting HLAs.

**Table 5.**
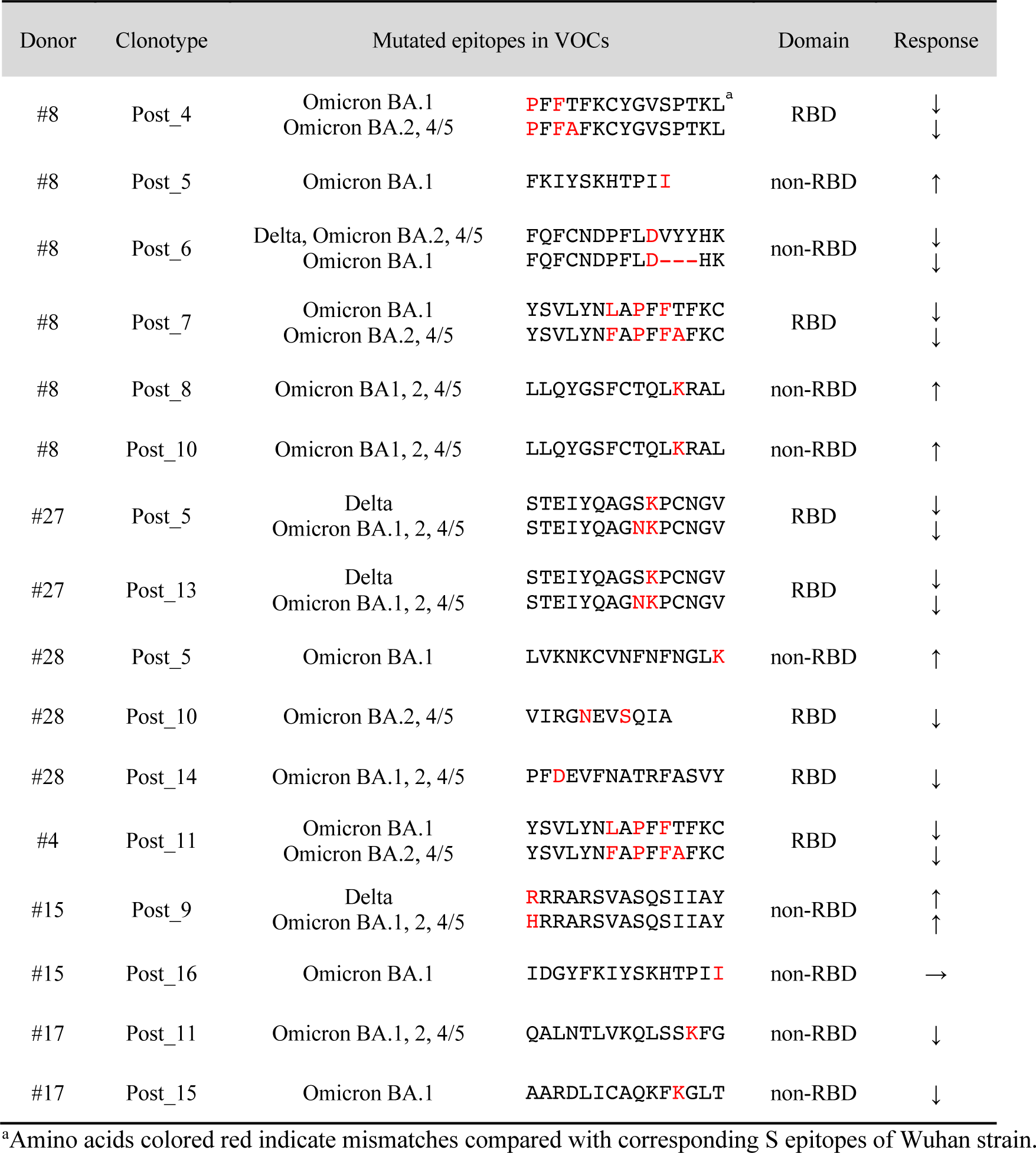
Reactivity of each clonotype to mutated epitopes in SARS-CoV-2 VOCs.

### Identification of S epitopes and cross-reactive antigens of pre-existing T cell clonotypes

Before the pandemic, T cells cross-reacting to S antigen were present in the peripheral blood [13–17]. To characterize these pre-existing S-reactive cells, we analyzed the PBMCs collected from donors who consented to blood sample donation before vaccination (#4, #8, #13, #15, and #17). PBMCs were stimulated with the S peptide pool for 10 days, and proliferated T cells were sorted and analyzed by scTCR/RNA-seq. Similar to vaccine-induced S-reactive T cells (Fig. 2B), characteristics of pre-existing S-reactive T cells were diverse (Fig. 4A). To track the dynamics of cross-reactive clones after vaccination, we combined the single-cell sequencing data of pre- and post-vaccinated PBMCs and analyzed the clonotypes that have more than 50 cells in total (Fig. 4B). We did find some cross-reactive clonotypes that were further expanded by vaccination, and most of these clonotypes had cytotoxic features, being CD8^+^ effector memory T cells (Tem) or minor CD4^+^ cytotoxic T cells (CTLs). In contrast, most of the cross-reactive CD4^+^ T cells became minor clonotypes after vaccination.

**Fig. 4.**
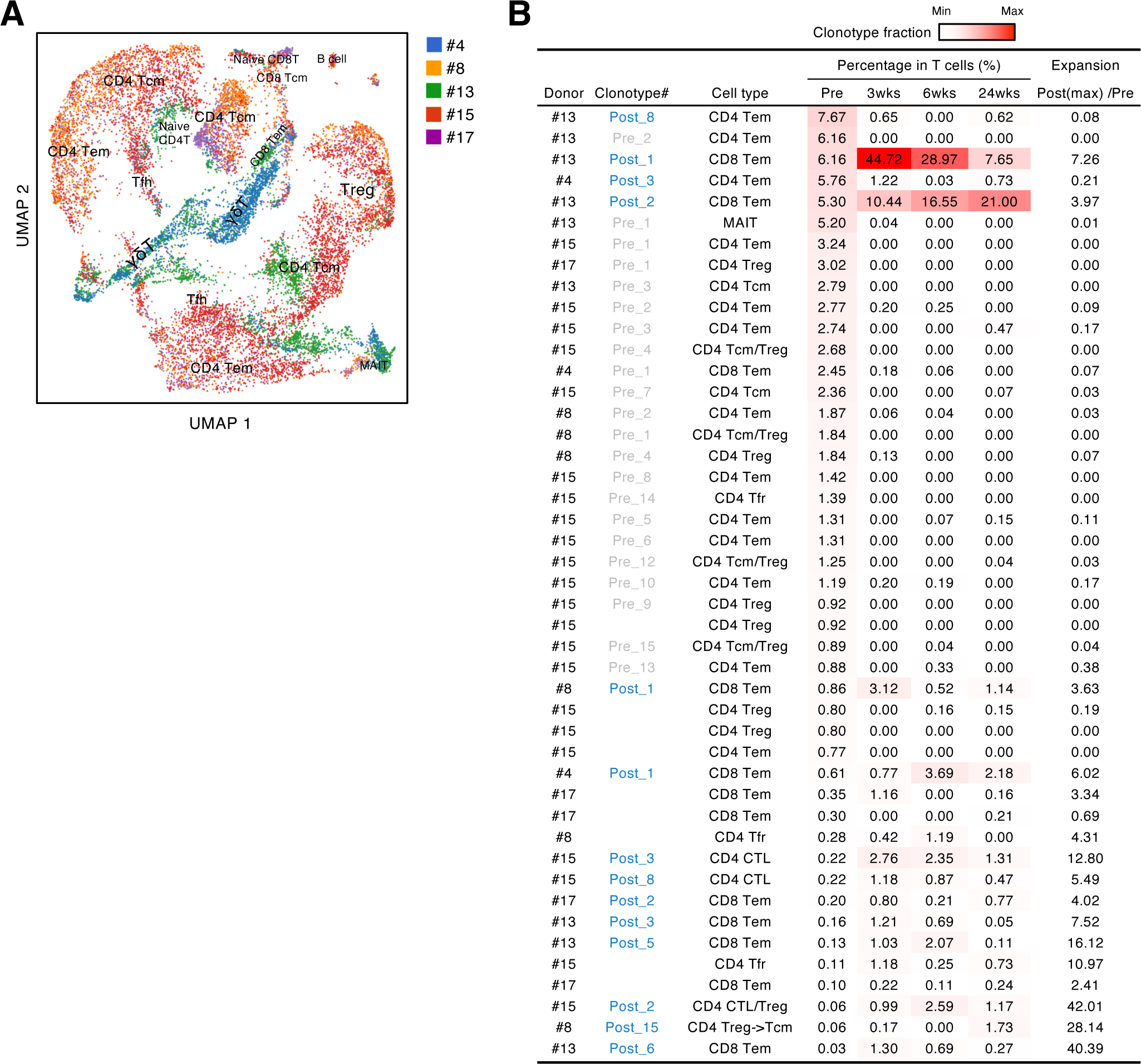
Characteristics and dynamics of S-cross-reactive clonotypes. (**A**) UMAP projection of T cells in single-cell analysis of pre-vaccinated T cells from donors #4, #13, #15, #17, and #8. Each dot corresponds to a single cell and is colored according to the samples from different donors. Annotated cell types are shown. (**B**) Donor, name of reconstituted clonotypes, cell type, clonotype fraction in T cells from each time points, and expansion ratio of clonotypes that were found in pre-vaccinated samples and had more than 50 cells in the combined pre- and post-vaccinated sample set. For clonotypes that showed more than one type, the major type is listed in the front. The expansion ratio was calculated using the maximum cell fraction at post-vaccination points divided by the cell fraction at the pre-vaccination point of each clonotype. Clonotypes that have an expansion ratio larger than 1 are considered as expanded post-vaccination. Cell fractions at individual time points are shown as heat map. Tfr, follicular regulatory T cells; MAIT, mucosal-associated invariant T cells.

We also explored the epitopes of the top 16 expanded clonotypes in each pre-vaccinated donor by reconstituting the TCRs into reporter cell lines. We identified 18 epitopes from S protein and determined some possible cross-reactive antigens (Fig. 5, Table 6, Fig. S5). Most of these cross-reactive antigens originated from environmental or symbiotic microbes (Table 6). Furthermore, majority of the reactive T clonotypes showed regulatory T cell (Treg) signatures (Fig. 5). Six of these 80 analyzed clonotypes could also be frequently detected in the public TCR database Adaptive [23,24]. Notably, most of these clonotypes, except for one case, showed comparable frequencies between pre-pandemic healthy donors and COVID-19 convalescent patients (Fig. 6), suggesting that these clonotypes did not expand upon SARS-CoV-2 infection, despite they were present before the pandemic. Thus, it is unlikely that these cross-reactive T clonotypes contribute to the establishment of S-reactive T cell pools during either vaccination or infection.

**Fig. 5.**
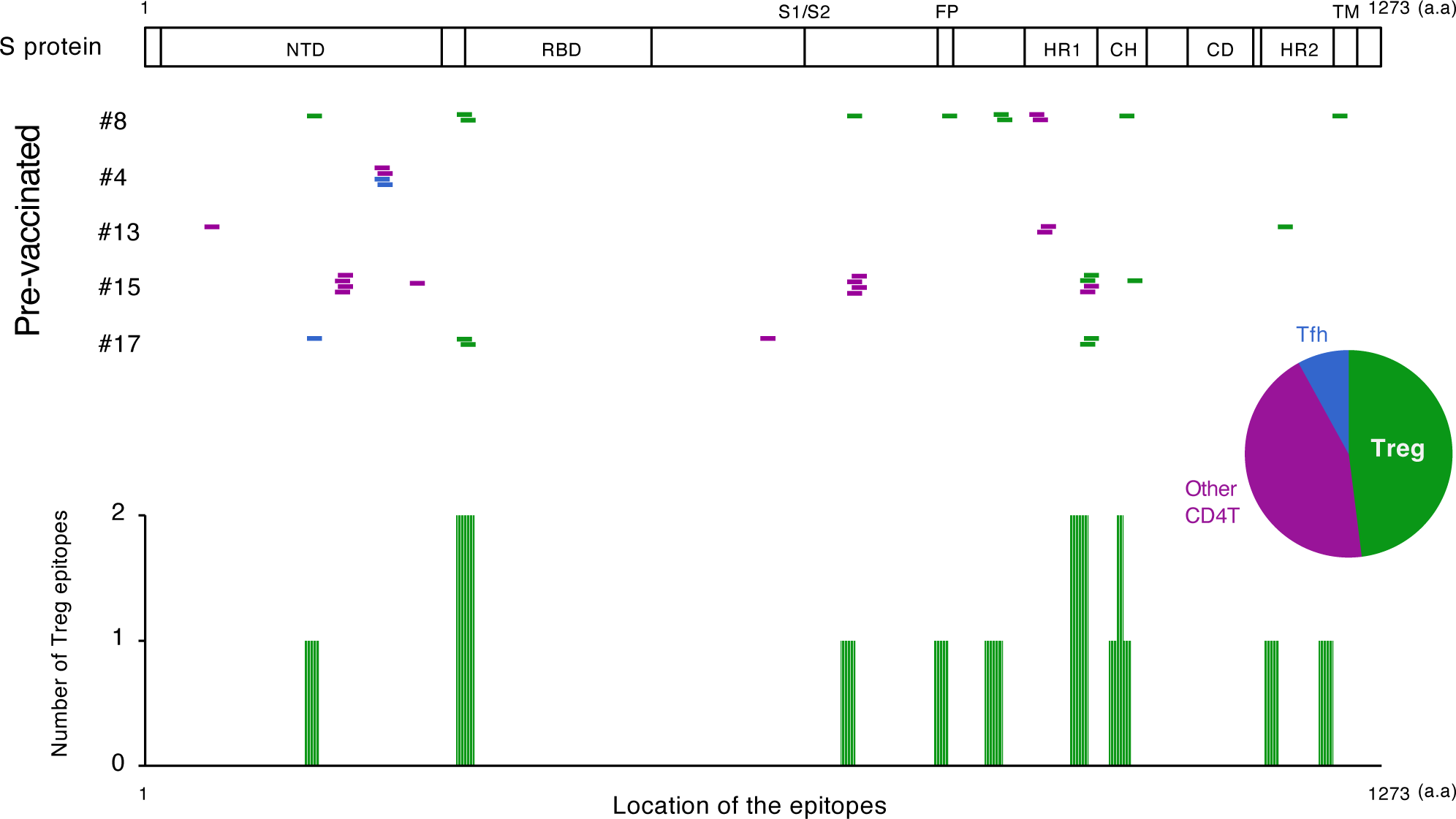
The location of S epitopes of pre-existing S-reactive T cells. S epitopes recognized by top expanded TCR clonotypes in pre-vaccinated samples are mapped by their locations in S protein. Each short bar indicates a 15-mer peptide that activated the TCRs. Epitopes are shown in different colors according to the subtypes of the T cells they activated. Relative frequencies of the T cell subtypes from all five donors are shown in the pie chart. Numbers of identified epitopes recognized by a dominant T subset of pre-existing clonotypes (Treg) from all donors are shown in green bars.

**Fig. 6.**
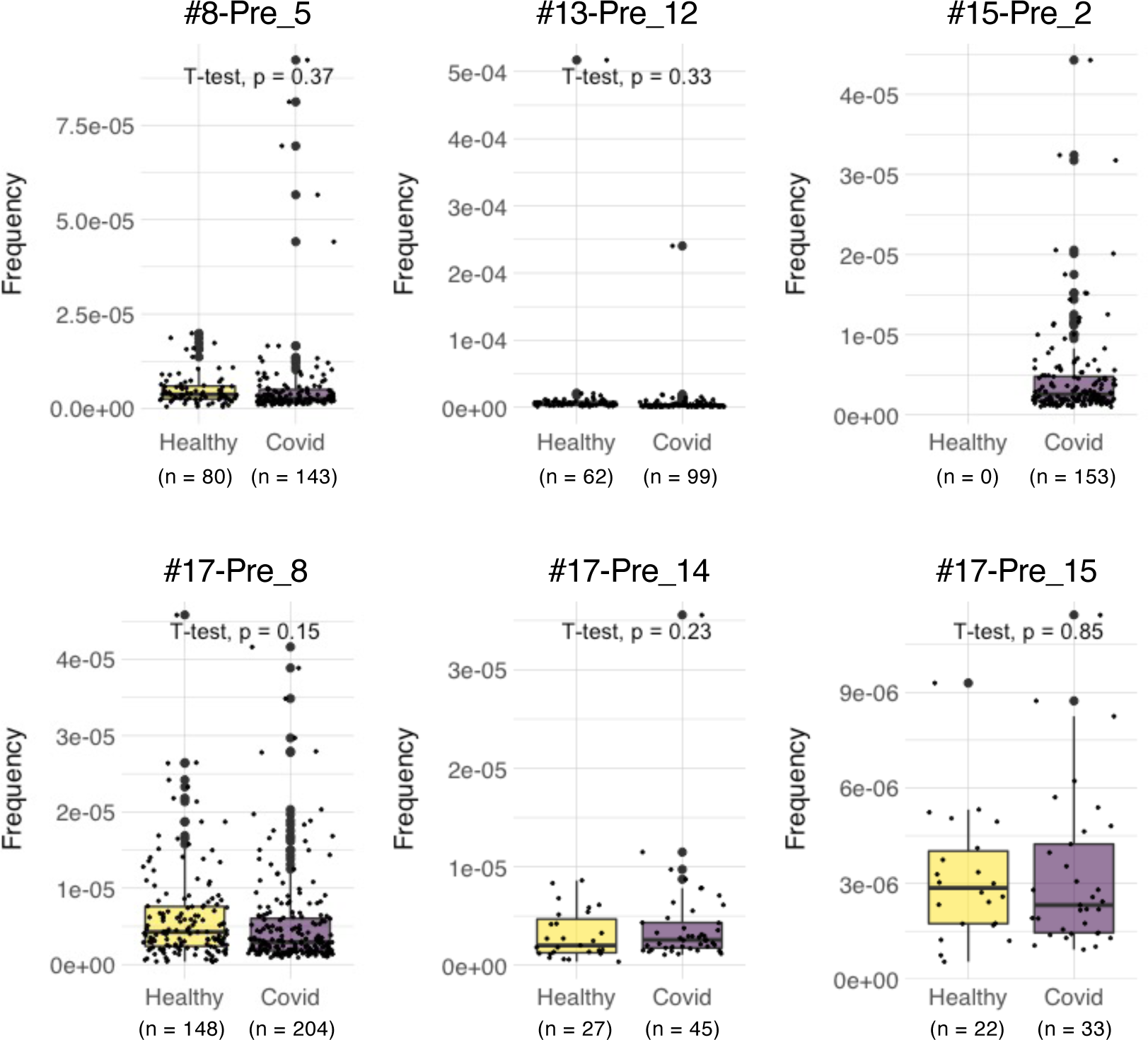
Frequencies of pre-existing S-reactive clonotypes in the public database of uninfected and infected cohorts. TCRβ sequences of the top expanded clonotypes in pre-vaccinated samples were investigated in the Adaptive database. Frequencies of detected clonotypes are shown in box plot. Healthy, dataset from 786 healthy donors. COVID, dataset from 1487 COVID-19 patients.

**Table 6.** S-cross-reactive TCR clonotypes expanded in pre-vaccinated samples and their TCR usages, epitopes, restricting HLAs and cross-reactive epitopes in microbes other than SARS-CoV-2.

| Donor | Clonotype | TRBV | CDR3β | TRBJ | TRAV | CDR3α | TRAJ | S epitope | Restricting HLA | Cross-reactive antigen [species] | Cross-reactive peptide | Post-vaccinated expansion |
| --- | --- | --- | --- | --- | --- | --- | --- | --- | --- | --- | --- | --- |
| #4 | Pre_5 | 6-6 | CASSYPGGGGSETQYF | 2-5 | 35 | CAGVAVQGAQKLVF | 54 | LLALHRSYLT<sup>a</sup><sub>241-251</sub> | DRA-DRB1*14:54 | Phosphoribosylformylglycinamide cyclo-ligase [*Firmicutes bacterium*] | VAEALLAVHRSYLT<sup>b</sup><sub>220-234</sub> | No |
| #4 | Pre_7 | 6-6 | CASSYPGGSGGELFF | 2-2 | 21 | CAVENSNTPLVF | 29 | LLALHRSYLT<sup>a</sup><sub>241-251</sub> | DQA1*01:04-DQB1*05:03 | Phosphoribosylformylglycinamide cyclo-ligase [*Firmicutes bacterium*] | VAEALLAVHRSYLT<sup>b</sup><sub>220-234</sub> | No |
| #8 | Pre_1 | 6-2 | CASRPNRGRFRGNQPQHF | 1-5 | 23/DV6 | CAGEEKETSGSRLTF | 58 | NCTFEYVSQPFLMDL<sup>a</sup><sub>165-179</sub> | DRA-DRB1*15:02 | Fumarylacetoacetate hydrolase family protein [*Alcaligenes faecalis*]<br>Hypothetical protein [*Planctomycetales bacterium*] | ASLIEYVSQPFLEP<sup>c</sup><sub>225-239</sub><br>AAGFEYVSQPFSLPL<sup>c</sup><sub>533-547</sub> | No |
| #8 | Pre_2 | 6-1 | CASIRDRVADTQYF | 2-3 | 30 | CGTETDTSWGKLQF | 24 | RFNGIGVTQNV<sup>a</sup><sub>905-915</sub> | DQA1*03:02-DQB1*03:03 | SEL1-like repeat protein [*Bacteroidaceae bacterium*]<sup>c</sup> | LGYYYFNGIGVTQDQ<sup>c</sup><sub>236-250</sub> | No |
| #8 | Pre_3 | 27 | CATKGEANYGYTF | 1-2 | 12-3 | CAMSEMTGFQKLVF | 8 | SIVRFPNITNL<sup>a</sup><sub>4325-335</sub> | DRA-DRB1*15:02 | LTA synthase family protein [*Dechloromonas denitrificans*] | LPKGSVVRWPNITNL<sup>c</sup><sub>330-344</sub> | No |
| #8 | Pre_5 | 5-1 | CASSLRTGELFF | 2-2 | 8-1 | CAVNGRNTGFQKLVF | 8 | NFTISVTTEILPVSM<sup>a</sup><sub>717-731</sub> | DRA-DRB1*09:01 | Major capsid protein [*Human papillomavirus 145*]<br>Periplasmic trehalase [*Chlamydia bacterium*] | NFTISVTTDAGDINE<sup>c</sup><sub>350-364</sub><br>LSTIVTTEILPVDL<sup>c</sup><sub>288-301</sub> | No |
| #8 | Pre_9 | 7-2 | CASAAGTGGETQYF | 2-5 | 5 | CAETPFLSGTYKYIF | 40 | YIKWPWYIWLGFIA<sup>a</sup><sub>1209-1223</sub> | DRA-DRB5*01:02 | Spike glycoprotein [*Human coronavirus HKU1*] | VKWPWYVWLLISFSF<sup>c</sup><sub>1297-1311</sub> | No |
| #8 | Pre_10 | 6-6 | CASSLGQGIHEQYF | 2-7 | 26-1 | CIVERGGSNYKLTf | 53 | SKRSFIEDLLFNKVT<sup>a</sup><sub>813-827</sub> | DPA1*01:03-DPB1*04:02 | Hypothetical protein, partial [*Acinetobacter baumannii*]<br>Spike protein [*Feline coronavirus*]<br>Spike protein [*Canine coronavirus*] | GKRSAVEDLLFNKVV<sup>c</sup><sub>204-218</sub><br>GKRSAVEDLLFNKVV<sup>c</sup><sub>980-994</sub><br>GKRSAVEDLLFNKVV<sup>c</sup><sub>977-991</sub> | No |
| #8 | Pre_14 | 4-3 | CASSQRQAGDGTQYF | 2-3 | 19 | CALSEAGIQGAQKLVF | 54 | IDRLITGRLQSLQTY<sup>a</sup><sub>993-1007</sub> | DQA1*01:03-DQB1*06:01 | Excinuclease ABC subunit UvrA [*Lentisphaeria bacterium*] | VDRLITGRLESSRLN<sup>c</sup><sub>208-222</sub> | No |
| #8 | Pre_15 | 20-1 | CSAKDRIYGYTF | 1-2 | 26-1 | CIVRSPSGSARQLTF | 22 | MIAQYTSALLA<sup>a</sup><sub>869-879</sub> | DRA-DRB1*15:02 | MATE family efflux transporter [*Selenomonas noxia*] | ATTIAQYTSALLALR<sup>c</sup><sub>242-256</sub> | No |
| #13 | Pre_5 | 4-3 | CASSQVSTGTGITGANVLTF | 2-6 | 5 | CARRSSSASKIIF | 3 | QNVLYENQKL<sup>a</sup><sub>913-923</sub> | DRA-DRB5*01:01 | Hypothetical protein [*Neobacillus vireti*] | TNVLYENQKLFLNL<sup>c</sup><sub>169-183</sub> | No |
| #13 | Pre_8 | 18 | CASSPRAPPYEQYF | 2-7 | 21 | CAVRPAGGTGNQYF | 49 | DKYFKNHTSPDVLG<sup>a</sup><sub>1153-1167</sub> | DRA-DRB1*15:01 | Type VI secretion system contractile sheath large subunit [*Salmonella enterica*] | DYYFDHTSPDVLGL<sup>c</sup><sub>167-181</sub> | No |
| #13 | Pre_12 | 4-2 | CASSQEGNTEAFF | 1-1 | 20 | CGCRGGSYSGKLTf | 52 | NVTWFHAIHVSGTNG<sup>a</sup><sub>61-75</sub> | DQA1*01:02-DQB1*06:02 | Dihydrofolate synthase [*Actinobaculum sp.* 313] | PQRSFHAIHVGTGTNG<sup>c</sup><sub>61-75</sub> | No |
| #15 | Pre_1 | 20-1 | CSARDLTASAHGYTF | 1-2 | 17 | CATDAGQGGKLIF | 23 | SVTTEILPVSM<sup>a</sup><sub>721-731</sub> | DQA1*01:03-DQB1*06:01 | Hypothetical protein [*Mycococcales bacterium*] | PVTTEILPVSDDDPG<sup>c</sup><sub>525-539</sub> | No |
| #15 | Pre_2 | 24-1 | CATSDLQPPQHF | 1-5 | 16 | CALSGYSGYSTLTf | 11 | SVTTEILPVSM<sup>a</sup><sub>721-731</sub> | DQA1*01:03-DQB1*06:01 | Hypothetical protein [*Mycococcales bacterium*] | PVTTEILPVSDDDPG<sup>c</sup><sub>525-539</sub> | No |
| #15 | Pre_3 | 6-1 | CASDPKNGGEQYF | 2-7 | 29/DV5 | CAASVGFGNVLHC | 35 | FKIYSKHTPIN<sup>a</sup><sub>201-211</sub> | DRA-DRB5*01:02 | Uncharacterized protein APUU_31289S [*Aspergillus puulaauensis*] | CRAAFKLYSKHTPVE<sup>c</sup><sub>123-137</sub> | No |
| #15 | Pre_4 | 19 | CASGLAGNTGELFF | 2-2 | 10 | CVPSSGGYNKLIF | 4 | QALNTLVKQLS<sup>a</sup><sub>957-967</sub> | DRA-DRB1*08:02 | 4-hydroxybenzoate octaprenyltransferase [*Pseudoduganella dura*] | IQPLNTLVKQLSVAA<sup>c</sup><sub>112-126</sub> | No |
| #15 | Pre_5 | 6-5 | CASSAGLAGGNTQYF | 2-3 | 5 | CAVISGSARQLTF | 22 | QALNTLVKQLS<sup>a</sup><sub>957-967</sub> | DRA-DRB1*08:02 | 4-hydroxybenzoate octaprenyltransferase [*Pseudoduganella dura*] | IQPLNTLVKQLSVAA<sup>c</sup><sub>112-126</sub> | No |
| #15 | Pre_6 | 2 | CASVGGNEQFF | 2-1 | 9-2 | CALTRFVGATNKLI | 32 | RTFLKYNENGTITD<sup>a</sup><sub>273-287</sub> | DRA-DRB1*15:02 | Unnamed protein product [*Mytilus edulis*] | NKKLLKYNENGTFIT<sup>c</sup><sub>277-291</sub> | No |
| #15 | Pre_7 | 4-1 | CASSHDGTPPDQYF | 2-3 | 29/DV5 | CAAYSNYQLIW | 33 | FKIYSKHTPIN<sup>a</sup><sub>201-211</sub> | DRA-DRB1*15:02 | Uncharacterized protein APUU_31289S [*Aspergillus puulaauensis*] | CRAAFKLYSKHTPVE<sup>c</sup><sub>123-137</sub> | No |
| #15 | Pre_15 | 2 | CASSETGRGTDQYF | 2-3 | 9-2 | CALYRGTYKYIF | 40 | LQSLQTYVTQQLIRA<sup>a</sup><sub>1001-1015</sub> | DRA-DRB1*15:02 | Dyp-type peroxidase [*Acinetobacter sp.*] | CTVLQTYVTQQLESV<sup>c</sup><sub>134-148</sub> | No |
| #17 | Pre_7 | 6-1 | CASSLRGAFGYTF | 1-2 | 35 | CAGHLYGGSQGNLIF | 42 | NCTFEYVSQPFLMDL<sup>a</sup><sub>165-179</sub> | DPA1*01:03-DPB1*04:02 | Fumarylacetoacetate hydrolase family protein [*Alcaligenes faecalis*]<br>Hypothetical protein [*Planctomycetales bacterium*] | ASLIEYVSQPFLEP<sup>c</sup><sub>225-239</sub><br>AAGFEYVSQPFSLPL<sup>c</sup><sub>533-547</sub> | No |
| #17 | Pre_8 | 5-1 | CASSLNSGANVLTF | 2-6 | 13-1 | CAASIVDQKLVF | 8 | LTPTRVYSTGSNVF<sup>a</sup><sub>629-643</sub> | DRA-DRB1*08:02 | Hypothetical protein [*Novosphingobium chloroacetimidivorans*] | APGTPTRVYSTART<sup>c</sup><sub>277-291</sub> | No |
| #17 | Pre_14 | 5-1 | CASSLGAGLYNEQFF | 2-1 | 38-1 | CAFINNAGNMLTF | 39 | QALNTLVKQLS<sup>a</sup><sub>957-967</sub> | DRA-DRB1*08:02 | 4-hydroxybenzoate octaprenyltransferase [*Pseudoduganella dura*] | IQPLNTLVKQLSVAA<sup>c</sup><sub>112-126</sub> | No |
| #17 | Pre_15 | 7-2 | CASSRTSGGTQYQYF | 2-7 | 25 | CAGQNTDKLIF | 34 | SIVRFPNITNL<sup>a</sup><sub>4325-335</sub> | DRA-DRB1*15:01 | LTA synthase family protein [*Dechloromonas denitrificans*] | LPKGSVVRWPNITNL<sup>c</sup><sub>330-344</sub> | Yes |

<sup>a</sup>Number ranges indicate the location of peptides in the proteins.
<sup>b</sup>Amino acids colored red indicate mismatches compared with corresponding S epitopes of Wuhan strain.
<sup>c</sup>Antigen names and peptide sequences in light gray indicate inactive antigens of the corresponding T clonotypes.

## Discussion

Previous studies showed that Tfh function and germinal center development were impaired in deceased COVID-19 patients [25] and Tfh cell number correlated with neutralizing antibody [26–28]. Consistent with the above studies, we found that the donors having sustained antibody titers between 6 to 24 weeks post-vaccination had more S antigen-responsive Tfh-like clonotypes maintained in the periphery as a memory pool. As circulating Tfh clonotypes can reflect the population of germinal center Tfh cells [29], it is possible that these maintained S-responsive Tfh cells contribute to the prolonged production of anti-S antibodies. These results imply that Tfh polarization of S-reactive T cells in the blood after 2nd vaccination can be a marker for the longevity of serum anti-S antibodies. Although monitoring of S-specific Tfh cells in germinal center is ideal [30], it is currently difficult for outpatients in clinics.

Since the antigen used for BNT162b2 is a full-length S protein from the Wuhan-Hu-1 strain, it is important to estimate whether vaccine-induced Wuhan S-reactive T cells recognize neutralizing antibody-evading VOCs, such as Omicron variants. To investigate the dominant T cell epitopes of vaccine-developed T cells, we utilized a proliferation-based sorting strategy to enrich the S-responsive T cells. The limitation of this strategy is that a 10-day stimulation would change the transcriptional profile and repertoire of T cells. However, this strategy allowed us to select the T cell clonotypes that vigorously responded to the S antigen stimulation, while weakly responsive cells and anergic cells will be less considered, which is exactly in line with our purpose. Consistent with previous reports [31–33], most of the epitopes determined in the current study were conserved in Delta and Omicron (BA.1, BA.2 and BA.4/5) strains, suggesting that vaccine-induced T cells are able to recognize the mutated S proteins from these variants, despite the B epitopes being largely mutated in these VOCs [31,32].

SARS-CoV-2-recognizing T cells existed prior to exposure to the S antigens [13–17], which is consistent with our observation with PBMCs from donors who were uninfected and pre-vaccinated. Among these pre-existing S-reactive clonotypes, CD8^+^ cytotoxic T clonotypes were expanded by the vaccination, whereas most CD4^+^ T clonotypes became less dominant after vaccination (Fig. 4B). Currently, the reason for the opposite tendency is unclear. In the present study, we showed that pre-existing T clonotypes cross-reacting to S protein are unlikely to contribute to vaccine-driven T cell immunity. This could be due to the fact that cross-reactive T cells had relatively low avidity to S protein [34]. Alternatively, but not mutually exclusively, considering that most of these cross-reactive T clonotypes have Treg signature (Fig. 5), they could be developed to tolerate symbiotic or environmental antigens, and might be ineffective to the defense against SARS-CoV-2 and thus replaced by the other effective T clonotypes induced by vaccination. One exceptional pre-existing clonotype was #15-Pre_2, as they vigorously expanded in COVID-19 patients (Fig. 6). This clonotype was clustered within a CD4^+^ Tem population and cross-reactive to environmental bacteria, *Myxococcales bacterium* (Table 6). Thus, in some particular settings, clonotypes primed by common bacterial antigens might potentially contribute during infection.

Common cold human coronavirus (HCoV)-derived S proteins are reported as potential cross-reactive antigens for pre-existing SARS-CoV-2 S-reactive T cells [15,18–20]. However, the highly responding SARS-CoV-2 S-reactive clonotypes in pre-vaccinated donors did not react with HCoV S proteins in the present study (Fig. S6), which might be partly due to the difference of cohorts or ethnicities. Instead, most of those T cells cross-reacted with environmental or symbiotic bacteria. These observations suggest that these cross-reactive T cells might have been developed to establish tolerance against less harmful microbes, and thus unlikely to efficiently contribute to the protective viral immunity. Vaccination may induce opposite tendencies on T cell clonotypes that recognize the same antigen [35], which is hardly detected by the bulk T cell analyses. The current study highlights the necessity of dynamic tracing of T cell responses in an epitope-specific clonotype resolution for the evaluation of vaccine-induced immunity.

The limitation of this study is the number of individuals we analyzed. However, chronological and clonological analysis of antigen-specific T cells in characteristic groups followed by epitope determination has not been performed before. This study suggests that mRNA vaccine is potent enough to prime rare T cell clonotypes that become dominant afterwards. Furthermore, we propose that the types of CD4^+^ T clonotypes developed shortly after two doses of vaccination could be an indication of the longevity of antibodies in the following months. Tfh-inducing adjuvants or Tfh-skewing epitope would be a promising “directional” booster in the post-vaccine era when most people worldwide were exposed to the same antigen in multiple doses within a short period. Furthermore, in addition to SARS-CoV-2, this strategy can also be applicable for the prevention of other infectious diseases of which neutralizing antibody titers are effective for protection.

## Materials and Methods

### Ethics statement and sample collection

This project was approved by Osaka University Institutional Review Board (IRB) (reference No. 21487). 43 volunteers were enrolled in this project. Informed consent was obtained from all participants before the first blood sampling. Samples (serum, whole blood, and PBMCs) were collected four times at 0–7 days before 1st dose vaccination as pre-vaccination, at 14–21 days after 1st dose vaccination as 3 weeks sample, at 35–49 days after 1st dose vaccination as 6 weeks sample, and at 154–182 days after 1st dose of vaccination as 24 weeks sample. At the same time of blood sampling, adverse event information was also collected from all participants. PBMCs were isolated using BD vacutainer® CPT™ cell separation tube (Beckton Dickinson), according to manufacturers’ instructions. Isolated PBMCs were stored in the vapor phase of liquid nitrogen until use.

### Antibody titer determination by enzyme-linked immunosorbent assay (ELISA)

Serum antibody titer was measured using ELISA. Briefly, recombinant ancestral S protein (S1+S2, Cell Signaling Technology; 1 µg/ml) or recombinant nucleocapsid protein (Acrobiosystems; 1 µg/ml) was coated on 96-well plate at 4 °C overnight. On the second day, wells were blocked with blocking buffer (PBS-T (0.05% tween®20) containing 5% skim milk) for 2 h at room temperature. The sera were diluted from 10 to 31,250 folds in blocking buffer and incubated overnight at 4 °C. The next day, wells were washed and incubated with horseradish peroxidase (HRP)-conjugated antibodies (GE Healthcare) for 3 h at room temperature. After being washed with PBS-T, wells were incubated with the peroxidase chromogenic substrate 3,3’-5,5’-tetramethyl benzidine (Sigma-Aldrich) for 30 min at room temperature, then the reaction was stopped by 0.5 N sulfuric acid (Sigma Aldrich). The absorbance of wells was immediately measured at 450 nm with a microplate reader (Bio-Rad).

The value of the half-maximal antibody titer of each sample was calculated from the highest absorbance in the dilution range by using Prism 8 software. The calculated antibody titer was converted to BAU/ml by using WHO International Standard 20/136 (NIBSC) for ancestral S-specific antibody titer.

### Whole blood interferon-gamma release immune assay (IGRA) for SARS-CoV-2 specific T cell responses using QuantiFERON

SARS-CoV-2 specific T cell immune responses were evaluated by QuantiFERON SARS-CoV-2 (Qiagen) [36], according to manufacturer’s instructions, in which CD4^+^ T cells were activated by epitopes coated on Ag1 tube, and CD4^+^ and CD8^+^ T cells were activated by epitopes coated on Ag2 tube. Briefly, 1 ml of whole blood sample with heparin is added into each of Nil (negative control), Mito (positive control), Ag1, and Ag2 tubes, and incubated at 37 °C for 22–24 h. Tubes were then centrifuged at 3,000 × g for 15 min for collecting plasma samples. IFNγ derived from activated T cells was measured with enzyme-linked immunosorbent assay (ELISA) (Qiangen) according to the manufacturer’s instructions. IFNγ concentration (IU/ml) was calculated with background (Nil tube) subtracted from values of Ag1 or Ag2 tubes.

### Pseudo-typed virus neutralization assay

The neutralizing activity of serum antibodies was analyzed with pseudo-typed VSVs as previously described [37]. Briefly, Vero E6 cells stably expressing TMPRSS2 were seeded on 96-well plates and incubated at 37 °C for 24 h. Pseudoviruses were incubated with a series of dilutions of inactivated serum for 1 h at 37 °C, then added to Vero E6 cells. At 24 h after infection, cells were lysed with cell culture lysis reagent (Promega), and luciferase activity was measured by Centro XS^3^ LB 960 (Berthold).

### In vitro stimulation of PBMCs

Cryopreserved PBMCs were thawed and washed with warm RPMI 1640 medium (Sigma) supplemented with 5% human AB serum (GeminiBio), Penicillin (Sigma), streptomycin (MP Biomedicals), and 2-mercaptoethanol (Nacalai Tesque). PBMCs were labeled with Cell Proliferation Kit (CellTrace™ Violet, ThermoFisher) following the manufacturer’s protocol and were stimulated in the same medium with S peptide pool (1 μg/ml per peptide, JPT) for 10 days, with human recombinant IL-2 (1 ng/ml, Peprotech), IL-7 (5 ng/ml, BioLegend) and IL-15 (5 ng/ml, Peprotech) supplemented on day 2, day 5 and day 8 of the culture. On day 10 cells were washed and stained with anti-human CD3 and TotalSeq-C Hashtags antibodies. Proliferated T cells (CD3^+^CTV^low^) were sorted by cell sorter SH800S (SONY) and used for single-cell TCR and RNA sequencing analyses.

### Single cell-based transcriptome and TCR repertoire analysis

Single cell library was prepared using the reagents from 10x Genomics following the manufacturer’s instructions. After reverse transcription, cDNA was amplified for 14 cycles, and up to 50 ng of cDNA was used for construction of gene expression and TCR libraries. Libraries were sequenced in paired-end mode, and the raw reads were processed by Cell Ranger 6.0.0 (10x Genomics). Distribution of the mitochondrial gene percentage, n_counts and n_genes were fitted with a one-variable, two-component mixed Gaussian model using the Python package scikit-learn [38] and divided into two distributions corresponding to high and low levels, respectively. The cutting threshold values were the middle value of the means of the two fitted Gaussian distributions. A package call Scrublet was also applied [39], and the events whose main hashtag reads are less than 95% of the total hashtag reads were gated out before the UMAP plots were exported using BBrowser [40]. Tfh signature score was generated using canonical Tfh marker genes (*IL21*, *ICOS*, *CD200, PDCD1*, *POU2AF1*, *BTLA*, *CXCR5,* and *CXCL13*). Other cell populations were annotated using the following markers: Treg, *CD4*^+^*FOXP3*^+^; CD4T, *CD3E*^+^*CD4*^+^; CD8T, *CD3E*^+^*CD8A*^+^; central memory (cm) cells, *SELL*(CD62L)^hi^ cells although sometimes *CCR7* expression is vague; effector memory (em) cells, *SELL*^low/–^*CCR7*^-^ and *IFNG*-expressing cells containing populations; naïve cells, *CCR7*^+^*TCF7*^+^; cycling cells, *MKI67*^hi^; γδT, *TRDC*^+^; B cells, *CD19*^+^; Monocyte, *CD14*^+^; MAIT, *CD3E*^+^*KLRB1*^+^*IL18R1*^+^; Tfr, *FOXP3*^+^*NRN1*^+^ in cells with high Tfh score; CD4-CTL, *GZMB*^+^ in CD4T cells [14,41–43].

### Reporter cell establishment and stimulation

TCRα and β chain cDNA sequences were introduced into a mouse T cell hybridoma lacking TCR and having a nuclear factor of activated T-cells (NFAT)-green fluorescent protein (GFP) reporter gene [44] using retroviral vectors. TCR-reconstituted cells were co-cultured with 1 μg/ml of peptides in the presence of antigen-presenting cells (APCs). After 20 h, cell activation was assessed by GFP and CD69 expression.

### Antigen-presenting cells

Transformed B cells and HLA-transfected HEK293T cells used as APCs were generated as described [21]. For transformed B cells, 3 × 10^5^ PBMCs were incubated with the recombinant Epstein-Barr virus (EBV) suspension [45] for 1 h at 37°C with mild shaking every 15 min. The infected cells were cultured in RPMI 1640 medium supplemented with 20% fetal bovine serum (FBS, CAPRICORN SCIENTIFIC GmbH) containing cyclosporine A (CsA, 0.1 μg/ml, Cayman Chemical). Immortalized B lymphoblastoid cell lines were obtained after 3 weeks of culture and used as APCs. For HLA-transfected HEK293T cells, plasmids encoding HLA class I/II alleles [46] were transfected in HEK293T cells with PEI MAX (Polysciences).

### Determination of epitopes and restricting HLA

15-mer peptides with 11 amino acids overlap that cover the full length of S protein of SARS-CoV-2 were synthesized (GenScript). Peptides were dissolved in DMSO at 12 mg/ml and 12-15 peptides were mixed to create 26 different semi-pools. TCR-reconstituted reporter cells were stimulated with 1 μg/ml of S peptide pool (1 μg/ml per peptide, JPT), then 36-peptide pools that consist of 3 semi-pools each, then semi-pools, and then 12 individual peptides in the presence of autologous B cells to identify epitope peptides. To determine the restricting HLA, HLAs were narrowed down by co-culturing reporter cells with autologous and various heterologous B cells in the presence of 1 μg/ml of the epitope peptide. HLAs shared by activatable B cells were transduced in HEK239T cells and used for further co-culture to identify the restricting HLA.

### Statistics

All values with error bars are presented as the mean ± SEM. One-way ANOVA followed by Turkey’s post hoc multiple comparison test was used to assess significant differences in each experiment using Prism 8 software (GraphPad Software). Differences were considered to be significant when *P* value was less than 0.05. *P* values in Fig. 6 were calculated with t-test using the “stat_compare_means” function in R.

## Data availability

Single cell-based transcriptome data have been deposited in Gene Expression Omnibus (GEO) datasets (accession number GSE246535). Other data needed to support the conclusion of this manuscript are included in the main text and Supplemental information.

## Acknowledgements

We thank S. Iwai, Y. Sakai, C. Günther, J. Sun, and Y. Yanagida for experimental support, D. Motooka, D. Okuzaki, and YC. Liu for bioinformatic data analysis and C. Schutt and Y. Yamagishi for discussion. This research was supported by Japan Agency for Medical Research and Development (JP223fa627002, JP223fa727001, JP23ym0126049 (SY), JP21ym0126049 (HN)) and Japan Society for the Promotion of Science Grants-in-Aid for Scientific Research (JP20H00505, JP22H05182, JP22H05183 (SY)). The Department of Health Development and Medicine is an endowed department supported by AnGes, Daicel, and FunPep.

---

**Fig. S1.**
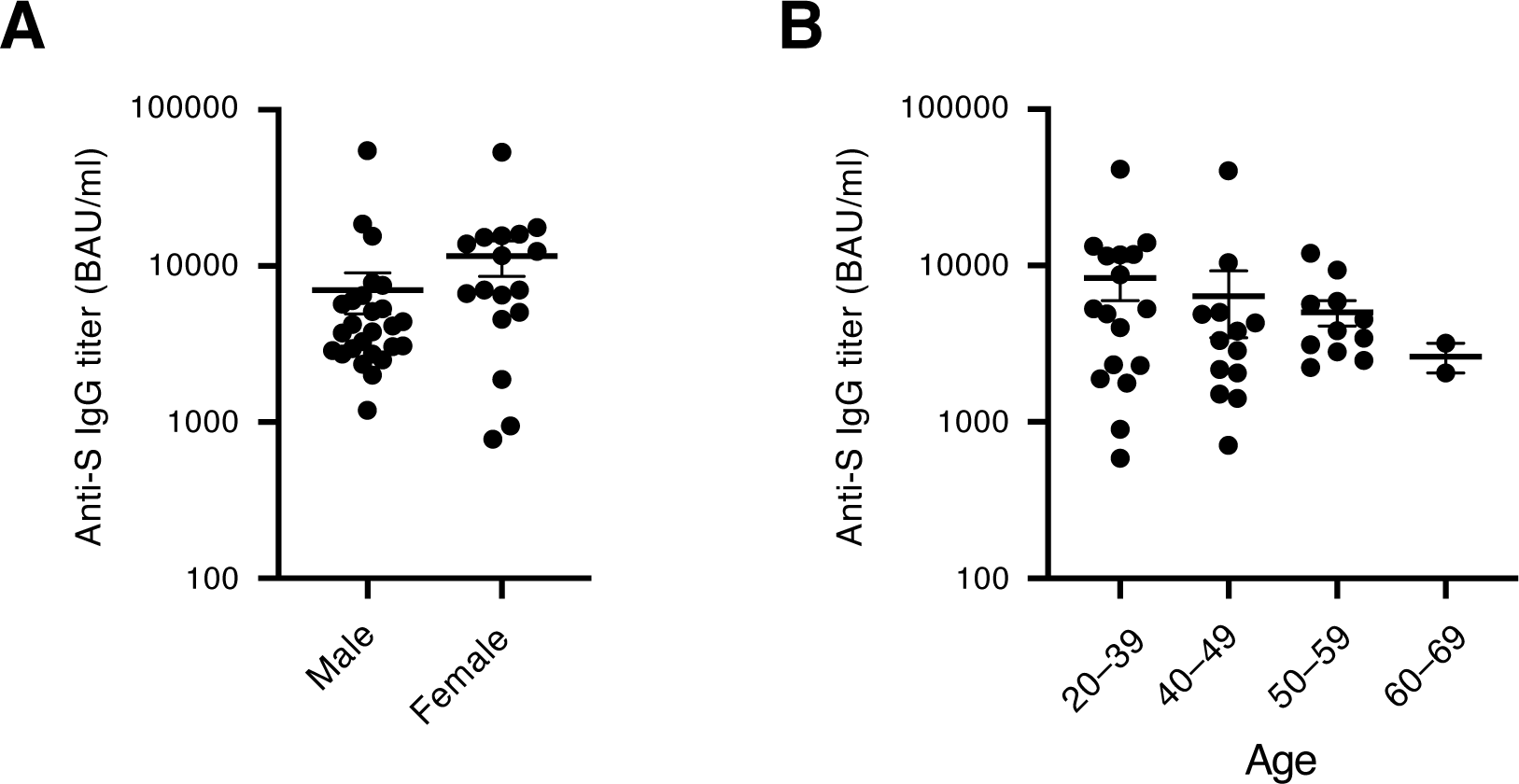
Humoral immune response of BNT162b2 vaccinees. (**A** and **B**) Anti-S IgG titer from all donors at 6 weeks after vaccination was compared between male and female vaccinees (A) or among different age groups (20–39, 40–49, 50–59, 60–69) (B).

**Fig. S2.**
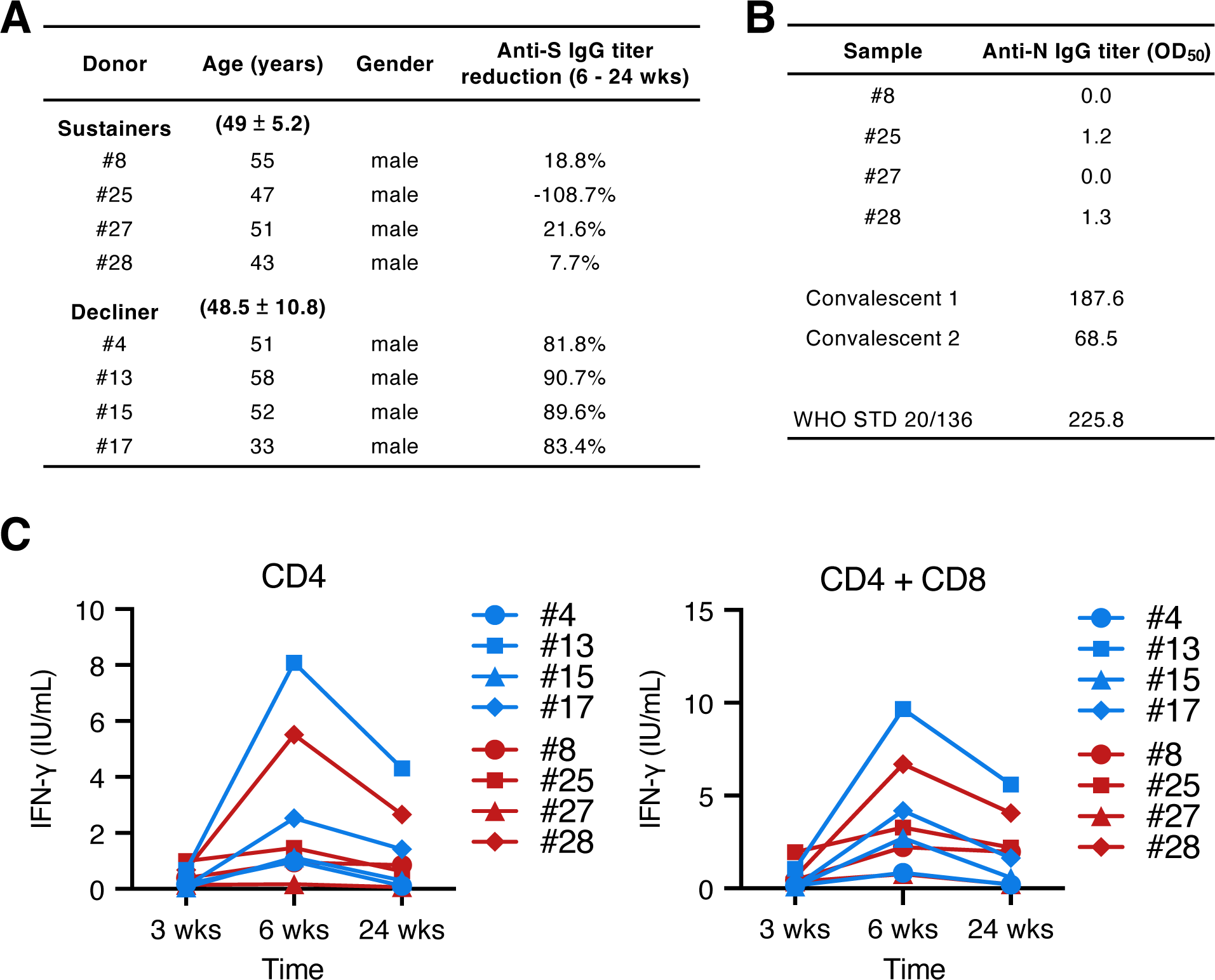
Humoral and cellular immune responses of sustainers and decliners. **(A)** Demographic data and magnitude of anti-S IgG titer reduction of the sustainers and decliners. Anti-S IgG titer reduction is calculated as the titers at (6 wks – 24 wks) /6 wks. **(B)** Anti-N IgG titer of serum samples from sustainers at 24 weeks after vaccination. (**C**) S-specific IFNγ release from bulk CD4^+^ T cells (left) or CD4^+^ and CD8^+^ T cells (right) of sustainers (red) and decliners (blue) was measured using QuantiFERON.

**Fig. S3.**
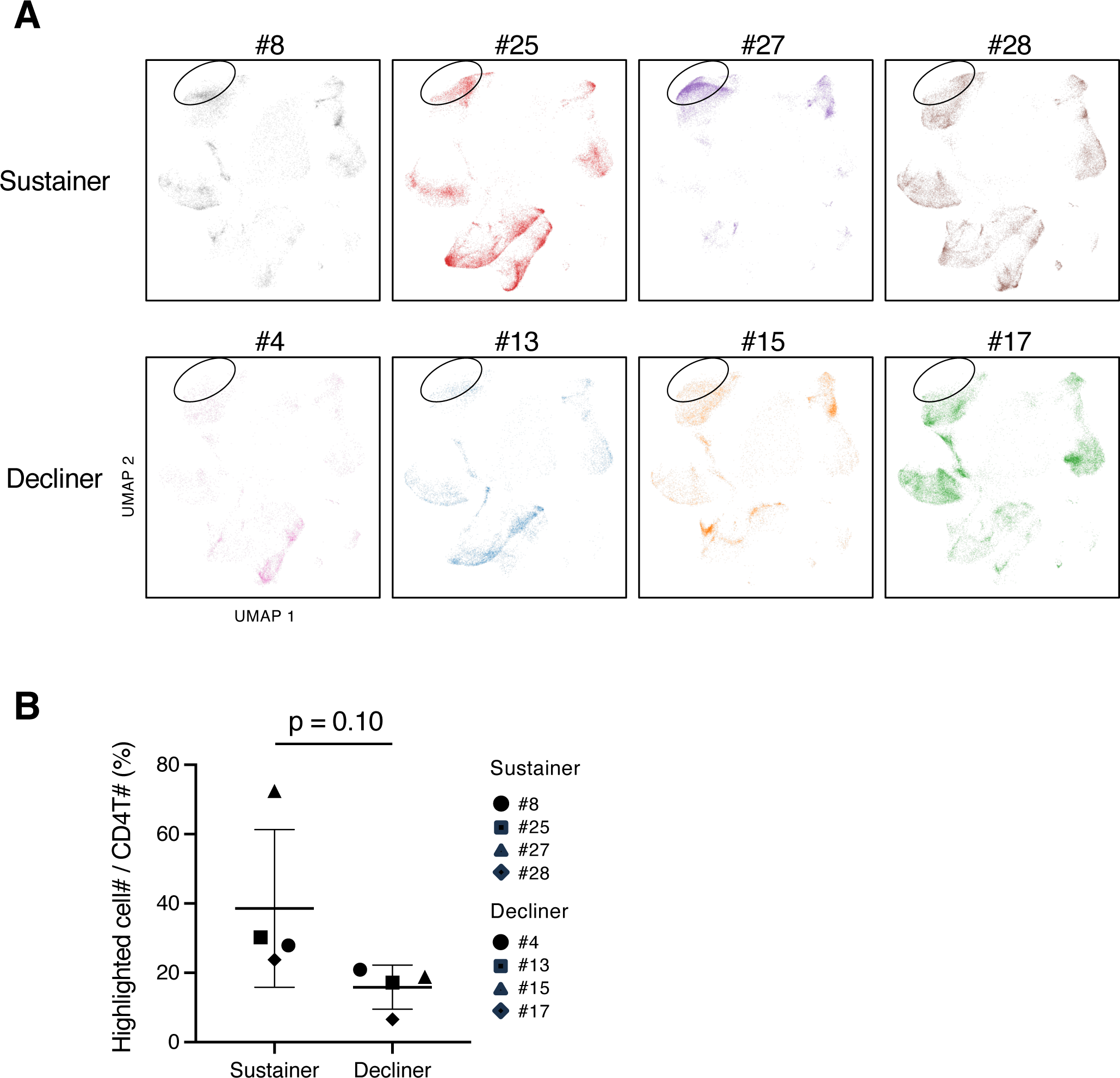
Sustainer individuals had more cells in the circled region than decliner individuals. **(A)** UMAP projection of single-cell analysis of post-vaccinated samples shown in individuals. **(B)** The percentage of circled cells in (A) in CD4^+^ T cells of each individual is shown. *P* value was calculated using t-test.

**Fig. S4.**
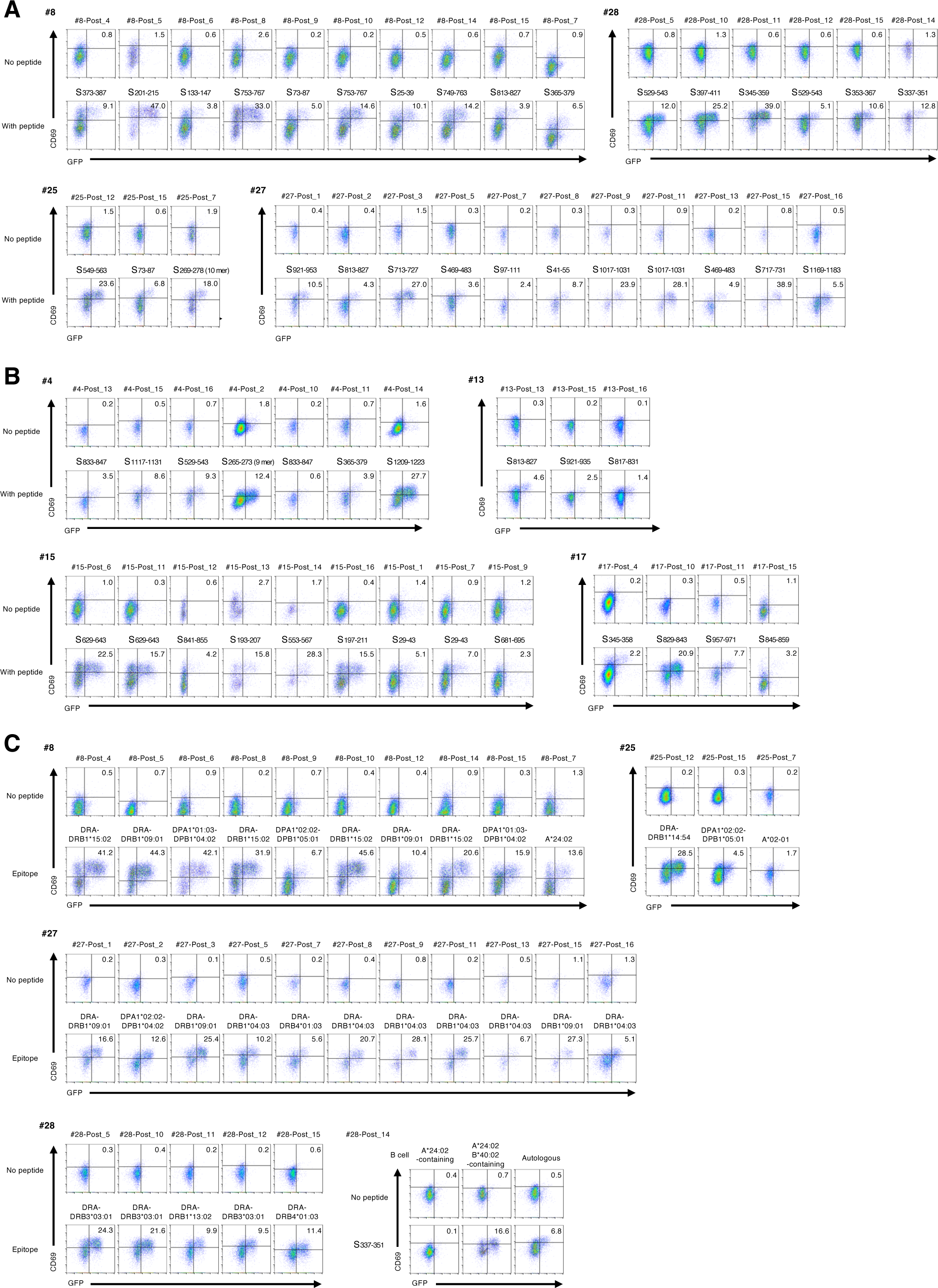
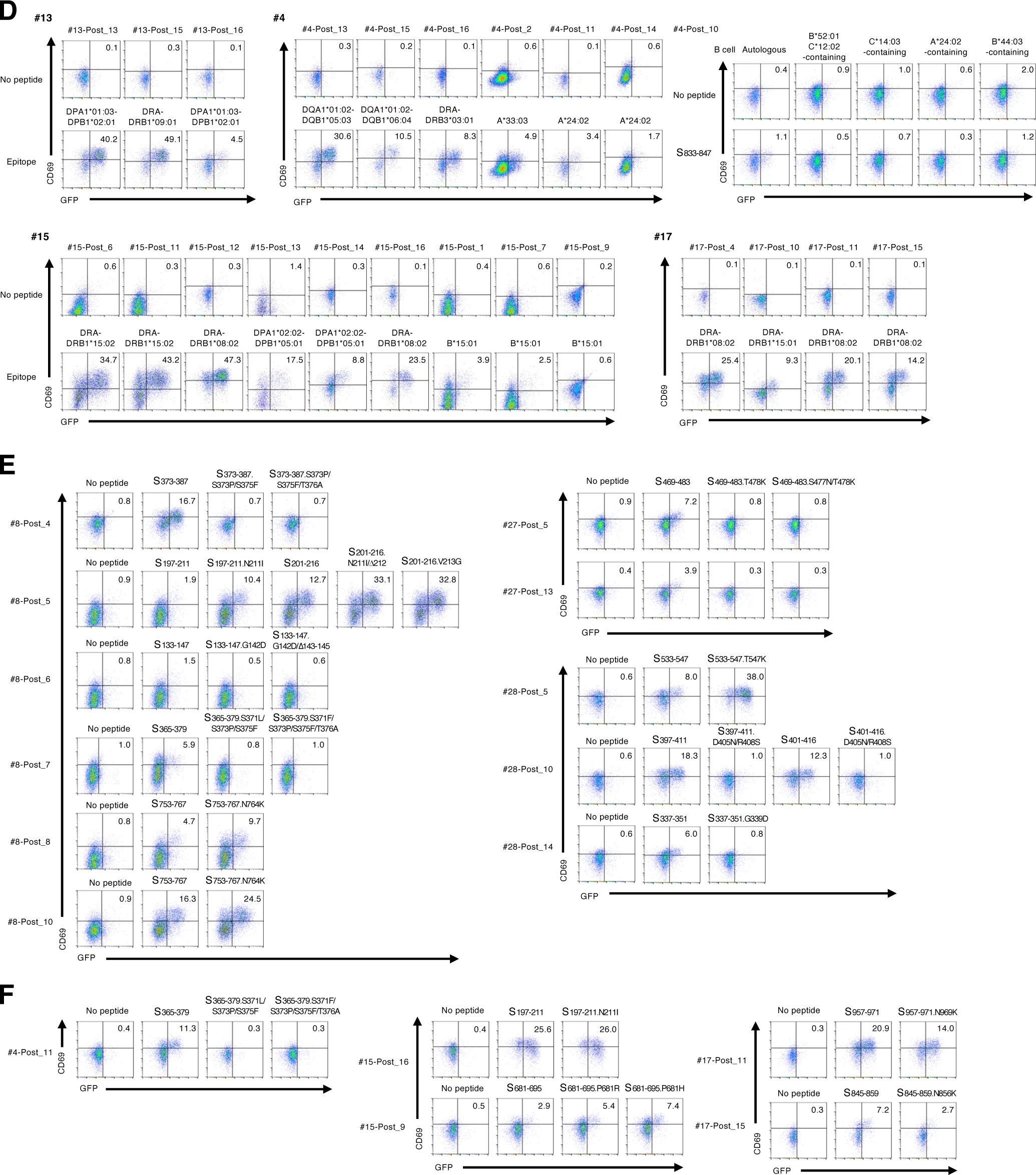
Determination of S epitopes, restricting HLAs and mutated epitope antigenicity for post-vaccinated T cell clonotypes expanded in sustainers and decliners. Reporter cells expressing TCRs were stimulated with 1 µg/ml of indicated S protein peptide in the presence of transformed B cells or HEK 293T cells expressing indicated HLAs for overnight, and analyzed for GFP and CD69 expression. (**A** and **B**) Determination of S epitopes of TCR clonotypes expanded in sustainers (A) and decliners (B). (**C** and **D**) Determination of restricting HLAs of TCR clonotypes expanded in sustainers (C) and decliners (D). (**E** and **F**) Determination of antigenicity of mutated epitopes of TCR clonotypes expanded in sustainers (E) and decliners (F).

**Fig. S5.**
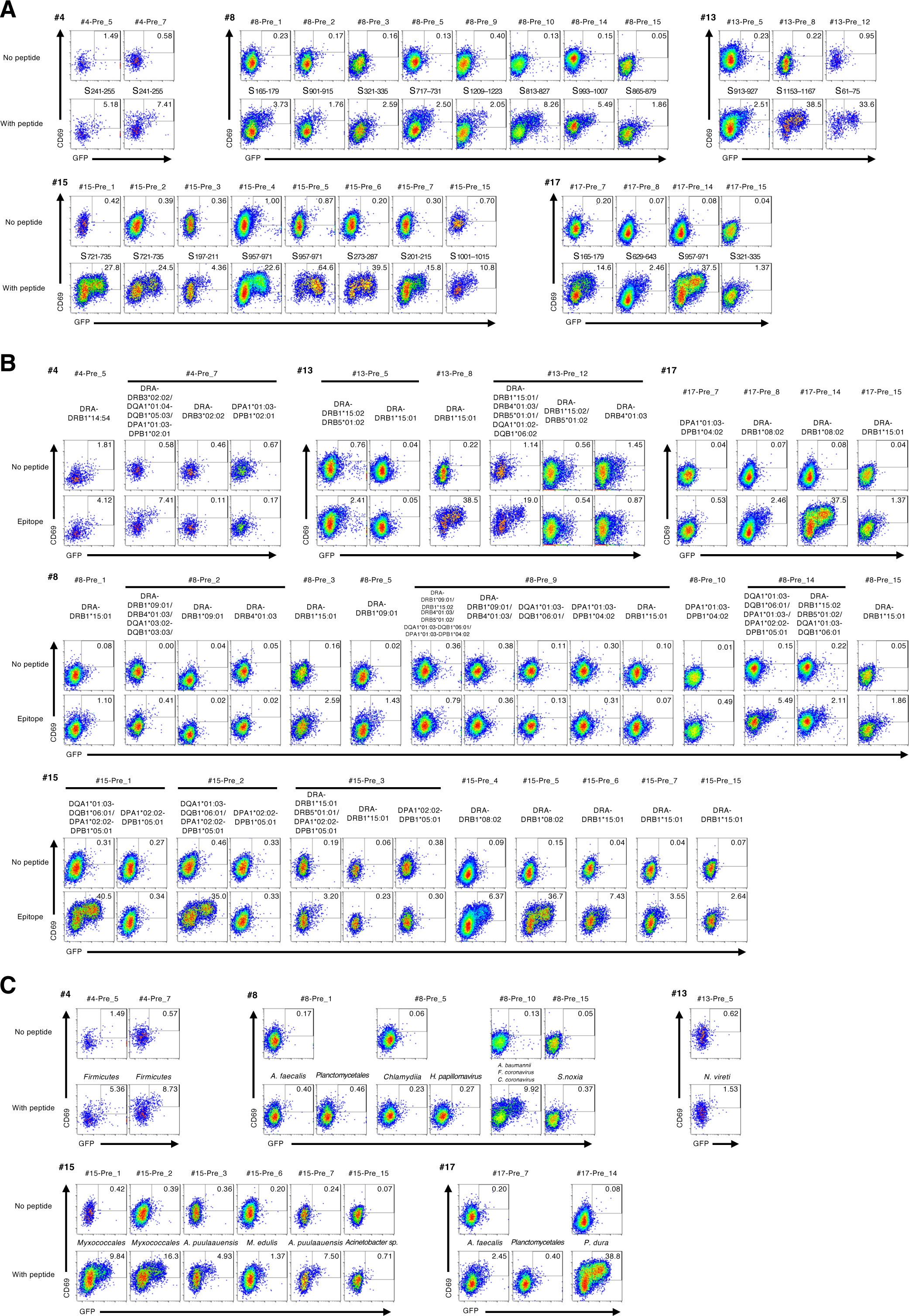
Determination of S epitopes, restricting HLAs and cross-reactive epitopes for pre-existing T cell clonotypes expanded by S stimulation. Reporter cells expressing TCRs were stimulated with 1 µg/ml of indicated S peptides in the presence of transformed B cells or HEK 293T cells expressing indicated HLAs for overnight, and analyzed for GFP and CD69 expression. (**A**) Determination of S epitopes of T cell clonotypes. (**B**) Determination of restricting HLAs of T cell clonotypes. (**C**) Determination of cross-reactive epitopes of T cell clonotypes. Sequences of cross-reactive peptides are in Table 6.

**Fig. S6.**
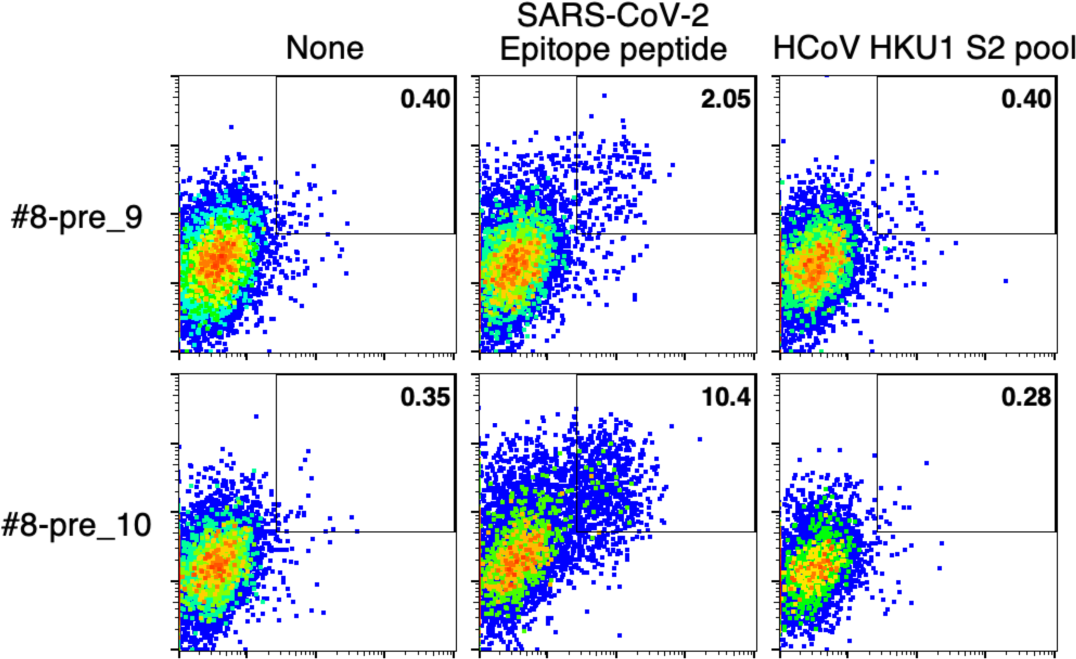
The pre-existing S-reactive T cell clonotypes did not recognize HCoV epitopes. The only two clonotypes whose epitope sequences were relatively conserved in HCoV strains, donor #8-pre_9 and pre_10, were tested for their reactivity to the similar HCoV epitope counterparts. Reporter cell lines of these clonotypes were co-cultured with indicated peptides as well as APCs, and analyzed for GFP and CD69 expression.

