## Supplementary material for "Early acquisition of S-specific Tfh clonotypes after SARS-CoV-2 vaccination is associated with the longevity of anti-S antibodies": Post_vaccination_samples.html

Yamazaki\_Iu\_scRNASeq\_all\_pfizer


In [1]:

```
import import_ipynb
import Scanpy_functions_v03262021 as sc_pipe
import scvelo as scv
scv.logging.print_version()
import warnings
import scirpy as ir
import scanpy as sc
import numpy as np
import scipy as sp
import pandas as pd
import matplotlib.pyplot as plt
from matplotlib import rcParams
from matplotlib import colors
import seaborn as sb
import bbknn
import logging
from sklearn.mixture import GaussianMixture
from scipy.stats     import norm
import glob
import os
import hvplot.pandas
import docx
from docx import Document
from docx.shared import Inches
from docx.shared import Pt
from scipy import sparse
import scanpy.external as sce
import holoviews as hv
import panel as pn
import bokeh
from bokeh.resources import INLINE
import scanorama
import gseapy
```

```
importing Jupyter notebook from Scanpy_functions_v03262021.ipynb
Running scvelo 0.2.3 (python 3.8.5) on 2021-11-25 10:09.
Running scvelo 0.2.3 (python 3.8.5) on 2021-11-25 10:09.
```

In [2]:

```
sc.logging.print_versions()
```

```
WARNING: If you miss a compact list, please try `print_header`!
-----
anndata     0.7.6
scanpy      1.8.1
sinfo       0.3.1
-----
Levenshtein                 NA
PIL                         8.3.1
Scanpy_functions_v03262021  NA
adjustText                  NA
airr                        1.3.1
anndata                     0.7.6
annoy                       NA
anyio                       NA
appdirs                     1.4.4
async_generator             1.10
attr                        21.2.0
babel                       2.9.1
backcall                    0.2.0
bbknn                       NA
bioservices                 1.7.11
bokeh                       2.3.3
bottleneck                  1.3.2
brotli                      NA
bs4                         4.9.3
cairo                       1.19.1
certifi                     2021.05.30
cffi                        1.14.6
chardet                     4.0.0
cloudpickle                 1.6.0
colorama                    0.4.4
colorcet                    1.0.0
colorlog                    NA
cycler                      0.10.0
cython_runtime              NA
cytoolz                     0.11.0
dask                        2.30.0
dateutil                    2.8.2
decorator                   5.0.9
defusedxml                  0.7.1
docutils                    0.17.1
docx                        0.8.10
easydev                     0.11.0
entrypoints                 0.3
fbpca                       NA
google                      NA
gseapy                      0.10.4
h5py                        2.10.0
holoviews                   1.14.2
html5lib                    1.1
hvplot                      0.7.1
idna                        2.10
igraph                      0.9.6
import_ipynb                NA
intervaltree                NA
ipykernel                   5.3.4
ipython_genutils            0.2.0
ipywidgets                  7.6.3
jedi                        0.17.1
jinja2                      3.0.1
joblib                      1.0.1
json5                       NA
jsonschema                  3.2.0
jupyter_server              1.4.1
jupyterlab_pygments         0.1.2
jupyterlab_server           2.6.1
kiwisolver                  1.3.1
leidenalg                   0.8.2
llvmlite                    0.34.0
louvain                     0.7.0
lxml                        4.6.3
markupsafe                  2.0.1
matplotlib                  3.3.2
mistune                     0.8.4
mkl                         2.3.0
mpl_toolkits                NA
natsort                     7.1.1
nbclassic                   NA
nbclient                    0.5.3
nbconvert                   6.1.0
nbformat                    5.1.3
networkx                    2.6.2
numba                       0.51.2
numexpr                     2.7.3
numpy                       1.19.2
packaging                   21.0
pandas                      1.2.4
pandocfilters               NA
panel                       0.10.3
param                       1.11.1
parasail                    1.2.4
parso                       0.7.0
pexpect                     4.8.0
pickleshare                 0.7.5
pkg_resources               NA
prometheus_client           NA
prompt_toolkit              3.0.17
psutil                      5.8.0
ptyprocess                  0.7.0
pvectorc                    NA
pycparser                   2.20
pygments                    2.9.0
pylab                       NA
pynndescent                 0.5.2
pyparsing                   2.4.7
pyrsistent                  NA
pytoml                      NA
pytz                        2021.1
pyviz_comms                 2.0.2
requests                    2.25.1
requests_cache              0.5.2
scanorama                   1.7.1
scanpy                      1.8.1
scipy                       1.6.2
scirpy                      0.9.1
scvelo                      0.2.3
seaborn                     0.11.0
send2trash                  NA
setuptools_scm              NA
sinfo                       0.3.1
six                         1.16.0
sklearn                     0.24.2
sniffio                     1.2.0
socks                       1.7.1
sortedcontainers            2.4.0
soupsieve                   2.2.1
sphinxcontrib               NA
statsmodels                 0.12.2
storemagic                  NA
tables                      3.6.1
tblib                       1.7.0
testpath                    0.5.0
texttable                   1.6.4
threadpoolctl               2.2.0
tlz                         0.11.0
toolz                       0.11.1
tornado                     6.1
tqdm                        4.61.2
tracerlib                   NA
traitlets                   5.0.5
typing_extensions           NA
umap                        0.4.6
urllib3                     1.26.6
wcwidth                     0.2.5
webencodings                0.5.1
wrapt                       1.11.2
yaml                        5.4.1
yamlordereddictloader       NA
zipp                        NA
zmq                         20.0.0
zope                        NA
-----
IPython             7.22.0
jupyter_client      6.1.12
jupyter_core        4.7.1
jupyterlab          3.0.14
notebook            6.4.0
-----
Python 3.8.5 (default, Sep  4 2020, 07:30:14) [GCC 7.3.0]
Linux-3.10.0-514.2.2.el7.x86_64-x86_64-with-glibc2.10
352 logical CPU cores, x86_64
-----
Session information updated at 2021-11-25 10:09
```

In [3]:

```
# define sample metadata. Usually read from a file.
exclude_genes = ['RPL', 'RPS', 'TRAV', 'TRAJ', 'TRBJ', 'TRBV','MRP','FAU','DAP3','UBA52','IGHV', 'IGKV', 'IGLV']
antilist = [

]

In_path = '/user/ifrec/liuyuchen/scRNASeq_DATA/Yamazaki_lu_scRNASeq_all_pfizer/'

out_path = '/user/ifrec/liuyuchen/Analysis_Reports/Yamazaki_lu_scRNASeq_all_pfizer/'
```

In [4]:

```
samples = {}
for line in open(In_path+'sample_list.txt'):
    sample = line.strip()
    samples[sample]={}
    samples[sample]['gex']=sample
    samples[sample]['TCR']=sample.replace('5DE','TCR')
```

In [5]:

```
samples
```

Out[5]:

```
{'25_and_28_1_5DE': {'gex': '25_and_28_1_5DE', 'TCR': '25_and_28_1_TCR'},
 '25_and_28_2_5DE': {'gex': '25_and_28_2_5DE', 'TCR': '25_and_28_2_TCR'},
 '25_and_28_3_5DE': {'gex': '25_and_28_3_5DE', 'TCR': '25_and_28_3_TCR'},
 '25_and_28_4_5DE': {'gex': '25_and_28_4_5DE', 'TCR': '25_and_28_4_TCR'},
 '25_and_28_5_5DE': {'gex': '25_and_28_5_5DE', 'TCR': '25_and_28_5_TCR'},
 '25_and_28_6_5DE': {'gex': '25_and_28_6_5DE', 'TCR': '25_and_28_6_TCR'},
 '25_and_28_7_5DE': {'gex': '25_and_28_7_5DE', 'TCR': '25_and_28_7_TCR'},
 '25_and_28_8_5DE': {'gex': '25_and_28_8_5DE', 'TCR': '25_and_28_8_TCR'},
 '27_1_5DE': {'gex': '27_1_5DE', 'TCR': '27_1_TCR'},
 '27_2_5DE': {'gex': '27_2_5DE', 'TCR': '27_2_TCR'},
 '4_and_17_1_5DE': {'gex': '4_and_17_1_5DE', 'TCR': '4_and_17_1_TCR'},
 '4_and_17_2_5DE': {'gex': '4_and_17_2_5DE', 'TCR': '4_and_17_2_TCR'},
 '4_and_17_3_5DE': {'gex': '4_and_17_3_5DE', 'TCR': '4_and_17_3_TCR'},
 '4_and_17_4_5DE': {'gex': '4_and_17_4_5DE', 'TCR': '4_and_17_4_TCR'},
 '4_and_17_5_5DE': {'gex': '4_and_17_5_5DE', 'TCR': '4_and_17_5_TCR'},
 '4_and_17_6_5DE': {'gex': '4_and_17_6_5DE', 'TCR': '4_and_17_6_TCR'},
 '4_and_17_7_5DE': {'gex': '4_and_17_7_5DE', 'TCR': '4_and_17_7_TCR'},
 '4_and_17_8_5DE': {'gex': '4_and_17_8_5DE', 'TCR': '4_and_17_8_TCR'},
 '8and_13and_15_1_5DE': {'gex': '8and_13and_15_1_5DE',
  'TCR': '8and_13and_15_1_TCR'},
 '8and_13and_15_2_5DE': {'gex': '8and_13and_15_2_5DE',
  'TCR': '8and_13and_15_2_TCR'},
 '8and_13and_15_3_5DE': {'gex': '8and_13and_15_3_5DE',
  'TCR': '8and_13and_15_3_TCR'},
 '8and_13and_15_4_5DE': {'gex': '8and_13and_15_4_5DE',
  'TCR': '8and_13and_15_4_TCR'},
 '8and_13and_15_5_5DE': {'gex': '8and_13and_15_5_5DE',
  'TCR': '8and_13and_15_5_TCR'},
 '8and_13and_15_6_5DE': {'gex': '8and_13and_15_6_5DE',
  'TCR': '8and_13and_15_6_TCR'},
 '8and_13and_15_7_5DE': {'gex': '8and_13and_15_7_5DE',
  'TCR': '8and_13and_15_7_TCR'},
 '8and_13and_15_8_5DE': {'gex': '8and_13and_15_8_5DE',
  'TCR': '8and_13and_15_8_TCR'}}
```

In [6]:

```
adatalist = []
for sample, sample_meta in samples.items():
    gex_file = In_path+sample_meta["gex"]+'/outs/filtered_feature_bc_matrix.h5'
    adata = sc.read_10x_h5(gex_file, gex_only=False)
    tcr_file = In_path+sample_meta["TCR"]+'/outs/filtered_contig_annotations.csv'
    adata_tcr = ir.io.read_10x_vdj(tcr_file)
    ir.pp.merge_with_ir(adata, adata_tcr)
    adata.var_names_make_unique()
    adata.obs["Sample"] = sample
    hashtag = adata.var_names[adata.var_names.str.contains('Hashtag')].tolist()
    for h in hashtag:
        adata.obs[h] = adata[:, h].X.A
    adata.obs['Hashtag'] = adata.obs[hashtag].idxmax(axis=1)
    antis = adata.var_names[adata.var_names.str.contains('human')].tolist()
    for a in antis:
        adata.obs[a] = adata[:, a].X.A
        data = adata[:, a].X.A
        data =  np.interp(data, (data.min(), data.max()), (0, 10))
        adata.obs[a+'_normalized'] = data 
    adata = adata[:,adata.var[adata.var['feature_types']!='Antibody Capture'].index]
    adatalist.append(adata)
```

```
Variable names are not unique. To make them unique, call `.var_names_make_unique`.
... storing 'feature_types' as categorical
... storing 'genome' as categorical
Variable names are not unique. To make them unique, call `.var_names_make_unique`.
... storing 'feature_types' as categorical
... storing 'genome' as categorical
Variable names are not unique. To make them unique, call `.var_names_make_unique`.
... storing 'feature_types' as categorical
... storing 'genome' as categorical
Variable names are not unique. To make them unique, call `.var_names_make_unique`.
... storing 'feature_types' as categorical
... storing 'genome' as categorical
Variable names are not unique. To make them unique, call `.var_names_make_unique`.
... storing 'feature_types' as categorical
... storing 'genome' as categorical
Variable names are not unique. To make them unique, call `.var_names_make_unique`.
... storing 'feature_types' as categorical
... storing 'genome' as categorical
Variable names are not unique. To make them unique, call `.var_names_make_unique`.
... storing 'feature_types' as categorical
... storing 'genome' as categorical
Variable names are not unique. To make them unique, call `.var_names_make_unique`.
... storing 'feature_types' as categorical
... storing 'genome' as categorical
Variable names are not unique. To make them unique, call `.var_names_make_unique`.
... storing 'feature_types' as categorical
... storing 'genome' as categorical
Variable names are not unique. To make them unique, call `.var_names_make_unique`.
... storing 'feature_types' as categorical
... storing 'genome' as categorical
Variable names are not unique. To make them unique, call `.var_names_make_unique`.
... storing 'feature_types' as categorical
... storing 'genome' as categorical
Variable names are not unique. To make them unique, call `.var_names_make_unique`.
... storing 'feature_types' as categorical
... storing 'genome' as categorical
Variable names are not unique. To make them unique, call `.var_names_make_unique`.
... storing 'feature_types' as categorical
... storing 'genome' as categorical
Variable names are not unique. To make them unique, call `.var_names_make_unique`.
... storing 'feature_types' as categorical
... storing 'genome' as categorical
Variable names are not unique. To make them unique, call `.var_names_make_unique`.
... storing 'feature_types' as categorical
... storing 'genome' as categorical
Variable names are not unique. To make them unique, call `.var_names_make_unique`.
... storing 'feature_types' as categorical
... storing 'genome' as categorical
Variable names are not unique. To make them unique, call `.var_names_make_unique`.
... storing 'feature_types' as categorical
... storing 'genome' as categorical
Variable names are not unique. To make them unique, call `.var_names_make_unique`.
... storing 'feature_types' as categorical
... storing 'genome' as categorical
Variable names are not unique. To make them unique, call `.var_names_make_unique`.
... storing 'feature_types' as categorical
... storing 'genome' as categorical
Variable names are not unique. To make them unique, call `.var_names_make_unique`.
... storing 'feature_types' as categorical
... storing 'genome' as categorical
Variable names are not unique. To make them unique, call `.var_names_make_unique`.
... storing 'feature_types' as categorical
... storing 'genome' as categorical
Variable names are not unique. To make them unique, call `.var_names_make_unique`.
... storing 'feature_types' as categorical
... storing 'genome' as categorical
Variable names are not unique. To make them unique, call `.var_names_make_unique`.
... storing 'feature_types' as categorical
... storing 'genome' as categorical
Variable names are not unique. To make them unique, call `.var_names_make_unique`.
... storing 'feature_types' as categorical
... storing 'genome' as categorical
Variable names are not unique. To make them unique, call `.var_names_make_unique`.
... storing 'feature_types' as categorical
... storing 'genome' as categorical
Variable names are not unique. To make them unique, call `.var_names_make_unique`.
... storing 'feature_types' as categorical
... storing 'genome' as categorical
```

In [7]:

```
#sc.set_figure_params(scanpy=True, dpi=200,  figsize=[12.8,9.6])
sc.settings.verbosity = 3
```

In [8]:

```
adata = sc_pipe.unify_value(adatalist)
```

In [9]:

```
adata.obs['Sample'].value_counts()
```

Out[9]:

```
8and_13and_15_3_5DE    25160
8and_13and_15_6_5DE    24872
8and_13and_15_8_5DE    24789
8and_13and_15_4_5DE    23650
8and_13and_15_7_5DE    23401
8and_13and_15_5_5DE    23164
8and_13and_15_1_5DE    22018
8and_13and_15_2_5DE    20872
25_and_28_1_5DE        12018
25_and_28_2_5DE        11776
25_and_28_8_5DE        11718
25_and_28_7_5DE        11636
25_and_28_3_5DE        11076
25_and_28_5_5DE        10171
27_1_5DE               10171
4_and_17_4_5DE         10152
25_and_28_6_5DE        10151
25_and_28_4_5DE         9899
4_and_17_5_5DE          9484
4_and_17_3_5DE          9328
4_and_17_1_5DE          9120
4_and_17_2_5DE          8624
4_and_17_6_5DE          8582
4_and_17_8_5DE          8209
4_and_17_7_5DE          8006
27_2_5DE                6746
Name: Sample, dtype: int64
```

In [10]:

```
sc.pl.highest_expr_genes(adata)
```

```
normalizing counts per cell
    finished (0:00:08)
```

In [11]:

```
adata.obs['Hashtag'].value_counts()
```

Out[11]:

```
_8_3wks_Hashtag_1      44273
_15_6wks_Hashtag_8     38721
17_24wks_Hashtag_6     35986
_8_6wks_Hashtag_2      31020
_28_3wks_Hashtag_4     25259
_13_24wks_Hashtag_6    22591
_28_6wks_Hashtag_5     21037
_25_6wks_Hashtag_2     20554
17_6wks_Hashtag_5      20396
_13_3wks_Hashtag_4     19101
_25_24wks_Hashtag_3    14779
_8_24wks_Hashtag_3     11740
27_6wks_Hashtag_2      10775
_13_6wks_Hashtag_5      9022
_15_24wks_Hashtag_9     8325
_28_24wks_Hashtag_6     6136
27_24wks_Hashtag_3      5886
4_6wks_Hashtag_2        4964
4_24wks_Hashtag_3       3643
17_3wks_Hashtag_4       3322
4_3wks_Hashtag_1        3194
_15_3wks_Hashtag_7      3133
_25_3wks_Hashtag_1       680
27_3wks_Hashtag_1        256
Name: Hashtag, dtype: int64
```

In [12]:

```
hashtag = adata.obs['Hashtag'].unique().tolist()
```

In [13]:

```
tags ={x:x.split('_Hashtag_')[0] for x in hashtag}
```

In [14]:

```
hashtag
```

Out[14]:

```
['_25_6wks_Hashtag_2',
 '_28_24wks_Hashtag_6',
 '_28_6wks_Hashtag_5',
 '_25_24wks_Hashtag_3',
 '_28_3wks_Hashtag_4',
 '_25_3wks_Hashtag_1',
 '27_6wks_Hashtag_2',
 '27_24wks_Hashtag_3',
 '27_3wks_Hashtag_1',
 '17_24wks_Hashtag_6',
 '17_6wks_Hashtag_5',
 '4_3wks_Hashtag_1',
 '17_3wks_Hashtag_4',
 '4_6wks_Hashtag_2',
 '4_24wks_Hashtag_3',
 '_8_6wks_Hashtag_2',
 '_8_3wks_Hashtag_1',
 '_13_24wks_Hashtag_6',
 '_15_6wks_Hashtag_8',
 '_8_24wks_Hashtag_3',
 '_13_3wks_Hashtag_4',
 '_13_6wks_Hashtag_5',
 '_15_24wks_Hashtag_9',
 '_15_3wks_Hashtag_7']
```

In [15]:

```
adata.obs['Batch']=adata.obs['Sample']
```

In [16]:

```
adata.obs['Sample']=adata.obs['Hashtag'].map(tags)
```

In [17]:

```
adata.obs['Sample'].value_counts()
```

Out[17]:

```
_8_3wks      44273
_15_6wks     38721
17_24wks     35986
_8_6wks      31020
_28_3wks     25259
_13_24wks    22591
_28_6wks     21037
_25_6wks     20554
17_6wks      20396
_13_3wks     19101
_25_24wks    14779
_8_24wks     11740
27_6wks      10775
_13_6wks      9022
_15_24wks     8325
_28_24wks     6136
27_24wks      5886
4_6wks        4964
4_24wks       3643
17_3wks       3322
4_3wks        3194
_15_3wks      3133
_25_3wks       680
27_3wks        256
Name: Sample, dtype: int64
```

In [18]:

```
adata = sc_pipe.qc_and_preprocess(adata,out_path,multi_sample=True)
```

```
normalizing counts per cell
    finished (0:00:08)
WARNING: saving figure to file figures/highest_expr_genes_before_filter.png
```

```
Running Scrublet
filtered out 9560 genes that are detected in less than 3 cells
normalizing counts per cell
    finished (0:00:05)
extracting highly variable genes
    finished (0:03:17)
--> added
    'highly_variable', boolean vector (adata.var)
    'means', float vector (adata.var)
    'dispersions', float vector (adata.var)
    'dispersions_norm', float vector (adata.var)
normalizing counts per cell
    finished (0:00:00)
normalizing counts per cell
    finished (0:01:05)
Embedding transcriptomes using PCA...
Detected doublet rate = 0.3%
Estimated detectable doublet fraction = 24.0%
Overall doublet rate:
	Expected   = 5.0%
	Estimated  = 1.1%
    Scrublet finished (1:06:37)
```

```
... storing 'IR_VJ_1_c_call' as categorical
... storing 'IR_VJ_1_j_call' as categorical
... storing 'IR_VJ_2_j_call' as categorical
... storing 'IR_VDJ_2_j_call' as categorical
... storing 'IR_VJ_1_junction' as categorical
... storing 'IR_VJ_2_junction' as categorical
... storing 'IR_VDJ_1_junction' as categorical
... storing 'IR_VDJ_2_junction' as categorical
... storing 'IR_VJ_1_junction_aa' as categorical
... storing 'IR_VJ_2_junction_aa' as categorical
... storing 'IR_VDJ_1_junction_aa' as categorical
... storing 'IR_VDJ_2_junction_aa' as categorical
... storing 'IR_VJ_1_v_call' as categorical
... storing 'IR_VJ_2_v_call' as categorical
... storing 'IR_VDJ_1_v_call' as categorical
... storing 'IR_VDJ_2_v_call' as categorical
... storing 'Sample' as categorical
... storing 'Hashtag' as categorical
... storing 'Batch' as categorical
```

```
WARNING: saving figure to file figures/violin_QC_of_entire_set.pdf
```

```
WARNING: saving figure to file figures/violin_QC_by_sample_1.png
```

```
WARNING: saving figure to file figures/violin_QC_by_sample_2.png
```

```
WARNING: saving figure to file figures/violin_QC_by_sample_3.png
```

```
WARNING: saving figure to file figures/violin_QC_by_sample_4.png
```

```
WARNING: saving figure to file figures/violin_scrublet_sample.png
```

```
WARNING: saving figure to file figures/scatter_count_to_mito_1.png
```

```
WARNING: saving figure to file figures/scatter_count_to_gene.png
```

```
filtered out 0 cells that have less than 67 counts
filtered out 9560 genes that are detected in less than 3 cells
filtered out 9560 genes that are detected in less than 3 cells
filtered out 0 cells that have more than 41504487 counts
filtered out 13510 cells that has over 19% reads belong to mitochondrial genes
```

```
Trying to set attribute `.obs` of view, copying.
```

```
filtered out 0 cells that have less than 176 genes expressed
filtered out 9967 cells that have over 5784 genes expressed
WARNING: saving figure to file figures/violin_QC_of_entire_set_after_filtration.pdf
```

In [19]:

```
adata = sc_pipe.feature_selection(adata, out_path, exclude_genes)
#adata = adata[:, adata.var.highly_variable]
#adata = sc_pipe.clustering(adata,hash_samples)
adata = sc_pipe.BBKNN_clustering(adata,out_path, resol=1, multi_sample=True)
```

```
normalizing counts per cell
    finished (0:00:03)
normalizing counts per cell
    finished (0:00:07)
WARNING: saving figure to file figures/highest_expr_genes_after_QC.png
```

```
normalizing counts per cell
    finished (0:00:06)
WARNING: saving figure to file figures/highest_expr_genes_after_exclude.png
```

```
If you pass `n_top_genes`, all cutoffs are ignored.
extracting highly variable genes
    finished (0:00:08)
--> added
    'highly_variable', boolean vector (adata.var)
    'means', float vector (adata.var)
    'dispersions', float vector (adata.var)
    'dispersions_norm', float vector (adata.var)

 Number of highly variable genes: 4000
WARNING: saving figure to file figures/filter_genes_dispersion_highly_variable_genes.png
```

```
computing PCA
    on highly variable genes
    with n_comps=50
    finished (0:00:51)
computing neighbors
    using 'X_pca' with n_pcs = 50
    finished: added to `.uns['neighbors']`
    `.obsp['distances']`, distances for each pair of neighbors
    `.obsp['connectivities']`, weighted adjacency matrix (0:01:51)
computing UMAP
    finished: added
    'X_umap', UMAP coordinates (adata.obsm) (0:05:17)
computing batch balanced neighbors
	finished: added to `.uns['neighbors']`
	`.obsp['distances']`, distances for each pair of neighbors
	`.obsp['connectivities']`, weighted adjacency matrix (0:06:22)
running Leiden clustering
    finished: found 16 clusters and added
    'leiden', the cluster labels (adata.obs, categorical) (0:21:20)
running PAGA
    finished: added
    'paga/connectivities', connectivities adjacency (adata.uns)
    'paga/connectivities_tree', connectivities subtree (adata.uns) (0:00:45)
--> added 'pos', the PAGA positions (adata.uns['paga'])
WARNING: saving figure to file figures/paga_Graph.png
```

```
computing UMAP
    finished: added
    'X_umap', UMAP coordinates (adata.obsm) (0:06:28)
WARNING: saving figure to file figures/umap_Leiden_clustering.png
```

```
WARNING: saving figure to file figures/umap_no_bbknn.png
```

```
WARNING: saving figure to file figures/umap_by_sample.png
```

In [20]:

```
adata = sc_pipe.differential(adata,out_path, multi_sample = True)
```

```
WARNING: Default of the method has been changed to 't-test' from 't-test_overestim_var'
ranking genes
    finished: added to `.uns['rank_genes_groups']`
    'names', sorted np.recarray to be indexed by group ids
    'scores', sorted np.recarray to be indexed by group ids
    'logfoldchanges', sorted np.recarray to be indexed by group ids
    'pvals', sorted np.recarray to be indexed by group ids
    'pvals_adj', sorted np.recarray to be indexed by group ids (0:03:01)
WARNING: saving figure to file figures/rank_genes_groups_leiden_by_cluster.png
```

```
WARNING: saving figure to file figures/umap_top2_in_clusters.png
```

```
WARNING: dendrogram data not found (using key=dendrogram_leiden). Running `sc.tl.dendrogram` with default parameters. For fine tuning it is recommended to run `sc.tl.dendrogram` independently.
    using 'X_pca' with n_pcs = 50
Storing dendrogram info using `.uns['dendrogram_leiden']`
WARNING: saving figure to file figures/dotplot_in_clusters.png
```

```
WARNING: Gene labels are not shown when more than 50 genes are visualized. To show gene labels set `show_gene_labels=True`
WARNING: saving figure to file figures/heatmap_in_clusters.png
```

```
WARNING: saving figure to file figures/matrixplot_in_clusters.png
```

```
WARNING: saving figure to file figures/stacked_violin_in_clusters.png
```

```
WARNING: saving figure to file figures/tracksplot_in_clusters.png
```

```
WARNING: Default of the method has been changed to 't-test' from 't-test_overestim_var'
ranking genes
    finished: added to `.uns['rank_genes_samples']`
    'names', sorted np.recarray to be indexed by group ids
    'scores', sorted np.recarray to be indexed by group ids
    'logfoldchanges', sorted np.recarray to be indexed by group ids
    'pvals', sorted np.recarray to be indexed by group ids
    'pvals_adj', sorted np.recarray to be indexed by group ids (0:04:22)
WARNING: saving figure to file figures/rank_genes_groups_Sample_compare.png
```

In [21]:

```
ir.tl.chain_qc(adata)
```

In [22]:

```
ax = ir.pl.group_abundance(adata, groupby="receptor_subtype", target_col="Sample")
```

```
... storing 'receptor_type' as categorical
... storing 'receptor_subtype' as categorical
... storing 'chain_pairing' as categorical
```

In [23]:

```
ax = ir.pl.group_abundance(adata, groupby="chain_pairing", target_col="Sample")
```

In [24]:

```
# using default parameters, `ir_dist` will compute nucleotide sequence identity
ir.pp.ir_dist(adata)
ir.tl.define_clonotypes(adata, receptor_arms="all", dual_ir="primary_only")
```

```
Computing sequence x sequence distance matrix for VJ sequences.
Computing sequence x sequence distance matrix for VDJ sequences.
Initializing lookup tables. 
--> Done initializing lookup tables. (0:00:05)
Computing clonotype x clonotype distances.
NB: Computation happens in chunks. The progressbar only advances when a chunk has finished.
```

```
  0%|          | 0/34406 [00:00<?, ?it/s]
```

```
--> Done computing clonotype x clonotype distances.  (0:01:33)
Stored clonal assignments in `adata.obs["clone_id"]`.
```

In [25]:

```
ir.tl.clonal_expansion(adata)
```

In [26]:

```
sc.pl.umap(adata, color=["clonal_expansion", "clone_id_size"])
```

```
... storing 'clone_id' as categorical
... storing 'clonal_expansion' as categorical
```

In [27]:

```
ir.pl.clonal_expansion(adata, groupby="Sample", clip_at=4, normalize=False)
```

Out[27]:

```
<AxesSubplot:>
```

In [28]:

```
adata.obs
```

Out[28]:

|  | is\_cell | high\_confidence | multi\_chain | extra\_chains | IR\_VJ\_1\_c\_call | IR\_VJ\_2\_c\_call | IR\_VDJ\_1\_c\_call | IR\_VDJ\_2\_c\_call | IR\_VJ\_1\_consensus\_count | IR\_VJ\_2\_consensus\_count | ... | pct\_counts\_in\_top\_200\_genes | pct\_counts\_in\_top\_500\_genes | log\_counts | leiden | receptor\_type | receptor\_subtype | chain\_pairing | clone\_id | clone\_id\_size | clonal\_expansion |
| --- | --- | --- | --- | --- | --- | --- | --- | --- | --- | --- | --- | --- | --- | --- | --- | --- | --- | --- | --- | --- | --- |
| AAACCTGAGAGCTGCA-1-0 | True | True | False | [] | TRAC | NaN | TRBC1 | NaN | 5681.0 | NaN | ... | 44.478503 | 59.537845 | 9.451167 | 10 | TCR | TRA+TRB | single pair | 0 | 28.0 | >= 3 |
| AAACCTGAGCGTAGTG-1-0 | True | True | False | [] | TRAC | NaN | TRBC2 | NaN | 777.0 | NaN | ... | 47.661587 | 64.766159 | 8.244334 | 2 | TCR | TRA+TRB | single pair | 1 | 54.0 | >= 3 |
| AAACCTGAGCGTGAGT-1-0 | None | None | None | NaN | NaN | NaN | NaN | NaN | NaN | NaN | ... | 45.398517 | 60.137888 | 9.166702 | 9 | no IR | no IR | no IR | NaN | NaN | nan |
| AAACCTGAGCGTTTAC-1-0 | True | True | False | [] | TRAC | NaN | TRBC1 | NaN | 2828.0 | NaN | ... | 45.864066 | 59.704965 | 9.913487 | 6 | TCR | TRA+TRB | single pair | 2 | 8.0 | >= 3 |
| AAACCTGAGGACAGCT-1-0 | True | True | False | [] | TRAC | NaN | TRBC2 | NaN | 3331.0 | NaN | ... | 46.676249 | 62.468559 | 8.624432 | 7 | TCR | TRA+TRB | single pair | 3 | 4.0 | >= 3 |
| ... | ... | ... | ... | ... | ... | ... | ... | ... | ... | ... | ... | ... | ... | ... | ... | ... | ... | ... | ... | ... | ... |
| TTTGTCATCGGCGCAT-1-25 | None | None | None | NaN | NaN | NaN | NaN | NaN | NaN | NaN | ... | 53.904282 | 91.687657 | 6.677083 | 1 | no IR | no IR | no IR | NaN | NaN | nan |
| TTTGTCATCGTTGCCT-1-25 | None | None | None | NaN | NaN | NaN | NaN | NaN | NaN | NaN | ... | 57.100592 | 100.000000 | 6.516193 | 5 | no IR | no IR | no IR | NaN | NaN | nan |
| TTTGTCATCTCGATGA-1-25 | True | True | False | [] | TRAC | NaN | TRBC2 | NaN | 3682.0 | NaN | ... | 48.046595 | 63.172043 | 8.626944 | 8 | TCR | TRA+TRB | single pair | 23806 | 6059.0 | >= 3 |
| TTTGTCATCTGCGTAA-1-25 | None | None | None | NaN | NaN | NaN | NaN | NaN | NaN | NaN | ... | 52.649870 | 78.714162 | 7.048387 | 1 | no IR | no IR | no IR | NaN | NaN | nan |
| TTTGTCATCTGGAGCC-1-25 | None | None | None | NaN | NaN | NaN | NaN | NaN | NaN | NaN | ... | 55.387931 | 87.715517 | 6.833032 | 5 | no IR | no IR | no IR | NaN | NaN | nan |

341316 rows × 106 columns

In [29]:

```
sc.pl.umap(adata, color=["CD4", "CD8A"],use_raw=False,color_map= 'OrRd')
```

In [30]:

```
sc.pl.umap(adata, color=antis,use_raw=False,color_map= 'OrRd')
```

In [31]:

```
ad_ucsc=sc.read_h5ad('/user/ifrec/liuyuchen/Azimuth_ref/PBMC/Azimuth_ref.h5ad')
types = ['celltype.l1','celltype.l2','celltype.l3']

ad_ingest = adata[:, adata.var.highly_variable].copy()
var_names = ad_ucsc.var_names.intersection(ad_ingest.var_names)
ad_ucsc = ad_ucsc[:, var_names]
ad_ingest= ad_ingest[:, var_names]
sc.tl.ingest(ad_ingest, ad_ucsc, obs=types)
adata.obs  = ad_ingest.obs
sc.pl.umap(adata, color=types,legend_loc='on data')
```

```
running ingest
    finished (0:07:26)
```

In [32]:

```
ad_ucsc=sc.read_h5ad('/user/ifrec/liuyuchen/scRNASeq_DATA/COVID19/public_B_science/E-MTAB-10026/covid_portal_210320_with_raw.h5ad')
sc.pp.highly_variable_genes(ad_ucsc)
types = ['full_clustering']

sc.tl.pca(ad_ucsc)
sc.pp.neighbors(ad_ucsc)
ad_ingest = adata.copy()
var_names = ad_ucsc.var_names.intersection(ad_ingest.var_names)
ad_ucsc = ad_ucsc[:, var_names]
ad_ingest= ad_ingest[:, var_names]

sc.tl.ingest(ad_ingest, ad_ucsc, obs=types)
adata.obs  = ad_ingest.obs
sc.pl.umap(adata, color=types)
```

```
extracting highly variable genes
    finished (0:00:37)
--> added
    'highly_variable', boolean vector (adata.var)
    'means', float vector (adata.var)
    'dispersions', float vector (adata.var)
    'dispersions_norm', float vector (adata.var)
computing PCA
    on highly variable genes
    with n_comps=50
    finished (0:00:24)
computing neighbors
    using 'X_pca' with n_pcs = 50
    finished: added to `.uns['neighbors']`
    `.obsp['distances']`, distances for each pair of neighbors
    `.obsp['connectivities']`, weighted adjacency matrix (0:16:21)
running ingest
    finished (0:10:24)
```

In [33]:

```
adata.obs["Top_expanded_clone"]= 'Others'
adata.obs.loc[adata.obs["clone_id_size"]>100,"Top_expanded_clone"]=adata.obs.loc[adata.obs["clone_id_size"]>100,'clone_id']
```

In [34]:

```
#gene_list = pd.read_csv('/user/ifrec/liuyuchen/Analysis_Reports/Yamazaki_1108/gene_list.csv')
gene_list = pd.read_csv('/user/ifrec/liuyuchen/Analysis_Reports/Yamazaki_1108/Tfh_list_211110_2.csv')
#TFH_genes = gene_list['Genes upregulated in TFH cells'].dropna().tolist()
TFH_genes = gene_list['list1'].dropna().tolist()
feature_genes = adata.var_names.intersection(TFH_genes)
feature_genes
data = np.sum(adata[:, feature_genes].X, axis=1).A1 / len(feature_genes)
data = np.interp(data, (data.min(), data.max()), (0, 10))
adata.obs['Tfh_score_1'] = data
#TFH_genes = gene_list['Genes upregulated in TFH cells'].dropna().tolist()
TFH_genes = gene_list['list2'].dropna().tolist()
feature_genes = adata.var_names.intersection(TFH_genes)
feature_genes
data = np.sum(adata[:, feature_genes].X, axis=1).A1 / len(feature_genes)
data = np.interp(data, (data.min(), data.max()), (0, 10))
adata.obs['Tfh_score_2'] = data
```

In [35]:

```
sc.pl.violin(adata,keys='Tfh_score_1',groupby='leiden',rotation=90)
```

```
... storing 'Top_expanded_clone' as categorical
```

In [36]:

```
sc.pl.violin(adata,keys='Tfh_score_2',groupby='leiden',rotation=90)
```

In [37]:

```
adata.obs['time'] = adata.obs['Sample'].apply(lambda x: x.split('_')[1])
```

In [38]:

```
sc.pl.umap(adata, groups ='3wks' ,color='time')
```

```
... storing 'time' as categorical
```

In [39]:

```
adata.write(out_path+'/Merged_pfizer_tcells.h5ad')
```

In [43]:

```
adata.obs
```

Out[43]:

|  | is\_cell | high\_confidence | multi\_chain | extra\_chains | IR\_VJ\_1\_c\_call | IR\_VJ\_2\_c\_call | IR\_VDJ\_1\_c\_call | IR\_VDJ\_2\_c\_call | IR\_VJ\_1\_consensus\_count | IR\_VJ\_2\_consensus\_count | ... | clone\_id\_size | clonal\_expansion | celltype.l1 | celltype.l2 | celltype.l3 | full\_clustering | Top\_expanded\_clone | Tfh\_score\_1 | Tfh\_score\_2 | time |
| --- | --- | --- | --- | --- | --- | --- | --- | --- | --- | --- | --- | --- | --- | --- | --- | --- | --- | --- | --- | --- | --- |
| cell |  |  |  |  |  |  |  |  |  |  |  |  |  |  |  |  |  |  |  |  |  |
| AAACCTGAGAGCTGCA-1-0 | True | True | False | [] | TRAC | NaN | TRBC1 | NaN | 5681.0 | NaN | ... | 28.0 | >= 3 | CD8 T | CD8 TEM | CD8 TEM\_2 | CD8.TE | Others | 0.630983 | 0.561950 | 25 |
| AAACCTGAGCGTAGTG-1-0 | True | True | False | [] | TRAC | NaN | TRBC2 | NaN | 777.0 | NaN | ... | 54.0 | >= 3 | CD4 T | CD4 TCM | CD4 TCM\_1 | CD4.CM | Others | 0.938146 | 0.742752 | 28 |
| AAACCTGAGCGTGAGT-1-0 | None | None | None | NaN | NaN | NaN | NaN | NaN | NaN | NaN | ... | NaN | nan | other T | gdT | gdT\_1 | NK\_16hi | Others | 1.033337 | 0.873725 | 25 |
| AAACCTGAGCGTTTAC-1-0 | True | True | False | [] | TRAC | NaN | TRBC1 | NaN | 2828.0 | NaN | ... | 8.0 | >= 3 | CD4 T | CD4 TCM | CD4 TEM\_3 | CD4.EM | Others | 0.725423 | 0.745934 | 28 |
| AAACCTGAGGACAGCT-1-0 | True | True | False | [] | TRAC | NaN | TRBC2 | NaN | 3331.0 | NaN | ... | 4.0 | >= 3 | CD4 T | CD4 TCM | CD4 TEM\_3 | CD4.CM | Others | 0.353632 | 0.363630 | 28 |
| ... | ... | ... | ... | ... | ... | ... | ... | ... | ... | ... | ... | ... | ... | ... | ... | ... | ... | ... | ... | ... | ... |
| TTTGTCATCGGCGCAT-1-25 | None | None | None | NaN | NaN | NaN | NaN | NaN | NaN | NaN | ... | NaN | nan | CD4 T | CD4 TEM | CD4 TEM\_3 | CD4.Naive | Others | 0.497584 | 0.511653 | 8 |
| TTTGTCATCGTTGCCT-1-25 | None | None | None | NaN | NaN | NaN | NaN | NaN | NaN | NaN | ... | NaN | nan | CD4 T | CD4 TEM | CD4 TEM\_3 | CD4.Naive | Others | 0.000000 | 0.000000 | 8 |
| TTTGTCATCTCGATGA-1-25 | True | True | False | [] | TRAC | NaN | TRBC2 | NaN | 3682.0 | NaN | ... | 6059.0 | >= 3 | CD8 T | CD8 TEM | CD8 TEM\_4 | NK\_16hi | 23806 | 1.287892 | 1.324306 | 13 |
| TTTGTCATCTGCGTAA-1-25 | None | None | None | NaN | NaN | NaN | NaN | NaN | NaN | NaN | ... | NaN | nan | CD4 T | CD4 TEM | CD4 TEM\_3 | CD4.CM | Others | 0.765889 | 0.787544 | 13 |
| TTTGTCATCTGGAGCC-1-25 | None | None | None | NaN | NaN | NaN | NaN | NaN | NaN | NaN | ... | NaN | nan | CD4 T | CD4 TEM | CD4 TEM\_3 | CD4.Naive | Others | 0.772122 | 0.793953 | 8 |

341316 rows × 114 columns

In [ ]:
