## Supplementary material for "Early acquisition of S-specific Tfh clonotypes after SARS-CoV-2 vaccination is associated with the longevity of anti-S antibodies": Pre_vaccination_samples.html

Yamazaki\_Iu\_scRNASeq\_pre\_only


In [1]:

```
import import_ipynb
import Scanpy_functions_v03262021 as sc_pipe
import scvelo as scv
scv.logging.print_version()
import warnings
import scirpy as ir
import scanpy as sc
import numpy as np
import scipy as sp
import pandas as pd
import matplotlib.pyplot as plt
from matplotlib import rcParams
from matplotlib import colors
import seaborn as sb
import bbknn
import logging
from sklearn.mixture import GaussianMixture
from scipy.stats     import norm
import glob
import os
import hvplot.pandas
import docx
from docx import Document
from docx.shared import Inches
from docx.shared import Pt
from scipy import sparse
import scanpy.external as sce
import holoviews as hv
import panel as pn
import bokeh
from bokeh.resources import INLINE
import scanorama
import gseapy
```

```
importing Jupyter notebook from Scanpy_functions_v03262021.ipynb
Running scvelo 0.2.3 (python 3.8.5) on 2021-12-10 14:24.
Running scvelo 0.2.3 (python 3.8.5) on 2021-12-10 14:24.
```

In [2]:

```
sc.logging.print_versions()
```

```
WARNING: If you miss a compact list, please try `print_header`!
-----
anndata     0.7.6
scanpy      1.8.1
sinfo       0.3.1
-----
Levenshtein                 NA
PIL                         8.3.1
Scanpy_functions_v03262021  NA
adjustText                  NA
airr                        1.3.1
anndata                     0.7.6
annoy                       NA
anyio                       NA
appdirs                     1.4.4
async_generator             1.10
attr                        21.2.0
babel                       2.9.1
backcall                    0.2.0
bbknn                       NA
bioservices                 1.7.11
bokeh                       2.3.3
bottleneck                  1.3.2
brotli                      NA
bs4                         4.9.3
cairo                       1.19.1
certifi                     2021.05.30
cffi                        1.14.6
chardet                     4.0.0
cloudpickle                 1.6.0
colorama                    0.4.4
colorcet                    1.0.0
colorlog                    NA
cycler                      0.10.0
cython_runtime              NA
cytoolz                     0.11.0
dask                        2.30.0
dateutil                    2.8.2
decorator                   5.0.9
defusedxml                  0.7.1
docutils                    0.17.1
docx                        0.8.10
easydev                     0.11.0
entrypoints                 0.3
fbpca                       NA
google                      NA
gseapy                      0.10.4
h5py                        3.6.0
holoviews                   1.14.2
html5lib                    1.1
hvplot                      0.7.1
idna                        2.10
igraph                      0.9.6
import_ipynb                NA
intervaltree                NA
ipykernel                   5.3.4
ipython_genutils            0.2.0
ipywidgets                  7.6.3
jedi                        0.17.1
jinja2                      3.0.1
joblib                      1.0.1
json5                       NA
jsonschema                  3.2.0
jupyter_server              1.4.1
jupyterlab_pygments         0.1.2
jupyterlab_server           2.6.1
kiwisolver                  1.3.1
leidenalg                   0.8.2
llvmlite                    0.34.0
louvain                     0.7.0
lxml                        4.6.3
markupsafe                  2.0.1
matplotlib                  3.3.2
mistune                     0.8.4
mkl                         2.3.0
mpl_toolkits                NA
natsort                     7.1.1
nbclassic                   NA
nbclient                    0.5.3
nbconvert                   6.1.0
nbformat                    5.1.3
networkx                    2.6.2
numba                       0.51.2
numexpr                     2.7.3
numpy                       1.19.2
packaging                   21.0
pandas                      1.2.4
pandocfilters               NA
panel                       0.10.3
param                       1.11.1
parasail                    1.2.4
parso                       0.7.0
pexpect                     4.8.0
pickleshare                 0.7.5
pkg_resources               NA
prometheus_client           NA
prompt_toolkit              3.0.17
psutil                      5.8.0
ptyprocess                  0.7.0
pvectorc                    NA
pycparser                   2.20
pygments                    2.9.0
pylab                       NA
pynndescent                 0.5.2
pyparsing                   2.4.7
pyrsistent                  NA
pytoml                      NA
pytz                        2021.1
pyviz_comms                 2.0.2
requests                    2.25.1
requests_cache              0.5.2
scanorama                   1.7.1
scanpy                      1.8.1
scipy                       1.6.2
scirpy                      0.9.1
scvelo                      0.2.3
seaborn                     0.11.0
send2trash                  NA
setuptools_scm              NA
sinfo                       0.3.1
six                         1.16.0
sklearn                     0.24.2
sniffio                     1.2.0
socks                       1.7.1
sortedcontainers            2.4.0
soupsieve                   2.2.1
sphinxcontrib               NA
statsmodels                 0.12.2
storemagic                  NA
tables                      3.6.1
tblib                       1.7.0
testpath                    0.5.0
texttable                   1.6.4
threadpoolctl               2.2.0
tlz                         0.11.0
toolz                       0.11.1
tornado                     6.1
tqdm                        4.61.2
tracerlib                   NA
traitlets                   5.0.5
typing_extensions           NA
umap                        0.4.6
urllib3                     1.26.6
wcwidth                     0.2.5
webencodings                0.5.1
wrapt                       1.11.2
yaml                        5.4.1
yamlordereddictloader       NA
zipp                        NA
zmq                         20.0.0
zope                        NA
-----
IPython             7.22.0
jupyter_client      6.1.12
jupyter_core        4.7.1
jupyterlab          3.0.14
notebook            6.4.0
-----
Python 3.8.5 (default, Sep  4 2020, 07:30:14) [GCC 7.3.0]
Linux-3.10.0-514.2.2.el7.x86_64-x86_64-with-glibc2.10
352 logical CPU cores, x86_64
-----
Session information updated at 2021-12-10 14:24
```

In [3]:

```
# define sample metadata. Usually read from a file.
exclude_genes = ['RPL', 'RPS', 'TRAV', 'TRAJ', 'TRBJ', 'TRBV','MRP','FAU','DAP3','UBA52','IGHV', 'IGKV', 'IGLV']
antilist = [

]

In_path = '/user/ifrec/liuyuchen/scRNASeq_DATA/Yamazaki_lu_scRNASeq_all_pfizer/'

out_path = '/user/ifrec/liuyuchen/Analysis_Reports/Yamazaki_lu_scRNASeq_pre/'
```

In [4]:

```
samples = {}
for line in open(In_path+'pre_list.txt'):
    sample = line.strip()
    samples[sample]={}
    samples[sample]['gex']=sample
    samples[sample]['TCR']=sample.replace('5DE','TCR')
```

In [5]:

```
samples
```

Out[5]:

```
{'Pre-1_5DE': {'gex': 'Pre-1_5DE', 'TCR': 'Pre-1_TCR'},
 'Pre-2_5DE': {'gex': 'Pre-2_5DE', 'TCR': 'Pre-2_TCR'},
 'Pre-3_5DE': {'gex': 'Pre-3_5DE', 'TCR': 'Pre-3_TCR'}}
```

In [6]:

```
adatalist = []
for sample, sample_meta in samples.items():
    gex_file = In_path+sample_meta["gex"]+'/outs/filtered_feature_bc_matrix.h5'
    adata = sc.read_10x_h5(gex_file, gex_only=False)
    tcr_file = In_path+sample_meta["TCR"]+'/outs/filtered_contig_annotations.csv'
    adata_tcr = ir.io.read_10x_vdj(tcr_file)
    ir.pp.merge_with_ir(adata, adata_tcr)
    adata.var_names_make_unique()
    adata.obs["Sample"] = sample
    hashtag = adata.var_names[adata.var_names.str.contains('Hashtag')].tolist()
    for h in hashtag:
        adata.obs[h] = adata[:, h].X.A
    adata.obs['Hashtag'] = adata.obs[hashtag].idxmax(axis=1)
    antis = adata.var_names[adata.var_names.str.contains('human')].tolist()
    for a in antis:
        adata.obs[a] = adata[:, a].X.A
        data = adata[:, a].X.A
        data =  np.interp(data, (data.min(), data.max()), (0, 10))
        adata.obs[a+'_normalized'] = data 
    adata = adata[:,adata.var[adata.var['feature_types']!='Antibody Capture'].index]
    adatalist.append(adata)
```

```
Variable names are not unique. To make them unique, call `.var_names_make_unique`.
... storing 'feature_types' as categorical
... storing 'genome' as categorical
Variable names are not unique. To make them unique, call `.var_names_make_unique`.
... storing 'feature_types' as categorical
... storing 'genome' as categorical
Variable names are not unique. To make them unique, call `.var_names_make_unique`.
... storing 'feature_types' as categorical
... storing 'genome' as categorical
```

In [7]:

```
#sc.set_figure_params(scanpy=True, dpi=200,  figsize=[12.8,9.6])
sc.settings.verbosity = 3
```

In [8]:

```
adata = sc_pipe.unify_value(adatalist)
```

In [9]:

```
adata.obs['Sample'].value_counts()
```

Out[9]:

```
Pre-3_5DE    9212
Pre-2_5DE    9108
Pre-1_5DE    8951
Name: Sample, dtype: int64
```

In [10]:

```
sc.pl.highest_expr_genes(adata)
```

```
normalizing counts per cell
    finished (0:00:00)
```

In [11]:

```
adata.obs['Hashtag'].value_counts()
```

Out[11]:

```
_15_pre_Hashtag_9     11034
_8_pre_Hashtag_7       5227
_13_pre_Hashtag_8      4016
_4_pre_Hashtag_6       3991
_17_pre_Hashtag_10     3003
Name: Hashtag, dtype: int64
```

In [12]:

```
for c in adata.obs.columns:
    if str(c).startswith('_'):
        adata.obs[c.strip('_')]=adata.obs[c]
        del adata.obs[c]
adata.obs
```

Out[12]:

|  | is\_cell | high\_confidence | multi\_chain | extra\_chains | IR\_VJ\_1\_c\_call | IR\_VJ\_2\_c\_call | IR\_VDJ\_1\_c\_call | IR\_VDJ\_2\_c\_call | IR\_VJ\_1\_consensus\_count | IR\_VJ\_2\_consensus\_count | ... | human\_CD4 | human\_CD4\_normalized | human\_CD8 | human\_CD8\_normalized | batch | 4\_pre\_Hashtag\_6 | 8\_pre\_Hashtag\_7 | 13\_pre\_Hashtag\_8 | 15\_pre\_Hashtag\_9 | 17\_pre\_Hashtag\_10 |
| --- | --- | --- | --- | --- | --- | --- | --- | --- | --- | --- | --- | --- | --- | --- | --- | --- | --- | --- | --- | --- | --- |
| AAACCTGAGATCCCAT-1-0 | True | True | False | [] | TRAC | NaN | TRBC1 | NaN | 1904.0 | NaN | ... | 357.0 | 0.876074 | 36.0 | 0.033664 | 0 | 3.0 | 3.0 | 4.0 | 374.0 | 1.0 |
| AAACCTGAGTCAATAG-1-0 | True | True | False | [] | TRAC | NaN | TRBC2 | NaN | 5323.0 | NaN | ... | 391.0 | 0.959509 | 3.0 | 0.002805 | 0 | 2.0 | 13.0 | 2448.0 | 8.0 | 7.0 |
| AAACCTGCATGGGACA-1-0 | True | True | False | [] | TRAC | NaN | TRBC1 | NaN | 1121.0 | NaN | ... | 359.0 | 0.880982 | 3.0 | 0.002805 | 0 | 11.0 | 313.0 | 8.0 | 7.0 | 4.0 |
| AAACCTGGTGTGAAAT-1-0 | True | True | False | [] | TRAC | NaN | TRBC1 | NaN | 768.0 | NaN | ... | 299.0 | 0.733742 | 17.0 | 0.015897 | 0 | 4.0 | 18.0 | 8.0 | 420.0 | 6.0 |
| AAACCTGTCACATAGC-1-0 | None | None | None | NaN | NaN | NaN | NaN | NaN | NaN | NaN | ... | 247.0 | 0.606135 | 7.0 | 0.006546 | 0 | 6.0 | 7.0 | 3.0 | 630.0 | 5.0 |
| ... | ... | ... | ... | ... | ... | ... | ... | ... | ... | ... | ... | ... | ... | ... | ... | ... | ... | ... | ... | ... | ... |
| TTTGTCAGTTGTTTGG-1-2 | None | None | None | NaN | NaN | NaN | NaN | NaN | NaN | NaN | ... | 407.0 | 0.209848 | 18.0 | 0.016648 | 2 | 9.0 | 11.0 | 6.0 | 3474.0 | 8.0 |
| TTTGTCATCAGAGACG-1-2 | True | True | False | [] | TRAC | NaN | TRBC2 | NaN | 981.0 | NaN | ... | 320.0 | 0.164991 | 6.0 | 0.005549 | 2 | 3.0 | 3.0 | 8.0 | 127.0 | 6.0 |
| TTTGTCATCCACGAAT-1-2 | True | True | False | [] | TRAC | NaN | TRBC2 | NaN | 3114.0 | NaN | ... | 796.0 | 0.410415 | 9.0 | 0.008324 | 2 | 6.0 | 5.0 | 3086.0 | 7.0 | 2.0 |
| TTTGTCATCCTGCAGG-1-2 | True | True | False | [] | TRAC | NaN | TRBC2 | NaN | 406.0 | NaN | ... | 642.0 | 0.331013 | 2.0 | 0.001850 | 2 | 7.0 | 4314.0 | 16.0 | 8.0 | 8.0 |
| TTTGTCATCTGCGGCA-1-2 | None | None | None | NaN | NaN | NaN | NaN | NaN | NaN | NaN | ... | 862.0 | 0.444444 | 24.0 | 0.022198 | 2 | 1735.0 | 6.0 | 14.0 | 494.0 | 5.0 |

27271 rows × 63 columns

In [13]:

```
hashtag = adata.obs['Hashtag'].unique().tolist()
```

In [14]:

```
tags ={x:x.split('_Hashtag_')[0] for x in hashtag}
```

In [15]:

```
hashtag
```

Out[15]:

```
['_15_pre_Hashtag_9',
 '_13_pre_Hashtag_8',
 '_8_pre_Hashtag_7',
 '_4_pre_Hashtag_6',
 '_17_pre_Hashtag_10']
```

In [16]:

```
adata.obs['Batch']=adata.obs['Sample']
```

In [17]:

```
adata.obs['Sample']=adata.obs['Hashtag'].map(tags)
```

In [18]:

```
adata.obs['Sample'].value_counts()
```

Out[18]:

```
_15_pre    11034
_8_pre      5227
_13_pre     4016
_4_pre      3991
_17_pre     3003
Name: Sample, dtype: int64
```

In [19]:

```
adata.obs['Sample']=adata.obs['Sample'].str.strip('_')
```

In [20]:

```
adata.obs['Hashtag']=adata.obs['Hashtag'].str.strip('_')
```

In [21]:

```
adata.obs['Sample'].value_counts()
```

Out[21]:

```
15_pre    11034
8_pre      5227
13_pre     4016
4_pre      3991
17_pre     3003
Name: Sample, dtype: int64
```

In [22]:

```
adata.obs['Hashtag'].value_counts()
```

Out[22]:

```
15_pre_Hashtag_9     11034
8_pre_Hashtag_7       5227
13_pre_Hashtag_8      4016
4_pre_Hashtag_6       3991
17_pre_Hashtag_10     3003
Name: Hashtag, dtype: int64
```

In [23]:

```
adata.obs['time'] = adata.obs['Sample'].apply(lambda x: x.split('_')[1])
```

In [24]:

```
adata.obs['time'].value_counts()
```

Out[24]:

```
pre    27271
Name: time, dtype: int64
```

In [25]:

```
adata = sc_pipe.qc_and_preprocess(adata,out_path,multi_sample=True)
```

```
normalizing counts per cell
    finished (0:00:00)
WARNING: saving figure to file figures/highest_expr_genes_before_filter.png
```

```
Running Scrublet
filtered out 14105 genes that are detected in less than 3 cells
normalizing counts per cell
    finished (0:00:00)
extracting highly variable genes
    finished (0:00:02)
--> added
    'highly_variable', boolean vector (adata.var)
    'means', float vector (adata.var)
    'dispersions', float vector (adata.var)
    'dispersions_norm', float vector (adata.var)
normalizing counts per cell
    finished (0:00:00)
normalizing counts per cell
    finished (0:00:00)
Embedding transcriptomes using PCA...
Detected doublet rate = 0.1%
Estimated detectable doublet fraction = 3.2%
Overall doublet rate:
	Expected   = 5.0%
	Estimated  = 2.2%
    Scrublet finished (0:01:01)
```

```
... storing 'IR_VJ_2_j_call' as categorical
... storing 'IR_VDJ_2_j_call' as categorical
... storing 'IR_VJ_1_junction' as categorical
... storing 'IR_VJ_2_junction' as categorical
... storing 'IR_VDJ_1_junction' as categorical
... storing 'IR_VDJ_2_junction' as categorical
... storing 'IR_VJ_1_junction_aa' as categorical
... storing 'IR_VJ_2_junction_aa' as categorical
... storing 'IR_VDJ_1_junction_aa' as categorical
... storing 'IR_VDJ_2_junction_aa' as categorical
... storing 'IR_VJ_2_v_call' as categorical
... storing 'IR_VDJ_1_v_call' as categorical
... storing 'IR_VDJ_2_v_call' as categorical
... storing 'Sample' as categorical
... storing 'Hashtag' as categorical
... storing 'Batch' as categorical
... storing 'time' as categorical
```

```
WARNING: saving figure to file figures/violin_QC_of_entire_set.pdf
```

```
WARNING: saving figure to file figures/violin_QC_by_sample_1.png
```

```
WARNING: saving figure to file figures/violin_QC_by_sample_2.png
```

```
WARNING: saving figure to file figures/violin_QC_by_sample_3.png
```

```
WARNING: saving figure to file figures/violin_QC_by_sample_4.png
```

```
WARNING: saving figure to file figures/violin_scrublet_sample.png
```

```
WARNING: saving figure to file figures/scatter_count_to_mito_1.png
```

```
WARNING: saving figure to file figures/scatter_count_to_gene.png
```

```
filtered out 0 cells that have less than 491 counts
filtered out 14105 genes that are detected in less than 3 cells
filtered out 14105 genes that are detected in less than 3 cells
filtered out 0 cells that have more than 54850411 counts
filtered out 297 cells that has over 13% reads belong to mitochondrial genes
filtered out 167 cells that have less than 591 genes expressed
```

```
Trying to set attribute `.obs` of view, copying.
```

```
filtered out 167 cells that have less than 591 genes expressed
filtered out 918 cells that have over 5000 genes expressed
WARNING: saving figure to file figures/violin_QC_of_entire_set_after_filtration.pdf
```

In [26]:

```
adata = sc_pipe.feature_selection(adata, out_path, exclude_genes)
#adata = adata[:, adata.var.highly_variable]
#adata = sc_pipe.clustering(adata,hash_samples)
adata = sc_pipe.BBKNN_clustering(adata,out_path, resol=1, multi_sample=True)
```

```
normalizing counts per cell
    finished (0:00:00)
normalizing counts per cell
    finished (0:00:00)
WARNING: saving figure to file figures/highest_expr_genes_after_QC.png
```

```
normalizing counts per cell
    finished (0:00:00)
WARNING: saving figure to file figures/highest_expr_genes_after_exclude.png
```

```
If you pass `n_top_genes`, all cutoffs are ignored.
extracting highly variable genes
    finished (0:00:00)
--> added
    'highly_variable', boolean vector (adata.var)
    'means', float vector (adata.var)
    'dispersions', float vector (adata.var)
    'dispersions_norm', float vector (adata.var)

 Number of highly variable genes: 4000
WARNING: saving figure to file figures/filter_genes_dispersion_highly_variable_genes.png
```

```
computing PCA
    on highly variable genes
    with n_comps=50
    finished (0:00:06)
computing neighbors
    using 'X_pca' with n_pcs = 50
    finished: added to `.uns['neighbors']`
    `.obsp['distances']`, distances for each pair of neighbors
    `.obsp['connectivities']`, weighted adjacency matrix (0:00:26)
computing UMAP
    finished: added
    'X_umap', UMAP coordinates (adata.obsm) (0:00:16)
computing batch balanced neighbors
	finished: added to `.uns['neighbors']`
	`.obsp['distances']`, distances for each pair of neighbors
	`.obsp['connectivities']`, weighted adjacency matrix (0:00:06)
running Leiden clustering
    finished: found 20 clusters and added
    'leiden', the cluster labels (adata.obs, categorical) (0:00:06)
running PAGA
    finished: added
    'paga/connectivities', connectivities adjacency (adata.uns)
    'paga/connectivities_tree', connectivities subtree (adata.uns) (0:00:01)
--> added 'pos', the PAGA positions (adata.uns['paga'])
WARNING: saving figure to file figures/paga_Graph.png
```

```
computing UMAP
    finished: added
    'X_umap', UMAP coordinates (adata.obsm) (0:00:15)
WARNING: saving figure to file figures/umap_Leiden_clustering.png
```

```
WARNING: saving figure to file figures/umap_no_bbknn.png
```

```
WARNING: saving figure to file figures/umap_by_sample.png
```

In [27]:

```
adata = sc_pipe.differential(adata,out_path, multi_sample = True)
```

```
WARNING: Default of the method has been changed to 't-test' from 't-test_overestim_var'
ranking genes
    finished: added to `.uns['rank_genes_groups']`
    'names', sorted np.recarray to be indexed by group ids
    'scores', sorted np.recarray to be indexed by group ids
    'logfoldchanges', sorted np.recarray to be indexed by group ids
    'pvals', sorted np.recarray to be indexed by group ids
    'pvals_adj', sorted np.recarray to be indexed by group ids (0:00:21)
WARNING: saving figure to file figures/rank_genes_groups_leiden_by_cluster.png
```

```
WARNING: saving figure to file figures/umap_top2_in_clusters.png
```

```
WARNING: dendrogram data not found (using key=dendrogram_leiden). Running `sc.tl.dendrogram` with default parameters. For fine tuning it is recommended to run `sc.tl.dendrogram` independently.
    using 'X_pca' with n_pcs = 50
Storing dendrogram info using `.uns['dendrogram_leiden']`
WARNING: saving figure to file figures/dotplot_in_clusters.png
```

```
WARNING: Gene labels are not shown when more than 50 genes are visualized. To show gene labels set `show_gene_labels=True`
WARNING: saving figure to file figures/heatmap_in_clusters.png
```

```
WARNING: saving figure to file figures/matrixplot_in_clusters.png
```

```
WARNING: saving figure to file figures/stacked_violin_in_clusters.png
```

```
WARNING: saving figure to file figures/tracksplot_in_clusters.png
```

```
WARNING: Default of the method has been changed to 't-test' from 't-test_overestim_var'
ranking genes
    finished: added to `.uns['rank_genes_samples']`
    'names', sorted np.recarray to be indexed by group ids
    'scores', sorted np.recarray to be indexed by group ids
    'logfoldchanges', sorted np.recarray to be indexed by group ids
    'pvals', sorted np.recarray to be indexed by group ids
    'pvals_adj', sorted np.recarray to be indexed by group ids (0:00:06)
WARNING: saving figure to file figures/rank_genes_groups_Sample_compare.png
```

In [28]:

```
ir.tl.chain_qc(adata)
```

In [29]:

```
ax = ir.pl.group_abundance(adata, groupby="receptor_subtype", target_col="Sample")
```

```
... storing 'receptor_type' as categorical
... storing 'receptor_subtype' as categorical
... storing 'chain_pairing' as categorical
```

In [30]:

```
ax = ir.pl.group_abundance(adata, groupby="chain_pairing", target_col="Sample")
```

In [31]:

```
# using default parameters, `ir_dist` will compute nucleotide sequence identity
ir.pp.ir_dist(adata)
ir.tl.define_clonotypes(adata, receptor_arms="all", dual_ir="primary_only")
```

```
Computing sequence x sequence distance matrix for VJ sequences.
Computing sequence x sequence distance matrix for VDJ sequences.
Initializing lookup tables. 
--> Done initializing lookup tables. (0:00:00)
Computing clonotype x clonotype distances.
NB: Computation happens in chunks. The progressbar only advances when a chunk has finished.
```

```
  0%|          | 0/5511 [00:00<?, ?it/s]
```

```
--> Done computing clonotype x clonotype distances.  (0:00:16)
Stored clonal assignments in `adata.obs["clone_id"]`.
```

In [32]:

```
ir.tl.clonal_expansion(adata)
```

In [33]:

```
sc.pl.umap(adata, color=["clonal_expansion", "clone_id_size"])
```

```
... storing 'clone_id' as categorical
... storing 'clonal_expansion' as categorical
```

In [34]:

```
ir.pl.clonal_expansion(adata, groupby="Sample", clip_at=4, normalize=False)
```

Out[34]:

```
<AxesSubplot:>
```

In [35]:

```
adata.obs
```

Out[35]:

|  | is\_cell | high\_confidence | multi\_chain | extra\_chains | IR\_VJ\_1\_c\_call | IR\_VJ\_2\_c\_call | IR\_VDJ\_1\_c\_call | IR\_VDJ\_2\_c\_call | IR\_VJ\_1\_consensus\_count | IR\_VJ\_2\_consensus\_count | ... | pct\_counts\_in\_top\_200\_genes | pct\_counts\_in\_top\_500\_genes | log\_counts | leiden | receptor\_type | receptor\_subtype | chain\_pairing | clone\_id | clone\_id\_size | clonal\_expansion |
| --- | --- | --- | --- | --- | --- | --- | --- | --- | --- | --- | --- | --- | --- | --- | --- | --- | --- | --- | --- | --- | --- |
| AAACCTGAGATCCCAT-1-0 | True | True | False | [] | TRAC | NaN | TRBC1 | NaN | 1904.0 | NaN | ... | 55.138640 | 69.292316 | 8.888618 | 1 | TCR | TRA+TRB | single pair | 0 | 404.0 | >= 3 |
| AAACCTGAGTCAATAG-1-0 | True | True | False | [] | TRAC | NaN | TRBC2 | NaN | 5323.0 | NaN | ... | 49.884423 | 62.276872 | 9.653358 | 15 | TCR | TRA+TRB | single pair | 1 | 7.0 | >= 3 |
| AAACCTGCATGGGACA-1-0 | True | True | False | [] | TRAC | NaN | TRBC1 | NaN | 1121.0 | NaN | ... | 72.374302 | 83.701117 | 8.876266 | 6 | TCR | TRA+TRB | single pair | 2 | 69.0 | >= 3 |
| AAACCTGGTGTGAAAT-1-0 | True | True | False | [] | TRAC | NaN | TRBC1 | NaN | 768.0 | NaN | ... | 52.491620 | 65.921788 | 9.099409 | 1 | TCR | TRA+TRB | single pair | 3 | 78.0 | >= 3 |
| AAACCTGTCACATAGC-1-0 | None | None | None | NaN | NaN | NaN | NaN | NaN | NaN | NaN | ... | 52.979817 | 65.860625 | 9.259511 | 11 | no IR | no IR | no IR | NaN | NaN | nan |
| ... | ... | ... | ... | ... | ... | ... | ... | ... | ... | ... | ... | ... | ... | ... | ... | ... | ... | ... | ... | ... | ... |
| TTTGTCAGTTCCACTC-1-2 | True | True | False | [] | TRAC | NaN | TRBC2 | NaN | 304.0 | NaN | ... | 50.311382 | 63.786935 | 8.928905 | 3 | TCR | TRA+TRB | single pair | 18 | 88.0 | >= 3 |
| TTTGTCATCAGAGACG-1-2 | True | True | False | [] | TRAC | NaN | TRBC2 | NaN | 981.0 | NaN | ... | 63.110902 | 75.861529 | 8.761550 | 0 | TCR | TRA+TRB | single pair | 2535 | 5.0 | >= 3 |
| TTTGTCATCCACGAAT-1-2 | True | True | False | [] | TRAC | NaN | TRBC2 | NaN | 3114.0 | NaN | ... | 50.734882 | 63.816897 | 9.095266 | 3 | TCR | TRA+TRB | single pair | 157 | 91.0 | >= 3 |
| TTTGTCATCCTGCAGG-1-2 | True | True | False | [] | TRAC | NaN | TRBC2 | NaN | 406.0 | NaN | ... | 62.047527 | 75.366123 | 8.887100 | 7 | TCR | TRA+TRB | single pair | 310 | 4.0 | >= 3 |
| TTTGTCATCTGCGGCA-1-2 | None | None | None | NaN | NaN | NaN | NaN | NaN | NaN | NaN | ... | 50.532817 | 62.193715 | 9.597845 | 3 | no IR | no IR | no IR | NaN | NaN | nan |

25889 rows × 88 columns

In [36]:

```
sc.pl.umap(adata, color=["CD4", "CD8A"],use_raw=False,color_map= 'OrRd')
```

In [37]:

```
sc.pl.umap(adata, color=antis,use_raw=False,color_map= 'OrRd')
```

In [38]:

```
ad_ucsc=sc.read_h5ad('/user/ifrec/liuyuchen/Azimuth_ref/PBMC/Azimuth_ref.h5ad')
types = ['celltype.l1','celltype.l2','celltype.l3']

ad_ingest = adata[:, adata.var.highly_variable].copy()
var_names = ad_ucsc.var_names.intersection(ad_ingest.var_names)
ad_ucsc = ad_ucsc[:, var_names]
ad_ingest= ad_ingest[:, var_names]
sc.tl.ingest(ad_ingest, ad_ucsc, obs=types)
adata.obs  = ad_ingest.obs
sc.pl.umap(adata, color=types,legend_loc='on data')
```

```
running ingest
    finished (0:00:44)
```

In [39]:

```
adata.obs["Top_expanded_clone"]= 'Others'
adata.obs.loc[adata.obs["clone_id_size"]>100,"Top_expanded_clone"]=adata.obs.loc[adata.obs["clone_id_size"]>100,'clone_id']
```

In [40]:

```
#gene_list = pd.read_csv('/user/ifrec/liuyuchen/Analysis_Reports/Yamazaki_1108/gene_list.csv')
gene_list = pd.read_csv('/user/ifrec/liuyuchen/Analysis_Reports/Yamazaki_1108/Tfh_list_211110_2.csv')
#TFH_genes = gene_list['Genes upregulated in TFH cells'].dropna().tolist()
TFH_genes = gene_list['list1'].dropna().tolist()
feature_genes = adata.var_names.intersection(TFH_genes)
feature_genes
data = np.sum(adata[:, feature_genes].X, axis=1).A1 / len(feature_genes)
data = np.interp(data, (data.min(), data.max()), (0, 10))
adata.obs['Tfh_score_1'] = data
#TFH_genes = gene_list['Genes upregulated in TFH cells'].dropna().tolist()
TFH_genes = gene_list['list2'].dropna().tolist()
feature_genes = adata.var_names.intersection(TFH_genes)
feature_genes
data = np.sum(adata[:, feature_genes].X, axis=1).A1 / len(feature_genes)
data = np.interp(data, (data.min(), data.max()), (0, 10))
adata.obs['Tfh_score_2'] = data
```

In [41]:

```
sc.pl.violin(adata,keys='Tfh_score_1',groupby='leiden',rotation=90)
```

```
... storing 'Top_expanded_clone' as categorical
```

In [42]:

```
sc.pl.violin(adata,keys='Tfh_score_2',groupby='leiden',rotation=90)
```

In [43]:

```
sc.pl.umap(adata, color='time')
```

In [44]:

```
adata.write(out_path+'/Merged_pre_all_cells.h5ad')
```

In [ ]:
