## Supplementary material for "Early acquisition of S-specific Tfh clonotypes after SARS-CoV-2 vaccination is associated with the longevity of anti-S antibodies": Scanpy_functions_v03262021.html


### scRNASeq Downstream analysis pipeline functions¶

- This is supposed to be the bench for the callable functions to be used in the future analysis

In [1]:

```
import scvelo as scv
scv.logging.print_version()
import warnings
import scirpy as ir
import scanpy as sc
import numpy as np
import scipy as sp
import pandas as pd
import matplotlib.pyplot as plt
from matplotlib import rcParams
from matplotlib import colors
import seaborn as sb
import bbknn
import logging
from sklearn.mixture import GaussianMixture
from scipy.stats     import norm
import glob
import os

from scipy import sparse
import scanpy.external as sce

import scanorama
```

```
Running scvelo 0.2.5 (python 3.9.16) on 2023-07-27 09:50.
```

```
ERROR: XMLRPC request failed [code: -32500]
RuntimeError: PyPI no longer supports 'pip search' (or XML-RPC search). Please use https://pypi.org/search (via a browser) instead. See https://warehouse.pypa.io/api-reference/xml-rpc.html#deprecated-methods for more information.
2023-07-27 09:50:54.408366: I tensorflow/core/platform/cpu_feature_guard.cc:193] This TensorFlow binary is optimized with oneAPI Deep Neural Network Library (oneDNN) to use the following CPU instructions in performance-critical operations:  SSE4.1 SSE4.2 AVX AVX2 AVX512F AVX512_VNNI FMA
To enable them in other operations, rebuild TensorFlow with the appropriate compiler flags.
2023-07-27 09:50:54.547844: I tensorflow/core/util/port.cc:104] oneDNN custom operations are on. You may see slightly different numerical results due to floating-point round-off errors from different computation orders. To turn them off, set the environment variable `TF_ENABLE_ONEDNN_OPTS=0`.
```

In [2]:

```
'''
import hvplot.pandas
import docx
from docx import Document
from docx.shared import Inches
from docx.shared import Pt
import holoviews as hv
import panel as pn
import bokeh
from bokeh.resources import INLINE
import gseapy
'''
```

Out[2]:

```
'\nimport hvplot.pandas\nimport docx\nfrom docx import Document\nfrom docx.shared import Inches\nfrom docx.shared import Pt\nimport holoviews as hv\nimport panel as pn\nimport bokeh\nfrom bokeh.resources import INLINE\nimport gseapy\n'
```

### Data IO and Integration¶

#### New version of the I/O takes¶

- Input pathway
- A dictionary of Samples
- Platform of the scRNASeq used, current default 10x

#### Returns¶

- A list of adata file with integration of potentially:
  1. TCR/BCR files
  2. Velocity loom files

In [3]:

```
#File I/O
def data_IO(samples,In_path,Platform = '10x_gex', IG = True,Velo=True):
    adatalist = []
    for sample, sample_meta in samples.items():
        gex_file = In_path+sample+'/outs/filtered_feature_bc_matrix.h5'
        if  Velo:
            loom_file = In_path+sample+'/velocyto/'+sample+'.loom'
            ldata = scv.read(loom_file, cache=True)
        if Platform == '10x_gex':
            adata = sc.read_10x_h5(gex_file, gex_only=False)
        
        if IG:
            tcr_file = In_path+sample_meta["TCR"]+'/outs/filtered_contig_annotations.csv'
            bcr_file = In_path+sample_meta["BCR"]+'/outs/filtered_contig_annotations.csv'
            adata_tcr = ir.io.read_10x_vdj(tcr_file)
            adata_bcr = ir.io.read_10x_vdj(bcr_file)
            ir.pp.merge_with_ir(adata, adata_bcr)
            adata.obs["is_cell"] = "None"
            adata_tcr.obs["is_cell"] = "None"
            adata.obs["high_confidence"] = "None"
            adata_tcr.obs["high_confidence"] = "None"
            ir.pp.merge_airr_chains(adata, adata_tcr)

        if  Velo:
            scv.utils.clean_obs_names(adata,id_length=16)
            scv.utils.clean_obs_names(ldata,id_length=16)
            adata = scv.utils.merge(adata, ldata)
        adata.var_names_make_unique()
        adata.obs["Sample"] = sample
        
        adatalist.append(adata)
    return adatalist
```

In [4]:

```
def unify_value(adatalist):
    uniset = set(adatalist[0].var_names)
    for adata in adatalist:
        uniset = uniset.intersection(set(adata.var_names))
        sample_tag = adata.obs["Sample"][0] 
        
    x = [g for g in uniset]
    gex = adatalist[0].concatenate(*adatalist[1:], join='outer')
    gex = gex[:,x]
    return gex
```

### Sparse and Dense matrix conversion¶

### Binarization of properities¶

In [5]:

```
def Thrshold_by_Gaussian_with_plot(adata, tag, out_path,  sample_tag, sparse = True):
    if tag in adata.obs.columns:
        data = adata.obs[tag].to_numpy().reshape(len(adata.obs[tag]),1)
    elif (sparse):
        data = adata[:, tag].X.A
        data =  np.interp(data, (data.min(), data.max()), (0, 10))
    else:
        data = adata[:, tag].X
        data =  np.interp(data, (data.min(), data.max()), (0, 10))
    
    N = len(data)
    
   
    x = np.linspace(0, 10, N+1)
    gmm = GaussianMixture(n_components=2, covariance_type='spherical')
    gmm.fit(data)
    mu1 = gmm.means_[0, 0]
    mu2 = gmm.means_[1, 0]

    var1, var2 = gmm.covariances_
    wgt1, wgt2 = gmm.weights_
    cut_off = (mu2 + mu1)/2
    delta  = abs(mu2 - mu1)
    var = (var1+var2)/2
    #data[data == 0] = np.nan
    outfile =out_path+'/figures/'+sample_tag+'_'+tag +'_Gaussian_Mixture_Model.png'
    plt.hist(data[data>0.1],bins='auto')
    plt.axvline(mu1,  color = 'y',linestyle=":",label='Fitted Mean 1')
    plt.axvline(mu2,  color = 'g',linestyle=":",label='Fitted Mean 2')
    plt.axvline(cut_off,  color = 'r', label='Threshold ')
    plt.legend()
    plt.title(sample_tag+' '+tag + ' Gaussian Mixture Model')
    plt.savefig(outfile)
    plt.show()
    plt.close()
    up_bound = max([mu1,mu2])
    low_bound = min([mu1,mu2])
    return [delta, var, cut_off,up_bound,low_bound]
```

### Quality Control¶

#### Adding Doublet Identification procedure into the QC steps¶

- Start from scrublet

#### Additional feature percentage calculation¶

- Percentage of Ribosome RNA
- Percentage of Himoglobin

#### Can not keep all the cell and genes without filtration¶

- Check the effects of removing the cells or not is a different story

#### Label them through threshold determination¶

- High concentration of mitochrondria
- Highest read counts
- Hemoglobin contamination

In [6]:

```
def qc_and_preprocess_lo_thres(adata, out_path, feature_tag = {'mt':'MT-','ribo':("RPS","RPL"),'hb':("HB")},
                      multi_sample=False, organism = "hsapiens"): 
    adata.var_names_make_unique()
    sc.pl.highest_expr_genes(adata, n_top=20, save='_before_filter.png')
    mito_tag = feature_tag['mt']
    for feature in feature_tag.keys():
        feature_genes = adata.var_names.str.startswith(feature_tag[feature]) 
        tag = 'pct_counts_'+str(feature)
        adata.obs[tag] = np.sum(
        adata[:, feature_genes].X, axis=1).A1 / np.sum(adata.X, axis=1).A1
    scrub_result = sce.pp.scrublet(adata,copy=True,threshold=0.5)
    adata.obs = scrub_result.obs
# for each cell compute fraction of counts in mito genes vs. all genes
# the `.A1` is only necessary as X is sparse (to transform to a dense array after summing)
#Convert all matrix into sparse matrix
    
# add the total counts per cell as observations-annotation to adata
    adata.obs['n_counts'] = adata.X.sum(axis=1).A1
    adata.obs['n_genes'] = (adata.X > 0).sum(axis=1).A1
  
    sc.pp.calculate_qc_metrics(adata,  inplace=True)
    sb.jointplot(data=adata.obs,
        x="log1p_total_counts", y="log1p_n_genes_by_counts",  kind="hex"
    )
    plt.savefig(out_path+'/figures/QC_hex.png')
    plt.close()
    %matplotlib inline
    sc.pl.violin(adata, ['n_genes', 'n_counts', 'doublet_score','pct_counts_mt','pct_counts_ribo', 'pct_counts_hb'],
                 jitter=0.4, multi_panel=True, save= '_QC_of_entire_set')
    adata.obs['log_counts'] = np.log(adata.obs['n_counts'])
    #Quality control - plot QC metrics
    #Sample quality plots
    #Beginning of the subsection

    if multi_sample:
        t1 = sc.pl.violin(adata, 'n_counts', groupby='Sample', size=2, log=True, cut=0,save = '_QC_by_sample_1.png',rotation = 90)
        t2 = sc.pl.violin(adata, 'pct_counts_mt', groupby='Sample',save = '_QC_by_sample_2.png',rotation = 90)
        sc.pl.violin(adata, 'pct_counts_ribo', groupby='Sample',save = '_QC_by_sample_3.png',rotation = 90)
        sc.pl.violin(adata,'pct_counts_hb', groupby='Sample',save = '_QC_by_sample_4.png',rotation = 90)
        t3 = sc.pl.violin(adata, 'doublet_score', groupby='Sample',save = '_scrublet_sample.png',rotation = 90)
        t4 = sc.pl.scatter(adata, x='total_counts', y='pct_counts_mt', color="Sample")
 
    
    sc.pl.scatter(adata, 'n_counts', 'pct_counts_mt',save = '_count_to_mito_1.png')
    p1 = sc.pl.scatter(adata, 'n_counts', 'n_genes', color='pct_counts_mt',save ='_count_to_gene.png')
    #Scatter plot zoom into the n_counts within the threshold
    #p2 = sc.pl.scatter(adata[adata.obs['n_counts']<10000], 'n_counts', 'n_genes', color='percent_mito',save ='_count_to_gene_threshold.png')
    #Thresholding decision: counts
    p3 = sb.distplot(adata.obs['n_counts'], kde=False)
    
    
    
    #Distribution plots 
    plt.savefig(out_path+'/figures/read_count_distribution.png')
    plt.show()
    plt.close()
#Thresholding decision: genes
    p6 = sb.distplot(adata.obs['n_genes'], kde=False, bins=100)
    plt.savefig(out_path+'/figures/n_genes_distribution.png')
    plt.show()
    plt.close()
 # Filter cells according to identified QC thresholds:
    delta, var, cut_off,up_bound,low_bound= Thrshold_by_Gaussian_with_plot(adata, 'pct_counts_mt', out_path, 'All_samples', sparse = False)
    mito_cut = cut_off
    
    delta, var, cut_off,up_bound,low_bound= Thrshold_by_Gaussian_with_plot(adata, 'n_counts', out_path, 'All_samples', sparse = False)
    count_up = int(up_bound+var)
    count_low = int(low_bound/16)
    
    delta, var, cut_off,up_bound,low_bound= Thrshold_by_Gaussian_with_plot(adata, 'n_genes', out_path, 'All_samples', sparse = False)
    gene_up = int(up_bound+delta)
    gene_low = int(low_bound/4)
    
    
    

    
    origin = adata.n_obs
    
    sc.pp.filter_cells(adata, min_counts = count_low)
    c1 =  origin - adata.n_obs 
    origin = adata.n_obs
    print('filtered out {:d} cells that have less than {:d} counts'.format(c1,count_low))
    ogene= adata.n_vars
    sc.pp.filter_genes(adata, min_cells=3)
    g1 = ogene - adata.n_vars
    ogene = adata.n_vars
    print('filtered out {:d} genes that are detected in less than 3 cells'.format(g1))
    

    sc.pp.filter_cells(adata, max_counts = count_up)
    c2 = origin - adata.n_obs  
    origin = adata.n_obs
    print('filtered out {:d} cells that have more than {:d} counts'.format(c2,count_up))
    mito_cut=0.2
    adata = adata[adata.obs['pct_counts_mt'] <  mito_cut]
    c3 = origin - adata.n_obs  
    origin = adata.n_obs
    print('filtered out {:d} cells that has over {:d}% reads belong to mitochondrial genes'.format(c3,int(mito_cut*100)))

   # adata = adata[adata.obs['pct_counts_ribo'] > 0.05, :]
   # cx = origin - adata.n_obs
   # origin = adata.n_obs
   # print('filtered out {:d} cells that has less than 5% reads belong to ribosome genes'.format(cx))
    #Change the threshold for Togashi dataset to avoid filter out 8k cells ~James
    sc.pp.filter_cells(adata, min_genes = gene_low)
    c4 =  origin - adata.n_obs  
    origin = adata.n_obs
    print('filtered out {:d} cells that have less than {:d} genes expressed'.format(c4,gene_low))
    
    adata = adata[adata.obs['n_genes'] < gene_up, :]
    c5 =  origin - adata.n_obs  
    origin = adata.n_obs
    
    print('filtered out {:d} cells that have over {:d} genes expressed'.format(c5,gene_up))
    sc.pl.violin(adata, ['n_genes', 'n_counts', 'doublet_score','pct_counts_mt','pct_counts_ribo', 'pct_counts_hb'],
                 jitter=0.4, multi_panel=True,save= '_QC_of_entire_set_after_filtration')
    
    return adata
```

In [7]:

```
def qc_and_preprocess(adata, out_path, feature_tag = {'mt':'MT-','ribo':("RPS","RPL"),'hb':("HB")},
                      multi_sample=False, organism = "hsapiens"): 
    adata.var_names_make_unique()
    sc.pl.highest_expr_genes(adata, n_top=20, save='_before_filter.png')
    mito_tag = feature_tag['mt']
    for feature in feature_tag.keys():
        feature_genes = adata.var_names.str.startswith(feature_tag[feature]) 
        tag = 'pct_counts_'+str(feature)
        adata.obs[tag] = np.sum(
        adata[:, feature_genes].X, axis=1).A1 / np.sum(adata.X, axis=1).A1
    scrub_result = sce.pp.scrublet(adata,copy=True,threshold=0.5)
    adata.obs = scrub_result.obs
# for each cell compute fraction of counts in mito genes vs. all genes
# the `.A1` is only necessary as X is sparse (to transform to a dense array after summing)
#Convert all matrix into sparse matrix
    
# add the total counts per cell as observations-annotation to adata
    adata.obs['n_counts'] = adata.X.sum(axis=1).A1
    adata.obs['n_genes'] = (adata.X > 0).sum(axis=1).A1
  
    sc.pp.calculate_qc_metrics(adata,  inplace=True)
    sb.jointplot(data=adata.obs,
        x="log1p_total_counts", y="log1p_n_genes_by_counts",  kind="hex"
    )
    plt.savefig(out_path+'/figures/QC_hex.png')
    plt.close()
    %matplotlib inline
    sc.pl.violin(adata, ['n_genes', 'n_counts', 'doublet_score','pct_counts_mt','pct_counts_ribo', 'pct_counts_hb'],
                 jitter=0.4, multi_panel=True, save= '_QC_of_entire_set')
    adata.obs['log_counts'] = np.log(adata.obs['n_counts'])
    #Quality control - plot QC metrics
    #Sample quality plots
    #Beginning of the subsection

    if multi_sample:
        t1 = sc.pl.violin(adata, 'n_counts', groupby='Sample', size=2, log=True, cut=0,save = '_QC_by_sample_1.png',rotation = 90)
        t2 = sc.pl.violin(adata, 'pct_counts_mt', groupby='Sample',save = '_QC_by_sample_2.png',rotation = 90)
        sc.pl.violin(adata, 'pct_counts_ribo', groupby='Sample',save = '_QC_by_sample_3.png',rotation = 90)
        sc.pl.violin(adata,'pct_counts_hb', groupby='Sample',save = '_QC_by_sample_4.png',rotation = 90)
        t3 = sc.pl.violin(adata, 'doublet_score', groupby='Sample',save = '_scrublet_sample.png',rotation = 90)
        t4 = sc.pl.scatter(adata, x='total_counts', y='pct_counts_mt', color="Sample")
 
    
    sc.pl.scatter(adata, 'n_counts', 'pct_counts_mt',save = '_count_to_mito_1.png')
    p1 = sc.pl.scatter(adata, 'n_counts', 'n_genes', color='pct_counts_mt',save ='_count_to_gene.png')
    #Scatter plot zoom into the n_counts within the threshold
    #p2 = sc.pl.scatter(adata[adata.obs['n_counts']<10000], 'n_counts', 'n_genes', color='percent_mito',save ='_count_to_gene_threshold.png')
    #Thresholding decision: counts
    p3 = sb.distplot(adata.obs['n_counts'], kde=False)
    
    
    
    #Distribution plots 
    plt.savefig(out_path+'/figures/read_count_distribution.png')
    plt.show()
    plt.close()
#Thresholding decision: genes
    p6 = sb.distplot(adata.obs['n_genes'], kde=False, bins=100)
    plt.savefig(out_path+'/figures/n_genes_distribution.png')
    plt.show()
    plt.close()
 # Filter cells according to identified QC thresholds:
    delta, var, cut_off,up_bound,low_bound= Thrshold_by_Gaussian_with_plot(adata, 'pct_counts_mt', out_path, 'All_samples', sparse = False)
    mito_cut = cut_off
    
    delta, var, cut_off,up_bound,low_bound= Thrshold_by_Gaussian_with_plot(adata, 'n_counts', out_path, 'All_samples', sparse = False)
    count_up = int(up_bound+var)
    count_low = int(low_bound/16)
    
    delta, var, cut_off,up_bound,low_bound= Thrshold_by_Gaussian_with_plot(adata, 'n_genes', out_path, 'All_samples', sparse = False)
    gene_up = int(up_bound+delta)
    gene_low = int(low_bound/4)
    
    
    

    
    origin = adata.n_obs
    
    sc.pp.filter_cells(adata, min_counts = count_low)
    c1 =  origin - adata.n_obs 
    origin = adata.n_obs
    print('filtered out {:d} cells that have less than {:d} counts'.format(c1,count_low))
    ogene= adata.n_vars
    sc.pp.filter_genes(adata, min_cells=3)
    g1 = ogene - adata.n_vars
    ogene = adata.n_vars
    print('filtered out {:d} genes that are detected in less than 3 cells'.format(g1))
    

    sc.pp.filter_cells(adata, max_counts = count_up)
    c2 = origin - adata.n_obs  
    origin = adata.n_obs
    print('filtered out {:d} cells that have more than {:d} counts'.format(c2,count_up))

    adata = adata[adata.obs['pct_counts_mt'] <  mito_cut]
    c3 = origin - adata.n_obs  
    origin = adata.n_obs
    print('filtered out {:d} cells that has over {:d}% reads belong to mitochondrial genes'.format(c3,int(mito_cut*100)))

   # adata = adata[adata.obs['pct_counts_ribo'] > 0.05, :]
   # cx = origin - adata.n_obs
   # origin = adata.n_obs
   # print('filtered out {:d} cells that has less than 5% reads belong to ribosome genes'.format(cx))
    #Change the threshold for Togashi dataset to avoid filter out 8k cells ~James
    sc.pp.filter_cells(adata, min_genes = gene_low)
    c4 =  origin - adata.n_obs  
    origin = adata.n_obs
    print('filtered out {:d} cells that have less than {:d} genes expressed'.format(c4,gene_low))
    
    adata = adata[adata.obs['n_genes'] < gene_up, :]
    c5 =  origin - adata.n_obs  
    origin = adata.n_obs
    
    print('filtered out {:d} cells that have over {:d} genes expressed'.format(c5,gene_up))
    sc.pl.violin(adata, ['n_genes', 'n_counts', 'doublet_score','pct_counts_mt','pct_counts_ribo', 'pct_counts_hb'],
                 jitter=0.4, multi_panel=True,save= '_QC_of_entire_set_after_filtration')
    
    return adata
```

### Normalization and Feature Selection¶

- Should add the process of imputation here

In [8]:

```
# Normalization and preprocessing
def feature_selection(adata, out_path, filter_genes, exclude = True):
    sc.pp.normalize_total(adata, inplace=True)
    #sc.pp.normalize_per_cell(adata, counts_per_cell_after=1e6)
  
    sc.pl.highest_expr_genes(adata, n_top=20, save='_after_QC.png')
    if exclude:
        kept =  ~adata.var_names.str.startswith('RPL')
    
        for fit in filter_genes:
            fi_genes = ~adata.var_names.str.startswith(fit)
            kept = kept & fi_genes
        filtered = ~kept
        filtered_genes =  adata[:, filtered].var_names
 
        adata = adata[:, kept]
    
        sc.pl.highest_expr_genes(adata, n_top=20, save='_after_exclude.png')
    adata.raw = adata
    sc.pp.log1p(adata)
   
    sc.pp.highly_variable_genes(adata, flavor='cell_ranger', n_top_genes=4000)
    print('\n','Number of highly variable genes: {:d}'.format(np.sum(adata.var['highly_variable'])))
    sc.pl.highly_variable_genes(adata, save='_highly_variable_genes.png')
    sc.pp.scale(adata,  zero_center=False)
    return adata
```

### Batch Correction¶

#### Scanorama Rocks!¶

### Clustering and Dimentional Reduction¶

In [9]:

```
def clustering(adata,samples,resol=1,multi = True): 

   # bbknn.bbknn(adata,batch_key='Sample')
    if multi:
        adatas = [adata[adata.obs['Sample']== x] for x in samples.keys()]
        corrected = scanorama.correct_scanpy(adatas, return_dimred=True)
        del adatas
        corrected_X = corrected[0].concatenate(*corrected[1:], join='outer')
        del corrected
        adata.obsm["X_scanorama"] = corrected_X.obsm["X_scanorama"]
        del corrected_X
        sc.pp.neighbors(adata, use_rep="X_scanorama")
    
        #sce.pp.scanorama_integrate(adata, 'Sample',batch_size=sample_counts)/
        #sc.pp.neighbors(adata, use_rep='X_scanorama')
    else:
        sc.pp.pca(adata, n_comps=50, use_highly_variable=True, zero_center=False, svd_solver='arpack')
        sc.pp.neighbors(adata)
    
    sc.tl.leiden(adata,resolution=resol)
    #sc.tl.louvain(adata,resolution=2)
    sc.tl.paga(adata, groups='leiden')
    #sc.tl.tsne(adata, n_jobs=40) #Note n_jobs works for MulticoreTSNE, but not regular implementation)
    #sc.tl.diffmap(adata)
    #sc.tl.draw_graph(adata)
    sc.pl.paga(adata,node_size_power= 1.0,threshold=0.1 ,edge_width_scale = 0.1,save ='_Graph.png') 
 
    # remove `plot=False` if you want to see the coarse-grained graph
    sc.tl.umap(adata,  init_pos=sc.tl._utils.get_init_pos_from_paga(adata), maxiter=100,min_dist=0.1, spread=4.0)
    
    sc.pl.umap(adata, color='leiden', legend_loc='on data',legend_fontsize=10,save ='_Leiden_clustering.png') 
    if multi:
        sc.pl.umap(adata, color='Sample')
        adata.obs.groupby(['leiden'])['Sample'].value_counts(normalize = True).unstack().plot.bar(stacked=True)
        ax = plt.subplot(111)
        plt.ylabel('Composition of cells')
        plt.xlabel('Leiden cluster')
        box = ax.get_position()
        ax.set_position([box.x0, box.y0, box.width * 0.8, box.height])
        ax.legend(loc='center left', bbox_to_anchor=(1, 0.5))

        
        adata.obs.groupby(['leiden'])['Sample'].value_counts().unstack().plot.bar(stacked=True)
        plt.ylabel('Amount of cells')
        plt.xlabel('Leiden cluster')
       
    return adata
```

In [10]:

```
# Batch Correction

## Another function written to merge the samples with BBKNN

# Clustering and Dimentional Reduction
```

In [11]:

```
def BBKNN_clustering(adata,out_path, multi_sample, resol=2): 
    
    # Calculate the visualizations
    sc.pp.pca(adata, n_comps=50, use_highly_variable=True, svd_solver='arpack')
    #sc.tl.pca(adata, svd_solver='arpack')
    
    if multi_sample:
        No_combat = adata.copy()
        sc.pp.neighbors(No_combat)
        sc.tl.umap(No_combat)
        bbknn.bbknn(adata,batch_key='Sample')
    else:
        sc.pp.neighbors(adata, n_neighbors=10, n_pcs=50)
    sc.tl.leiden(adata,resolution=resol)
    #sc.tl.louvain(adata,resolution=2)
    sc.tl.paga(adata, groups='leiden')
    #sc.tl.tsne(adata, n_jobs=40) #Note n_jobs works for MulticoreTSNE, but not regular implementation)
    #sc.tl.diffmap(adata)
    #sc.tl.draw_graph(adata)
    sc.pl.paga(adata,node_size_power= 1.0,threshold=0.1 ,layout ='rt',edge_width_scale = 0.1,save ='_Graph.png') 
  
    
    # remove `plot=False` if you want to see the coarse-grained graph
    sc.tl.umap(adata,  init_pos=sc.tl._utils.get_init_pos_from_paga(adata), maxiter=100, min_dist=0.1, spread= 4.0)
    
    sc.pl.umap(adata, color='leiden', legend_loc='on data',legend_fontsize=10,save ='_Leiden_clustering.png') 
  
    
    if multi_sample:
        sc.pl.umap(No_combat, color='Sample',save ='_no_bbknn.png')
        sc.pl.umap(adata, color='Sample',save ='_by_sample.png')
        adata.obs.groupby(['leiden'])['Sample'].value_counts(normalize = True).unstack().plot.bar(stacked=True)
        ax = plt.subplot(111)
        plt.ylabel('Composition of cells')
        plt.xlabel('Leiden cluster')
        box = ax.get_position()
        ax.set_position([box.x0, box.y0, box.width * 0.8, box.height])
        ax.legend(loc='center left', bbox_to_anchor=(1, 0.5))

        adata.obs.groupby(['leiden'])['Sample'].value_counts().unstack().plot.bar(stacked=True)
        plt.ylabel('Amount of cells')
        plt.xlabel('Leiden cluster')

    return adata
```

### Spatial Integration¶

In [ ]:

### Differential Expression¶

In [12]:

```
def differential(adata,out_path, multi_sample, top_n= 2,organism="hsapiens"):
    sc.tl.rank_genes_groups(adata, "leiden",use_raw=False, n_genes=adata.var_names.size)
    sc.pl.rank_genes_groups(adata, key='rank_genes_groups', save = '_by_cluster.png')

    rank_genes = pd.DataFrame(adata.uns['rank_genes_groups']['names'])
    top_2 = list(set(sum(rank_genes.head(top_n).values.tolist(),[])))
    rank_pavlues = pd.DataFrame(adata.uns['rank_genes_groups']['pvals'])
    rank_pavlues.rename(columns=lambda x: x+'_pvals', inplace=True)
    rank_fold = pd.DataFrame(adata.uns['rank_genes_groups']['logfoldchanges'])
    rank_fold.rename(columns=lambda x: x+'_logfoldchanges', inplace=True)
    pd.concat([rank_genes, rank_pavlues,rank_fold], axis=1, join='inner').to_csv(out_path+'/DE_genes_in_clusters.csv')

    sc.pl.umap(adata,color=top_2,color_map= 'OrRd',save ='_top2_in_clusters.png')
    sc.pl.rank_genes_groups_dotplot(adata,use_raw=False,save ='in_clusters.png')
    sc.pl.rank_genes_groups_heatmap(adata,use_raw=False,standard_scale='var',dendrogram=True,save ='_in_clusters.png')
    sc.pl.rank_genes_groups_matrixplot(adata,save ='in_clusters.png')
    sc.pl.rank_genes_groups_stacked_violin(adata,save ='in_clusters.png')
    sc.pl.rank_genes_groups_tracksplot(adata,n_genes = 2,save ='_in_clusters.png')
    clusters = adata.obs['leiden'].unique().tolist()

    for c in clusters:
        q= rank_genes[c].head(200).tolist()
        sc.queries.enrich(q, org=organism,gprofiler_kwargs={'no_evidences':False}).to_csv(out_path+'/cluster_'+c+'_within_cluster_enrichment.csv')
    
    if multi_sample:
        sc.tl.rank_genes_groups(adata, "Sample",use_raw=False,key_added='rank_genes_samples', n_genes=adata.var_names.size)
        sc.pl.rank_genes_groups(adata, key='rank_genes_samples', save = '_compare.png')
 
    

        rank_genes = pd.DataFrame(adata.uns['rank_genes_samples']['names'])
        top_3 = list(set(sum(rank_genes.head(top_n).values.tolist(),[])))
        rank_pavlues = pd.DataFrame(adata.uns['rank_genes_samples']['pvals'])
        rank_pavlues.rename(columns=lambda x: x+'_pvals', inplace=True)
        rank_fold = pd.DataFrame(adata.uns['rank_genes_samples']['logfoldchanges'])
        rank_fold.rename(columns=lambda x: x+'_logfoldchanges', inplace=True)
        pd.concat([rank_genes, rank_pavlues,rank_fold], axis=1, join='inner').to_csv(out_path+'/DE_genes_in_samples.csv')

    return adata
```

In [13]:

```
# Enrichment
```

In [14]:

```
def Binarize_Anti(adata,antilist,out_path,sample_tag):
    for a in antilist:
        thres = Thrshold_by_Gaussian_with_plot(adata, a, out_path,  sample_tag,sparse = False)
        data = adata.obs[a]
        adata.obs[a+'_bin'] = np.where(data > thres[2], a, 'NAN')
    return adata
```

In [15]:

```
# Cell Typing

# Velocity
# TCR/BCR
# Pseudo time
# Gene Regulatory Networks
# Perturbation
# Cell-Cell Communication
# TCR/BCR
# Ingestion
# Gene/Condition corrrelation

# Docx Report Generation
# Cell Browser 
# Cirrocumulus
# HTML Dashborad
```

In [ ]:
